# A histidine-mediated, pendulum-like proton transport mechanism is required for the high catalytic activity of [FeFe]-hydrogenases

**DOI:** 10.64898/2026.09.22.753672

**Authors:** Jifu Duan, Zhaoya Yang, Oliver Lampret, Federica Arrigoni, Lukas van Impel, Feng Zheng, Julian Kleinhaus, Ulf-Peter Apfel, Claudio Greco, Luca De Gioia, Eckhard Hofmann, Anja Hemschemeier, Thomas Happe

## Abstract

[FeFe]-hydrogenases are molecular hydrogen (H_2_) converting enzymes that employ a hexanuclear iron-complex, the H-cluster, as catalytic cofactor, and a proton transfer pathway (PTP) that allows efficient proton coupled electron transfer (PCET). Recent phylogenetic analyses revealed different groups of [FeFe]-hydrogenases. Although only very few members from other groups have been characterized, current knowledge suggests high catalytic activity is predominantly associated with group A members. Here, we show that metal ions inhibit group A [FeFe]-hydrogenases by binding to three specific residues at the entrance of the PTP. Exchanging residue H565 of *Clostridium pasteurianum* CpI results in a metal-insensitive protein variant with wildtype like activity. In contrast, exchanging S320 and H569 in CpI, and their counterparts in additional group A [FeFe]-hydrogenases results in enzymes with strongly decreased activities and large overpotential requirements. These features are consistent with important roles of both residues in catalytic proton transfer. CpI structures reveal that H569, locally anchored by E278, can be present in two conformations so that we propose a histidine-dependent pendulum mechanism for exchanging protons between bulk solvent and the PTP, which is well supported by theoretical calculations. By that, our study widens information on how the mobility of histidine, governed by residues in the vicinity, contributes to the fundamental concept of PCET.

## Introduction

[FeFe]-hydrogenases are metalloenzymes that catalyze interconversions of protons, electrons and molecular hydrogen (H_2_) with turnover frequencies of up to 10,000 s^-1 1^. This extraordinary catalytic efficiency is enabled by the active center, the so-called H-cluster and a well-tuned protein environment. The H-cluster consists of a canonical [4Fe-4S] cluster ([4Fe]_H_) and a uniquely structured diiron site ([2Fe]_H_) which are connected through a thiolate ligand of a cysteine residue from the protein scaffold (Figure 1)^2-3^. In addition to anchoring [2Fe]_H_ within the hydrophobic protein environment, the protein scaffold determines the catalytic properties by tuning the redox potential of [2Fe]_H_ and building substrate/product transfer chains^4^. The protein environment is also involved in determining the sensitivity of [FeFe]-hydrogenases to inhibitors such as carbon monoxide (CO)^5^, cyanide (CN^−^)^6-7^, hydrogen sulfide (HS^−^)^8- 10^, chloride (Cl^−^)^11^, formaldehyde^12-14^ and oxygen (O_2_)^15-19^, which mostly directly interact with [2Fe]_H_.

**Figure 1.**
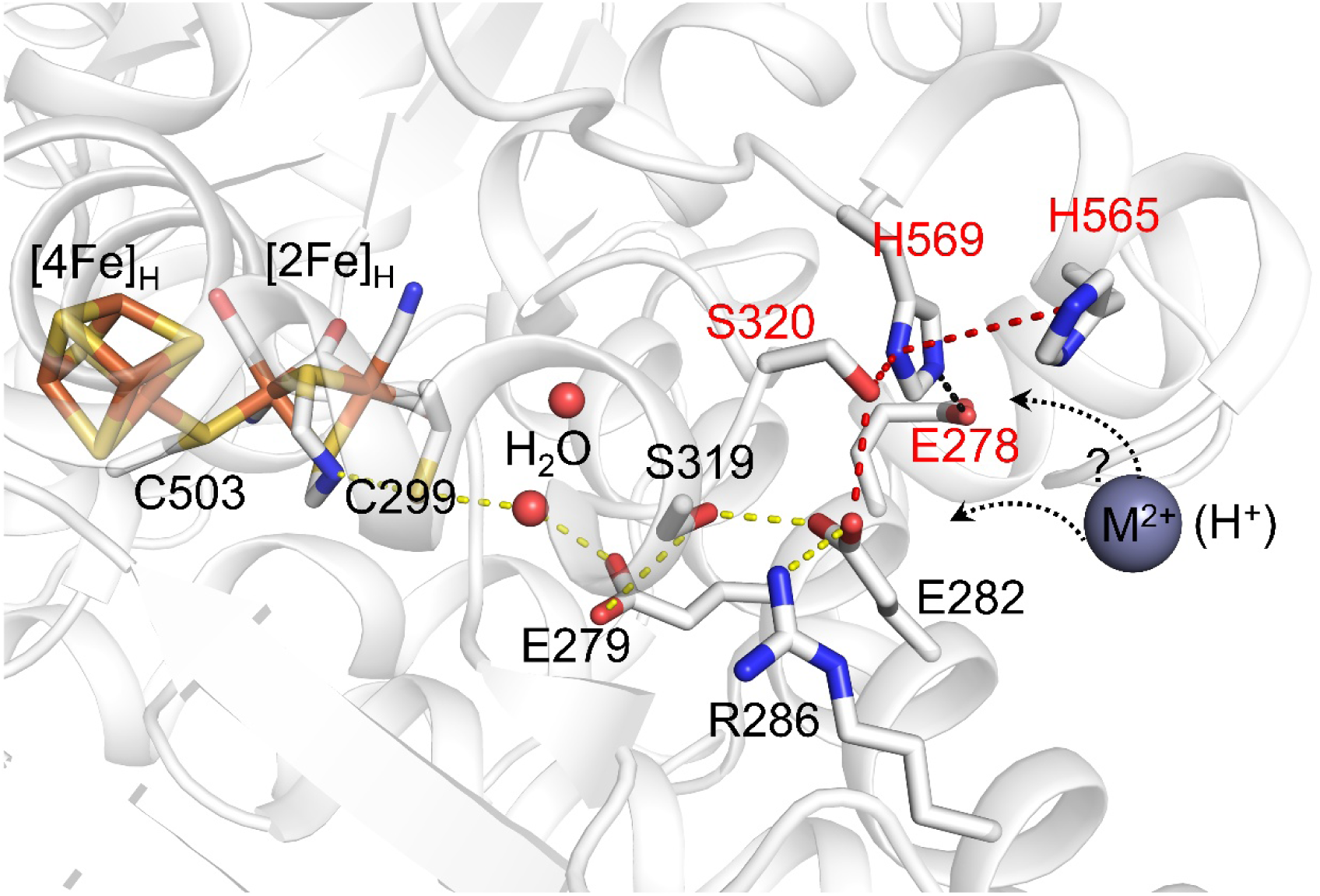
Active site and proton transfer pathway (PTP) of [FeFe]-hydrogenase CpI. The H-bond network of the PTP, indicated by yellow and dashed lines, connects the H-cluster to the protein surface. M^2+^ represents divalent metal ions that could bind to the surface residues of the PTP^6,30^. The arrows at the entrance of the PTP indicate possible binding sites of metal ions and entry points of catalytic protons. Residues S320, H569, E278, and H565 that we identified here as extensions of the previously defined PTP are labeled red. E278 is not directly involved in proton transfer, but important in anchoring H569, see details below. The figure is based on the CpI crystal structure with protein data bank (PDB) accession number 4XDC^39^.

Recent phylogenetic analysis of [FeFe]-hydrogenase protein sequences has substantially expanded our understanding of their biodiversity. Today, [FeFe]-hydrogenases are classified into seven groups (A-G). Group A is comprised of prototypical and electron-bifurcating [FeFe]-hydrogenases^20^. According to current knowledge, prototypical [FeFe]-hydrogenases exhibit the highest H_2_ production and/or oxidation activities. Several well-characterized examples that belong to this group are CpI^2^, DdH^3^, CbA5H^21^ and HydA1^22^ from *Clostridium pasteurianum, Desulfovibrio desulfuricans, Clostridium beijerinckii* and *Chlamydomonas reinhardtii*, respectively. All but a few of the enzymes from other groups that have been characterized so far showed much lower activities than these well-characterized prototypical candidates. Variations in residues surrounding [2Fe]_H_ and forming the proton transfer pathway (PTP) are probably responsible for the large differences in catalytic activities.^23-29^

The PTP was initially identified by Cornish et al. through site directed mutagenesis studies on CpI.^30^ Subsequent work from our group and others provided evidence that this PTP is conserved in other group A [FeFe]-hydrogenases.^31-32^ Through electrochemistry, we could demonstrate that the transport of protons through the pathway is thermodynamically coupled with electron transfer in CpI and HydA1.^33^ The PTP is very conserved in [FeFe]-hydrogenases of groups A and B but not in groups C to G^20, 25, 29^. In CpI, from the active site to the protein surface, it has been defined to consist of C299, E279, S319 and E282 with two protein-bound water molecules between C299 and E279 (Figure 1). The surface-exposed side chain of E282 establishes an H-bond and a salt bridge with S320 and R286, respectively. Notably, the actual entry point of protons remains unresolved, and it has been suggested that either E282 directly exchanges protons with the bulk solvent or that protons enter through R286 or S320^30, 32, 34-36^ (Figure 1).

Proton entry at the interface of the bulk solvent and the protein surface has been analyzed employing Zn^2+^ and other divalent metal ions as probes. These ions coordinate to specific surface residues that form the entrances of PTPs in several vital enzyme systems such as respiratory cytochrome *c* oxidase^37^ or photosynthetic reaction centers^38^. In a previous study, the effect of Zn^2+^ on CpI was tested and shown to inhibit hydrogenase activity by about 90%.^30^ The authors suggested that Zn^2+^ exerts this inhibitory effect on CpI by binding to the important PTP residue E282. In this regard, it caught our attention that another divalent metal ion, namely magnesium (Mg^2+^) from the crystallization cocktail, coordinated to the side chains of S320 and a nearby histidine (H565) in some of the known CpI crystal structures (Figure S1)^6^. These two previous studies prompted us to employ crystal structure analyses to investigate how Zn^2+^ and additional divalent metal ions interact with [FeFe]-hydrogenases and whether metal ions may be used as probes to advance our mechanistic understanding of catalytic proton transfer in these H_2_-converting enzymes. Combining protein crystallography, activity assays, electrochemistry and site-directed mutagenesis, we reveal molecular details on how Zn^2+^, Fe^2+^ and Ni^2+^ inhibit [FeFe]-hydrogenases. We show that the surface-exposed H565 is involved in the coordination of all three of these metal ions, and exchanging this residue results in metal-resistant enzymes with otherwise only slightly affected catalytic activity. Notably, coordination of these metal ions with H565 also involves the side chains of H569 or S320 and variants of these residues show strongly reduced catalytic activities and significantly increased overpotentials. In CpI structures obtained at different pH values, H569 was found to be present in different conformations resembling a pendulum, mounted by an H-bond to E278. Density functional theory (DFT) calculations indicate that both these motions and anchoring of the histidine side chain are important for efficient proton entry. Our study demonstrates important roles of H569 and S320 in mediating catalytic proton transfer in [FeFe]-hydrogenases and thereby reveals new players of PCET, particularly at the surface-solvent interface.

## Results

### H565, H569 and S320 at the entrance of the PTP of CpI are the primary targets of metal ions

To gain structural insights into how metal ions bind to [FeFe]-hydrogenases, X-ray crystallography was applied to CpI. The wildtype form of CpI (termed CpI-WT hereafter) was crystallized as before^39^ and the resulting crystals were soaked with ZnCl_2_, FeSO_4_, and NiCl_2_ (the crystallization and soaking conditions are listed in Table S1). The structures of these soaked crystals were refined to resolutions of 1.45 to 1.65 Å and revealed high similarities to the structures obtained without metal ion soaking as indicated by the root mean square deviation values (Table S2).

The 1.65 Å structure obtained from Zn^2+^-soaked CpI crystals at pH 8 (referred to as CpI-Zn in the following) shows additional electron densities at seven sites on the surface including one site at the entrance of the PTP. The presence and absence of anomalous densities from two X-ray datasets of the same crystal collected at 9.665 keV and 9.63 keV allows for conclusive assignments of Zn^2+^ at these sites (Figures S2-S3, Table S3)^40^. The binding modes of Zn^2+^ on the surface of the CpI protein mostly show a tetrahedral coordination and involve residues C39, C184, E444, H500, H511, E535, H565 and H569 (Figure S2). In addition to coordinating water molecules, several Zn^2+^ binding sites also involve Cl^−^ ions that originate from purification and crystallization buffers (Figures S2-S3). Of particular interest to us was the binding of Zn^2+^ at the entrance of the PTP. This Zn^2+^ ion is coordinated by the side chains of H565 and H569 and two Cl^−^ ions (Figure 2b; Figures S2-S3) with refined coordination distances of about 2.2 Å. The identity of Cl^−^ ions was confirmed by merged anomalous densities from two X-ray datasets collected at X-ray energies of 6 keV and 6.2 keV (Figure 2 and Figure S3, Table S3). This coordination requires slight conformational changes of the side chains of H565 and H569 (Figure 2a and 2b).

**Figure 2.**
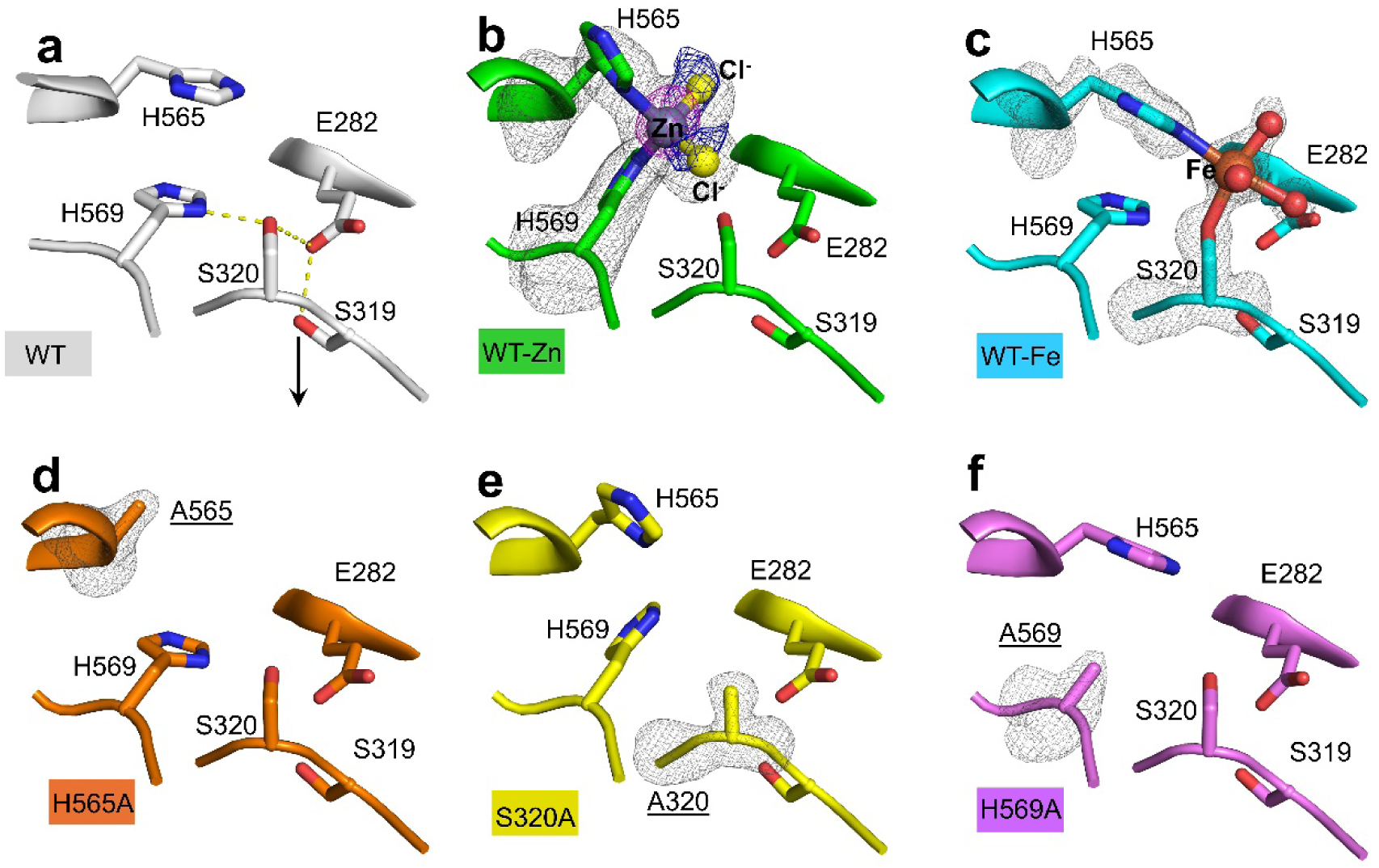
Configurations of the PTP entrance as observed in crystal structures of CpI-WT and CpI variants. The PTP entrances as observed in the crystal structures of CpI-WT (PDB: 4XDC^39^) (**a**), CpI-WT soaked with ZnCl2 (9S7W) (**b**), CpI-WT soaked with FeCl2 (9S9S) (**c**), variant H565A (9S9D) (**d**), variant S320A (9S9W) (**e**) and variant H569A (9S9V) (**f**) are shown, applying the same orientation for each structure. The structures are additionally distinguished by differently colored carbon atoms. In panel **a**, the black downwards arrow indicates the direction of the proton transfer towards the H-cluster. In panels **b** to **f**, the omit maps (gray mesh) were contoured at 3-4 σ. In panel **b**, the Zn^2+^-specific anomalous map obtained with X-ray data collected at 9.665 keV is colored pink and contoured at 8 σ. The anomalous map of the Cl^−^ ion (dark blue mesh) obtained with X-ray data collected at 6.2 and 6 keV was contoured at 3.5 σ.

The structure of CpI-WT soaked with Fe^2+^ at pH 6 (abbreviated as CpI-Fe here) could be resolved to a resolution of 1.5 Å and shows only one Fe^2+^-binding site (Table S4), which is at the entrance of the PTP (Figure 2c, Figure S4). The presence of Fe^2+^ was confirmed by the presence and absence of anomalous densities in two X-ray datasets of a second Fe^2+^-treated crystal (termed CpI-Fe’) collected at energies of 7.145 keV and 7.0 keV (Figure S5). The Fe^2+^ ion is octahedrally coordinated by the side chains of H565 and S320 together with four additional ligands. When modeling the ligands as water molecules, the temperature factors of the water molecules are very close to the temperature factors of the Fe^2+^ ion and its coordinated side chain of H565 and S320. When these ligands were modeled as chloride ions, their temperature factors were at least double those of the modeled water molecules, suggesting a much better fitting with water molecules. This coordination profile is very similar to the configuration we observed for the Mg^2+^ ion bound to CO- and CN^-^-treated CpI mentioned in the introduction (Figure S1)^6^.

In the 1.5 Å structure of CpI soaked with Ni^2+^ at pH 8 (referred to as CpI-Ni-1 here), the Ni^2+^ ion is octahedrally coordinated by the side chains of H565 and H569, three water molecules and a Cl^−^ ion at the entrance of the PTP (Figure S6). When the soaking buffer was at pH 6, the anomalous density of Ni^2+^ obtained from X-ray data collected at 8.35 keV revealed two Ni^2+^ sites with a distance of only about 2 Å at the PTP entrance, indicative of two different coordination conformations, which is consistent with two conformations (A and B) of the side chains of H565 and H569 that could be refined to occupancies of about 0.59 (conformation A) and 0.41 (conformation B) (Figures S6). In conformation B, the Ni^2+^ ion is coordinated by the side chains of H565 and H569, similar to the binding mode observed in CpI-Ni-1. In conformation A, the Ni^2+^ ion is coordinated by the side chains of H565 in the second conformation and of S320 (Figure S6). Note the coordinating partners of Ni^2+^ such as water molecules and Cl^−^ could not be modelled conclusively in CpI-Ni-2 because these two conformations overlap. The two Ni^2+^ sites were also confirmed by two data sets collected at X-ray energies of 8.35 keV and 8.335 keV from a second Ni^2+^-soaked CpI crystal at pH 6 (termed CpI-Ni-2’, Figure S7). In addition to the observed Ni^2+^ coordination sites at the PTP entrance, the structures of Ni^2+^-soaked CpI crystals showed several other Ni^2+^ binding sites on the surface, involving residues E52, E162, S320, E444, E535, H565 and H569 and some water molecules (Figure S8). Because we observed two different binding modes of Ni^2+^ at the entrance of the PTP depending on the pH of the soaking solution, we also performed the soaking experiments with Fe^2+^ and Zn^2+^ at the respective other pH environment. However, the binding modes of Fe^2+^ (shown here after crystal-soaking at pH 6) and Zn^2+^ (shown here after soaking the crystals at pH 8) at the entrance of the PTP remained unchanged in the respective different pH environment.

Summarizing the structural data obtained by soaking CpI-WT crystals with metals shows that although the binding modes of the metals at the PTP entrance differed moderately, all coordinate to the side chain of H565, and all might inhibit proton uptake as observed in other enzyme systems^38^ because they block the entry of the PTP. Therefore, we generated a CpI variant in which we exchanged H565 by alanine and tested the effect of the metal ions on the activities of both the wildtype enzyme and the H565A variant.

### Probing the inhibitory effects of metals in H2 production assays

Employing hydrogenase activity assays, H_2_ production activities of CpI were compared in the absence or presence of metals added individually to the reaction mixtures. The H_2_ evolving activity of CpI dropped by about 94% when 1 mM ZnCl_2_ was present in the assay at pH 8 (Figure 3a), while full activity was restored with the additional supplement of 1 mM ethylenediaminetetraacetic disodium (EDTA-2Na) salt, a well-known metal chelator (Figure S9). Because the Strep-tagII sequence is close to H569 and contains polar and potentially Zn^2+^-coordinating residues, including a histidine, we compared the inhibitory effect of ZnCl_2_ on CpI enzymes with and without Strep-tagII (see details on how the latter was obtained in the methods section). Non-tagged CpI was inhibited to the same extent as Strep-tagged CpI, which indicates that inhibitory effects of ZnCl_2_ were independent from the presence or absence of a Strep-tagII (Figure S9). Employing different concentrations of ZnCl_2_, a half-maximal inhibitory concentration (IC_50_) of about 0.03 mM was determined for CpI (Figure S10), which is lower than the Ki of 0.17 mM as reported by Cornish et al 2011^30^. The difference very likely results from the different pH conditions used in these two studies (see below). Nevertheless, both values are several orders of magnitude higher than the estimated labile pool of Zn^2+^ within cells.^41^ Thus, the inhibitory effects revealed in this study are very likely limited to in vitro conditions only.

**Figure 3.**
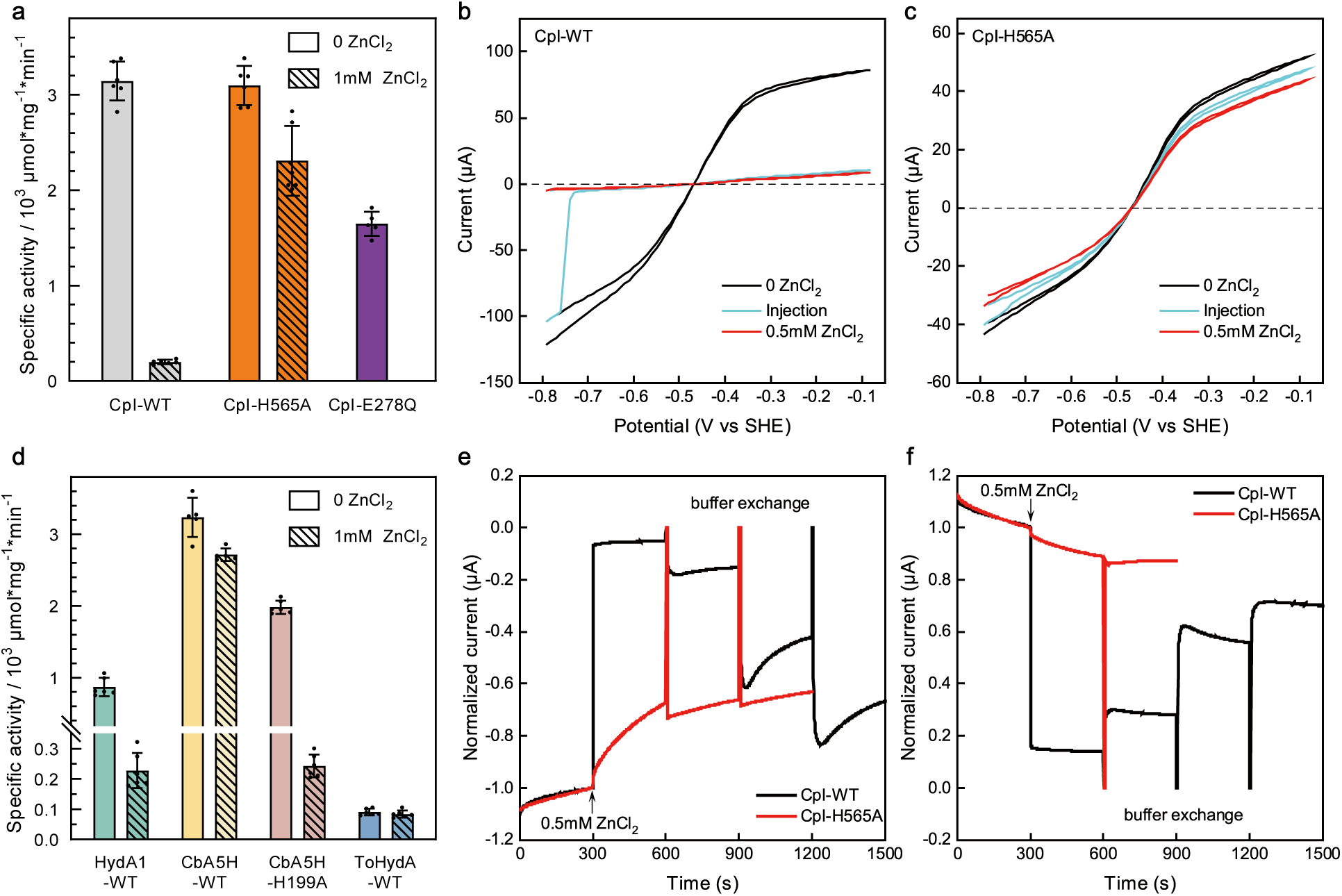
Kinetic assays of [FeFe]-hydrogenases in the absence and presence of ZnCl2. . Hydrogenase activity was tested in solution (**a**, **d**) or through protein film electrochemistry (PFE) (**b**, **c**, **e** and **f**). (**a** and **d**) The hydrogenase activity of the indicated [FeFe]-hydrogenases was tested in the absence (plain bars) or presence (striped bars) of 1 mM ZnCl2 in solution at pH 8. 100 mM sodium dithionite and 10 mM methyl viologen served as reducing reagent and electron mediator, respectively, and the reaction mixture additionally contained 500 mM NaCl. Samples were incubated upon shaking for 20 min at 37 °C before quantifying H2 in the headspace by gas chromatography. The bars show averages of two biological duplicates (referring to independent protein batches), determined in five to six technical duplicates, and the error bars indicate the standard deviation. (**b**-**c**) PFE was performed in the cyclic voltammetry mode, employing a mixed buffer system. Each one full scan was done before (black lines), during (cyan lines) and after (red lines) the injection of 0.5 mM ZnCl2. CV scans of CpI-WT and variant H565A are shown in (**b**) and (**c**), respectively. (**e**) and (**f**) show PFE experiments in the chronoamperometry mode, where the applied potential was –0.8 V to allow H2 production (**e**) or –0.1 V to induce H2 oxidation (**f**). In both (**e**) and (**f**), 0.5 mM ZnCl2 was injected after 300 s, as indicated by the arrows. After 600 s, 900 s and 1200 s, buffer exchange steps were performed. The currents were normalized to their values at 300 s. All PFE experiments were performed at pH 8 applying a working electrode rotation of 3000 rpm and a temperature of 10 °C. The flow of H2 gas was set to 30 L × min^-1^, except for the experiments shown in (**e**), where no H2 was applied. The scan rate in (**b**) and (**c**) was 10 mV × s^-1^. The experiments in (**b**, **c**, **e** and **f**) were conducted four times from two biological replicates with consistent results.

H_2_ production activities of variant H565A were tested under the same conditions as those of CpI-WT. Notably, the CpI-H565A variant exhibited the same level of H_2_ production activity as CpI-WT in the absence of ZnCl_2_, and its activity decreased only by about 25% in the presence of 1 mM ZnCl_2_ (Figure 3a). To analyze if this observation correlated with Zn^2+^ binding, crystal structures of the CpI-H565A variant were obtained without and with Zn^2+^ soaking at resolutions of 1.34 Å and 1.45 Å, respectively. Both structures showed an overall folding very similar to that of CpI-WT (Table S2). The diminished electron densities at the side chain of residue 565 confirmed the presence of an alanine instead of a histidine at this position (Figure 2d). Several of the Zn^2+^ binding sites on the surface of the CpI-H565 variant were the same as those observed for the Zn^2+^-treated CpI-WT protein, however, no significant densities resulting from Zn^2+^ could be identified at the entrance of the PTP and at position 565, respectively (Figure S11). This is a strong indicator that Zn^2+^ did not bind at this site where an alanine is now present instead of a histidine and suggested to us that the diminished inhibitory effect that Zn^2+^ had on the H_2_ evolution activity of the H565A variant might be related to the absence of this specific binding event. Infrared spectroscopy was employed to investigate whether the exchange of H565 or the addition of ZnCl_2_ would affect the H-cluster. However, we did not observe any noticeable changes in the H-clusters induced by the supplement of ZnCl_2_ for both WT and H565A enzymes (Figure S12). To corroborate the specificity of the Zn^2+^ inhibitory effect we generated additional six CpI variants in which residues involved in other Zn^2+^ coordination sites observed in the structure of CpI-Zn were exchanged by alanine (C39A, C184A, E444A, H500A, H511A and E535A). H_2_ production activities of these variants were tested in the absence or presence of ZnCl_2_ as had been done for the CpI-WT protein and variant H565A. All variants except H511A had hydrogenase activities very similar to the wildtype enzyme, and in all cases, the addition of ZnCl_2_ resulted in more than 80% inhibition of H_2_ production activities (Figure S9). These results indicate that while Zn^2+^ is readily coordinated to different sites of CpI, the metal ion inhibits H_2_ production mostly through binding to H565. As mentioned in the introduction, Mg^2+^ was found to coordinate to H565. The presence of MgCl_2_ in concentrations of up to 50 mM in the hydrogenase assay did not result in significant differences in H_2_ production activities of CpI (Figure S9), probably because Mg^2+^ usually does not form stable interactions with proteins^42^.

In view of the structural data we obtained by soaking CpI crystals with Fe^2+^ and Ni^2+^, we also attempted to test for inhibitory effects of these metal ions employing the same hydrogenase assays. However, we observed significant precipitation in the reaction tubes, suggesting that Fe^2+^ and Ni^2+^ - introduced by adding FeSO_4_ or NiCl_2_ - interact with reaction mixture components. This effect was mostly circumvented in protein film electrochemistry experiments that we applied to gain more detailed information about the kinetics of inhibitory effects by metal ions.

### Inhibitory effects of metal ions probed by protein film electrochemistry

In protein film electrochemistry (PFE), redox enzymes are immobilized on a suitable electrode, and currents that represent the enzymatic activity are monitored in dependence of an applied potential as driving force. PFE allows to analyze effects of additives on redox enzymes in kinetic resolution and at different redox potentials^12^. Here, we first used cyclic voltammetry (CV) during which the applied redox potential is continuously increased and decreased again, allowing to monitor the enzyme-generated currents as a function of potential. In the case of hydrogenases, negative and positive currents represent H_2_ production and H_2_ oxidation activities, respectively. We first tested whether the metals under investigation would have effects on the currents themselves and introduced ZnCl_2_, NiCl_2_ or FeSO_4_ as well as EDTA-2Na into the electrochemical cell in the absence of enzymes. None of the salts resulted in significant influences on the currents in the range of the potentials applied here, indicating that they did not specifically interact with the electrode (Figure S13).

When the CpI-WT enzyme was present on the electrode, introducing 0.5 mM ZnCl_2_ into the buffer at pH 8 caused a strong decrease of the currents (> 90%) over the applied potential range of –0.8 to –0.08 V vs SHE (standard hydrogen electrode; note that all potentials indicated in this study refer to SHE) (Figure 3b). In contrast, the current recorded for the CpI-H565A variant decreased by only about 15% when ZnCl_2_ was added (Figure 3c). The effect of ZnCl_2_ on the above-mentioned CpI exchange variants of residues involved in other Zn^2+^ coordination sites were also tested employing CV. The currents measured for each variant decreased by more than 85% after ZnCl_2_ had been injected into the electrochemical cell, very similar to what was observed in the case of CpI-WT (Figure S14).

To analyze the observed inhibitory effects of ZnCl_2_ during PFE experiments in a higher resolution and to allow for testing its reversibility without interfering current changes induced by a changing of redox potential, we employed chronoamperometry (CA). In CA, the redox potential is fixed at a certain potential that in the case of hydrogenases allows either H_2_ production or H_2_ uptake, and the currents are monitored in time. Under H_2_ production conditions (at a potential of –0.8 V), the current of the CpI-WT enzyme decreased slightly over the first 300 s when no additive was present. This is a typical observation in PFE experiments when the enzyme is not covalently coupled to the electrode, resulting in some film loss over time (Figure 3e). When 0.5 mM ZnCl_2_ was injected into the electrochemical cell after 300 s, an immediate decrease of the current by about 94% was observed. Exchanging the buffer by ZnCl_2_-free buffer three times (after 600, 900 and 1200 s) resulted in a recovery of the current to 16%, 60% and finally 83% of the current measured before introducing ZnCl_2_ (Figure 3e). Taking into account the film loss during the measurement, the current mostly recovered after three buffer exchanges, suggesting that the ZnCl_2_-mediated inhibition of CpI-WT under these conditions is almost fully reversible. Very similar results were obtained when the potential was set to –0.1 V to stimulate H_2_ uptake, namely a strong inhibition upon ZnCl_2_ injection and a stepwise recovery upon three buffer exchanges (Figure 3f). In this case, however, again accounting for film loss, the current reached its original trace, suggesting that the effect of Zn^2+^ is fully reversible in the direction of H_2_ oxidation (Figure 3f). We also tested whether the addition of EDTA-2Na would relieve the inhibitory effect of ZnCl_2_ on CpI-WT, and this was indeed the case as the currents in the directions of both H_2_ production and uptake were completely restored (Figure S15).

The effect of ZnCl_2_ on the CpI-H565A variant was also analyzed by CA experiments (Figure 3e and 3f). At a potential of –0.1 V that results in H_2_ oxidation, the injection of 0.5 mM ZnCl_2_ did not change the current significantly, indicating that ZnCl_2_ did not or hardly exert an inhibitory effect on the H565A variant under this condition (Figure 3f). Under H_2_ evolution conditions, induced by setting the potential to –0.8 V, the injection of 0.5 mM ZnCl_2_ resulted in a decrease of the current (Figure 3e). The decrease, however, was much slower than that observed for the wildtype CpI enzyme. The original current generated by the H565A dropped by only 33% over 300 s, whereas the inhibition by 94% observed for the CpI-WT enzyme was complete within 3 s. In contrast to the wildtype enzyme, two buffer exchanges did hardly result in any increase of the current of the ZnCl_2_-treated H565A variant (Figure 3e). This observation suggested that although the inhibitory effect of Zn^2+^ on the CpI-H565A variant appeared weaker, it was mostly irreversible in this PFE experiment. To test for this apparent irreversibility under different experimental conditions, we performed the dithionite-driven H_2_ production assay in solution, but preincubated both the wildtype CpI enzyme and the H565A variant with 1 mM of ZnCl_2_ for 5 and 60 min (Figure S16). Afterwards, the Zn^2+^-treated enzymes were transferred to the assay mixture, resulting in a 1000-fold dilution of the Zn^2+^ ions, and the hydrogenase activity assay was performed. This experiment mimics the buffer exchange step performed during the CA experiments, but on a much longer time scale. Both the Zn^2+^-preincubated CpI-WT enzyme and the H565A variant showed about the same hydrogenase activity as determined in the complete absence of ZnCl_2_, which suggests full reversibility of any inhibitory effect in the dithionite-driven assays in the case of both WT and H565A. The observation corroborates the results obtained with the CpI-WT protein during the CA experiments, but contrasts with the apparent irreversibility observed for the H565A variant (compare Figure 3e and Figure S16) which is probably due to a slow dissociation of Zn^2+^ from the latter.

PFE experiments were also carried out to investigate the inhibitory effects of 1 mM FeSO_4_ (Figure S17) and 1 mM NiCl_2_ (Figure S18) on the CpI-WT protein and variant H565A. Although the inhibitory effects of these metal ions were not that strong as those of ZnCl_2_, the results were similar to those obtained before in that the WT protein was reversibly inhibited by these metals under H_2_ production and uptake conditions, whereas the H565A variant appeared insensitive to the two metal salts as indicated by hardly changed currents after their addition (Figures S17 and S18).

### pH and Cl^−^-dependency of metal ions mediated inhibition

Summarizing the results we obtained with CpI-WT and H565A enzymes, it can be concluded that the analyzed metal ions inhibit CpI by binding to the side chains of the surface residues S320 or H569 together with H565, as presented in the structures (Figure 2). These binding events require the deprotonation of the side chains, which should be affected by pH. Therefore, we tested a pH effect on the inhibitory effects of Zn^2+^ and Ni^2+^. CV scans of CpI-WT in the absence and presence of Zn^2+^/Ni^2+^ were performed at pH 6 and the results were compared with those obtained at pH 8. Injection of 0.5 mM Zn^2+^ led to declines of currents by about 60-88% and 90-96% at pH 6 and pH 8, respectively (Figure S19). This suggested a stronger inhibition at pH 8, although the concentration of soluble Zn^2+^ at pH 8 should practically be even lower due to a decreased solubility at alkaline pH. In the dithionite-driven H_2_ production assays, activity of CpI was inhibited by 36% and 94% at pH 6 and 8, respectively (Figure S19). PFE experiments in the presence of Ni^2+^ also revealed a stronger inhibition at pH 8 than at pH 6 (Figure S18). These results suggest that the Zn^2+^ and Ni^2+^-mediated inhibition of CpI is indeed pH dependent.

Because the Zn^2+^ coordinated by H565 and H569 as observed in the CpI-Zn structure is also coordinated to two Cl^−^ ions (Figure 2b), we also tested whether the inhibitory effect of Zn^2+^ depends on the presence of chloride. For this purpose, we prepared CpI-WT samples in Cl^−^-free buffer and probed their sensitivity to ZnSO_4_ in a hydrogenase assay that was prepared by replacing NaCl by Na_2_SO_4_. The hydrogenase activity was very similar in both buffers, and the presence of 1 mM ZnSO_4_ in Cl^−^-free buffer resulted in a decrease of about 81% of the H_2_ production activity (Figure S19). This suggests that the Zn^2+^-mediated inhibition of CpI is mostly independent of the presence of Cl^−^ ions. As the CpI-Zn structure showed that some Zn^2+^ ions are coordinated by water molecules (Figure S2), it seems reasonable to suggest that water molecules can replace Cl^−^ ions to coordinate the Zn^2+^ ion at the entrance of the PTP.

### Zn^2+^ inhibits diverse [FeFe]-hydrogenases to various extents

To gain a broader view on the metal ion-mediated inhibition of [FeFe]-hydrogenases, we investigated the effect of Zn^2+^ on a selection of other [FeFe]-hydrogenases of different domain architecture and from different groups. We chose the small monomeric [FeFe]-hydrogenase from *C. reinhardtii*, HydA1, the O_2_-stable homodimeric CbA5H enzyme from *C. beijerinckii* (both group A [FeFe]-hydrogenases) and a group B [FeFe]-hydrogenase from *Thermosediminibacter oceani* (ToHydA)^29^. The residues forming the PTP and the two histidine residues that we could show to coordinate Zn^2+^ at the entrance of the PTP in CpI are structurally conserved in HydA1 and CbA5H^2, 18, 22^(Figure S20). Aligning the sequence of ToHydA^29^ to these three group A enzymes shows that the residues that form the PTP are conserved, as is the histidine that corresponds to CpI-H565, whereas H569 is not conserved (Figure S21). This information suggests similar inhibitory effects of Zn^2+^ on HydA1 and CbA5H as observed for CpI whereas ToHydA might be insensitive to Zn^2+^.

In the case of HydA1, we obtained a 2 Å structure of a Zn^2+^-soaked crystal of HydA1 without the diiron subsite of the H-cluster (termed HydA1-Δ[2Fe]_H_ here). It shows the presence of a Zn^2+^ ion at the entrance of the PTP that is coordinated by H478 and H482 (corresponding to H565 and H569 of CpI) and two Cl^−^ ions (Figures S22 and S23), in a very similar binding mode as that observed for CpI. The activity of the small algal [FeFe]-hydrogenase HydA1 was indeed about 74% lower when ZnCl_2_ was present in the hydrogenase assay (Figure 3d). The results suggest an overall very similar Zn^2+^-mediated inhibitory mechanism in HydA1 and CpI.

In the case of the homodimeric CbA5H enzyme, the presence of ZnCl_2_ in the hydrogenase assay resulted in a decrease of hydrogenase activity by only 16% (Figure 3d), which was surprising at first because of the high degree of conservation of the residues involved. An analysis of its published structure^18, 43^ with a focus on the entrance of the PTP revealed that the second subunit of the homodimer partially restricts solvent access to H630 and H634 (that correspond to CpI-H565 and H569), particularly for hydrated Zn²⁺ ions given their substantially larger size, in contrast to the monomeric CpI or HydA1 enzymes (Figure S24). To test the hypothesis that this may explain the marked resistance of CbA5H to Zn^2+^ inhibition we resorted to a CbA5H variant that we generated and characterized previously. In this variant, H199, which binds an intrinsic Zn^2+^ ion that, at this site, is necessary for stabilizing the homodimer, was exchanged to alanine. The H199A variant is present in the monomeric form while maintaining most of the H_2_ producing activity of the dimeric wildtype protein^43^. When we exposed this variant to ZnCl_2_, its hydrogenase activity indeed decreased by 88%, similar to the inhibition observed for the monomeric [FeFe]-hydrogenases CpI and HydA1 (Figure 3d). Notably, in contrast to the group A [FeFe]-hydrogenases tested here, the intrinsically low activity of ToHydA^29^ was only inhibited by ZnCl_2_ by about 8% (Figure 3d).

### Catalytic importance of H569 and S320 in proton transfer

The results we have described so far show that CpI H565, which is located at the interface of the PTP and the solvent, is required for the metal ions we tested here to exert their inhibitory function, although it apparently is not important for catalytic activity. However, the crystal structures we obtained showed that Zn^2+^ and Ni^2+^ also coordinate to the structurally adjacent H569, and Fe^2+^ as well as Ni^2+^ also coordinate to S320. Both H569 and S320, together with H565, appear to form a kind of entry point to the PTP. We therefore individually replaced H569 and S320 by alanine, which resulted in the two CpI variants H569A and S320A. In contrast to the H565A variant, which had wild type-like H_2_ evolution activity (Figure 3a), the H_2_ production activities of CpI-S320A and CpI-H569A reached only about 10% and 22% of the activity of CpI-WT (Figure 4a). The variants were crystallized employing the protocol established for CpI before^39^, and the structures could be refined to 1.35 Å (S320A) and 1.62 Å (H569A) (Table S5). Overall, both variant proteins showed the same folding as CpI-WT and had full occupancies of their H-clusters (Table S2). The electron densities at positions 320 and 569 show that the original residues S320 and H569 were indeed replaced (Figure 2). Except for this expected observation, no additional noticeable structural differences to the wildtype enzyme including residues of the PTPs were found (Figure S25). Infrared spectra confirmed the H-clusters were mostly not influenced by these exchanges (Figure S12). Thus, the loss of H_2_ production activities in these two variants by 78% to 90% is very likely due to the changes of their side chains, indicating an important functional role of these two side chains for catalytic proton transfer.

**Figure 4.**
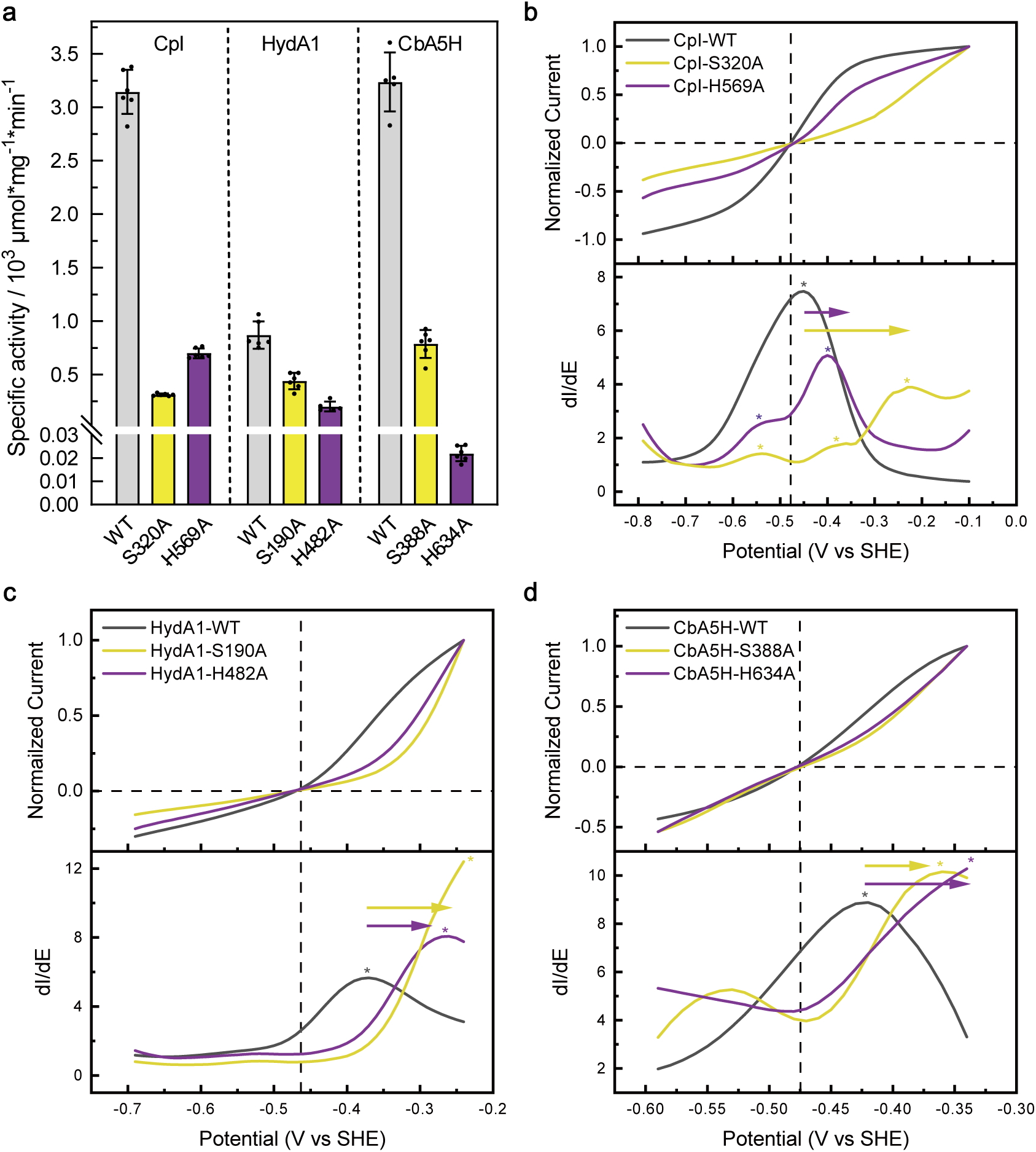
H_2_ evolution activities and electrochemical properties of variants targeting positions that correspond to CpI-H569 and CpI-S320. **a** H_2_ production activities of CpI, HydA1 and CbA5H variants in which the residues that correspond to CpI H569 and S320 were exchanged by alanine were analyzed by the sodium dithionite- and methyl viologen-dependent assay at pH 8 and subsequent gas chromatographic quantification of H_2_. The bars show averages of two biological duplicates (referring to independent protein batches), determined in five to six technical duplicates, and the error bars indicate the standard deviation. **b**, **c** and **d**: CV scans of the same variants (**b**: CpI, **c**: HydA1, **d**: CbA5H) were done as described for Figure 3b except that no ZnCl_2_ was added. The first derivate plots of the CV scans are presented underneath the CV scans. The asterisks show the local maximums in the plots. The arrows illustrate the shifts of local maximums of the variants in comparison to corresponding WT enzymes. The experiments in (**b**, **c** and **d**) were conducted four times from two biological replicates with consistent results.

Previously, we used PFE and CV analyses, respectively, to analyze four mildly active CpI and HydA1 variants of the PTP residues: namely D substitutions at positions of CpI-E279, CpI-E282, HydA1-E141 and HydA1-E144 in HydA1. All the four variants exhibited impaired proton transfer, and we found that, in contrast to the CpI-WT enzyme, significant overpotentials were required for achieving catalytic turnover currents similar to the WT enzyme in both H_2_ production and uptake directions.^33^ The term ‘overpotential’ refers to the potential difference to the equilibrium potential at which the catalytic current is zero, thus can be understood as the necessary driving force for the reaction. The requirement for a higher overpotential observed for the D-substituted variants can be explained by the concept of PCET, during which the simultaneous transport of protons reduces the energy requirement for electron transfer^44^. As a result, electrocatalytic reversibility is impaired when electron and proton transfers become temporally decoupled, which is visible in the CVs as a flattening of the curve around the equilibrium potential E_eq_^33, 45-47^. Therefore, CV analyses are valuable to determine if PCET is affected. When we subjected the CpI variants S320A and H569A to PFE analyses operated in CV mode, notable overpotential requirements were observed in the voltammograms of the S320A variant over the whole pH range tested (pH 5 to pH 9), whereas significant effects were only observed from pH 7 upwards in the case of the H569A variant (Figure S26). The first derivative plot (dI/dE) of a voltammogram aids to observe more subtle effects as it highlights where the current increases most steeply with potential. For CpI WT, the maximum in dI/dE at - 0.454 V corresponds closely to the equilibrium potential E_eq_ (∼ -0.47 V) at pH 8 (Figure 4b). Its single, sharp and high peak, which features a rapid rise and fall in both catalytic directions, highlights a well synchronized PCET with a single dominant rate-determining step and minimal heterogeneity of catalytic states^45, 48^. Regarding the herein investigated PTP variants, a lack in proton transfer efficiency would be manifested by a decreased peak amplitude, an increased peak width and asymmetry in the derivative plots. In general, these features report on kinetic dispersion, i.e. the co-occurrence of multiple partially populated catalytic states, rate-limiting steps (proton transfer and H₂ binding or release) as well as the coupling efficiency between electron transfer and proton transfer^33^.

In the case of the CpI-S320A variant, these changes became apparent in the first derivative plot of the CV recorded at pH 8 (Figure 4b). The absence of a distinct peak at E_eq_ indicates that there is no single, well-defined transition potential and that catalysis takes place over a wide potential range. Moreover, the maximum is shifted to much more positive currents (> -200 mV). In addition to these deviations from the behavior of the CpI-WT enzyme, a splitting of the maximum peak signal and several local maxima can be observed in the case of the CpI-S320A variant, which hints at kinetic dispersion and accumulation of several reduced states (Figure 4b). These features are characteristic for proton-transfer limited catalysis in CV experiments^33^ and indicate that in the CpI-S320A variant, the turnover rate is rather governed by slow proton delivery than fast electron transfer (ET).

Similar but less pronounced effects were observed in the case of the CpI-H569A variant (Figure 4b, Figure S26). At pH 8, the first derivative plot reveals two maxima, with the lower one shifted by -90 mV and the higher one, which appears more asymmetric compared to the signal of CpI-WT, shifted by +50 mV (Figure 4b). Again, the presence of more than one maximum indicates kinetic dispersion. The positive shift of the more prominent maximum suggests a destabilization of reduced catalytic states^49^, while its asymmetry, manifested by a slower decay at positive potentials, suggests altered proton release or reorganization steps (Figure 4b).

Similar results with regard to H_2_ production activity (Figure 4a) and changes of the CV graphs and their first derivative plots (Figure 4c and d, Figure S27) were obtained for variants of HydA1 and CbA5H in which we had exchanged the histidine and serine residues corresponding to CpI-H569 and CpI-S320. The effects appeared even more pronounced than in the CpI variants when analyzing the CVs and first derivative plots for the HydA1 variants compared to CpI (Figure 4c). Especially in the case of HydA1-H482A, the overpotential requirements as well as increased peak width, asymmetry and shifted maxima in the derivative plots almost match those of the HydA1-S190A variant, even at lower pH values (Figure S27). Apparently, catalytic deficiencies imposed by an exchange of this residue in HydA1 cannot be compensated to the same extent as in CpI at low pH values (Figure S26a), perhaps due to the absence of additional [Fe-S] clusters and an electron relay system, respectively.

The corresponding His and Ser exchange variants of CbA5H, which also had reduced H_2_ production activities (Figure 4a), also showed deviations to the CbA5H WT enzyme when analyzed by CV experiments (Figure 4d; Figure S27d-f). However, CV experiments for CbA5H are limited to examinations within a smaller potential window due to its inherent inactivation mechanism at higher potentials,^18, 50^ rendering this hydrogenase not ideal to probe the mechanistic principles of proton transfer in electrochemistry experiments.

CpI-H565 is located directly at the solvent interface, and it is conserved in group A hydrogenases^51^, which implies that it may play a role in proton transfer, too, although the CpI-H565A variant exhibited almost the same H_2_ production activity as the CpI-WT enzyme. Thus, a CV analysis was also applied to CpI-H565A to probe its electrocatalytic properties. Clear overpotential requirements of CpI-H565A were observed in the CV plot, and a less symmetric, lower peak with a shifted maximum in the derivative plot. These effects, however, appeared much less pronounced than observed for variant CpI-H569A (Figure S28).

### CpI-E278-mediated H-bond with CpI-H569 is important for efficient catalysis

Because S320, H569 and, to a lesser extent, H565 seemed important for catalysis, and proton transfer in particular, we examined this structural region more closely. We found that residue E278, which is mostly conserved in group A [FeFe]-hydrogenases^51^, also forms an H-bond with the side chain of H569 (Figure 5), which may imply a functional role of E278 in tuning protonation and deprotonation of H569. The CpI-WT structure shows that the side chain of E278 forms H-bonds with the side chain of H569 and the main chain amide N atom of the P559-G560 peptide bond (Figure 5a). The chemical nature of the Glu side chain suggests that it acts as an H-bond acceptor. We exchanged CpI-E278 with glutamine, whose amide nitrogen of side chain is mostly a hydrogen bond donor, and the E278Q variant exhibited a drop of about 50 % of the H_2_ production activity observed for CpI-WT (Figure 3a). Its 1.31 Å crystal structure reveals local changes in H-bond interactions around position 278 and notably allows for structural distinction of nitrogen and oxygen atoms of the amide side chain of Q278 (Figure 5d). Although the H-bond interactions of Q278 with the side chain of H569 and the main chain of the P559-G560 pair are preserved, their roles as H-bond acceptors and donors are likely swapped, and H569 and the P559-G560 peptide bond now become H-bond acceptors of the newly introduced amide side chain of Q278. A changed H-bond interaction pattern is supported by a 180°-flip of the main chain of P559-G560 and an H-bond between Q278 and the peptide bond carboxyl group (Figure 5). Catalytic proton transfer through H569 would require the second N atom of its side chain to be deprotonated in variant CpI-E278Q. Proton transfer through this putative double-deprotonated state would be less thermodynamically favorable than through the single deprotonated state of the H569 side chain as assumed to occur in CpI-WT. This hypothesis is supported by analyses of the CV and its first derivative of the E278Q variant, which shows a similar but slightly less pronounced effect as observed for variant CpI-H569A (Figure S28).

**Figure 5.**
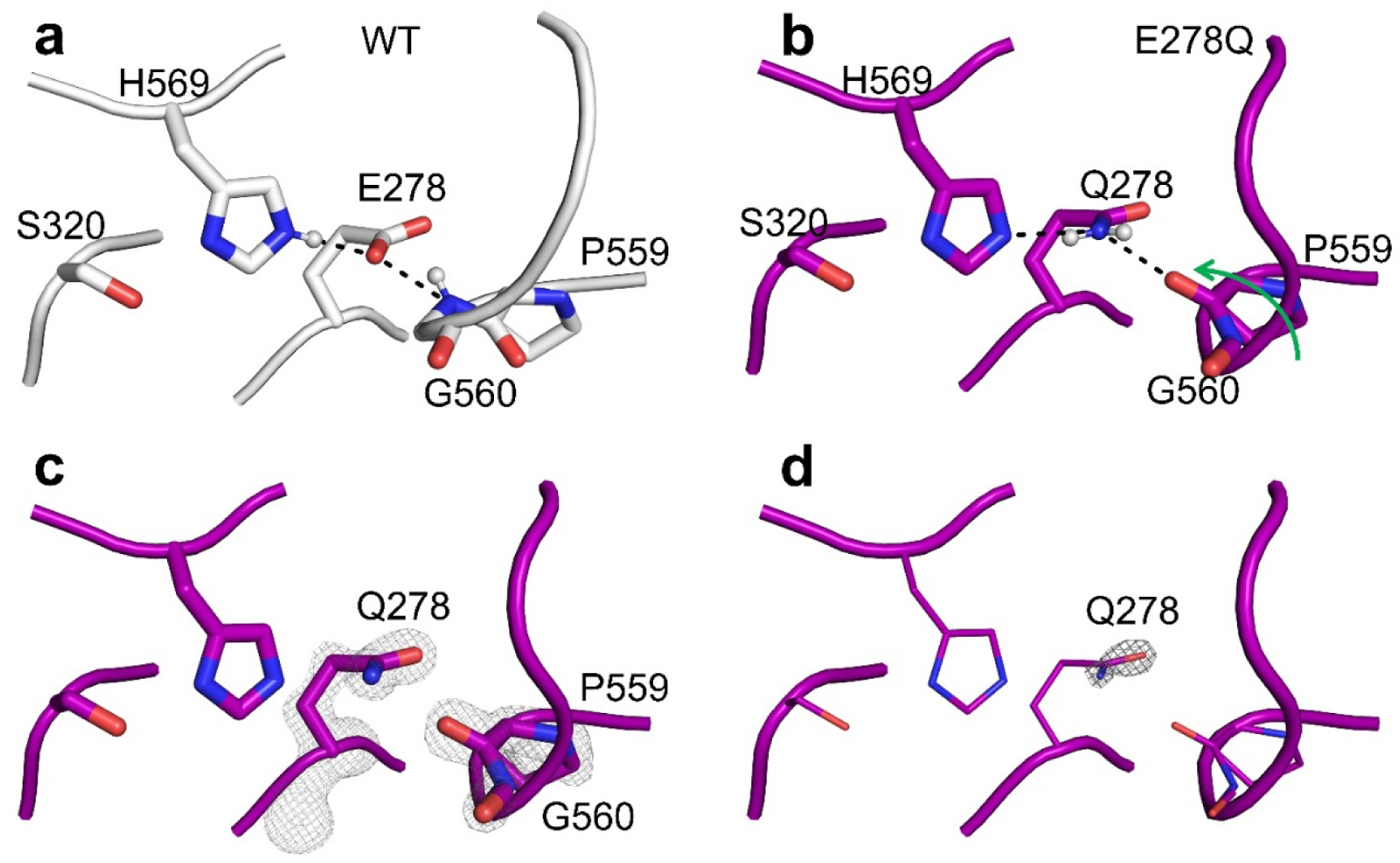
Structural analysis of CpI-E278Q. (**a**) In the structure of CpI-WT (PDB: 4XDC^39^), E278 forms H-bonds with the side chain of H569 and the main chain formed by P559-G560. Here, E278 acts as an acceptor whereas the side chain of H569 and the main chain nitrogen between P559 and G560 function as donors. (**b**) In the structure of the CpI-E278Q variant (PDB: 29EI), Q278 also forms H-bond interactions with H569 and the main chain of P559-G560. The assumed two hydrogen atoms (shown as spheres) of the amide group of Gln suggest that Q278 acts as an H-bond donor while the side chain of H569 and the main chain nitrogen between P559 and G560 act as acceptors. This H-bond interaction pattern requires a 180°-flip of the main chain at this position when compared to the structure of CpI-WT, which is illustrated by the green arrow. Panel (**c**) shows the electron densities of Q278, P559 and G560, and panel (**d**) shows a comparison of the electron densities of the side chain N and O atoms of Q278. The Fo-Fc omit maps were contoured at 6 and 10 σ in panels **c** and **d**, respectively.

### H569-mediated proton transfer dynamics probed by DFT calculations

We employed density functional theory (DFT) calculations to obtain energy profiles of proton transfer (PT) steps through H569. Two models with identical composition were generated starting from the CpI-WT structures mentioned above, PDB 6N59^27^ and 4XDC^39^, which display distinct orientations of H569 (Figure S29). These are denoted here as “IN” when the imidazole ring of H569 points inward toward S320, and as “OUT” when oriented toward H565 and the protein surface, respectively. The PT process was modeled as two steps: H565-H569 and H569-S320. Both “IN” and “OUT” conformations of H569 were computed in the calculations. In both cases, the process was modeled to start from a protonated H565, which transfers the proton to H569 via transition state 1 (TS1), followed by a second transfer through TS2. Each intermediate (Int) and transition state is labeled IN or OUT according to the orientation of H569 (Figure 6).

**Figure 6.**
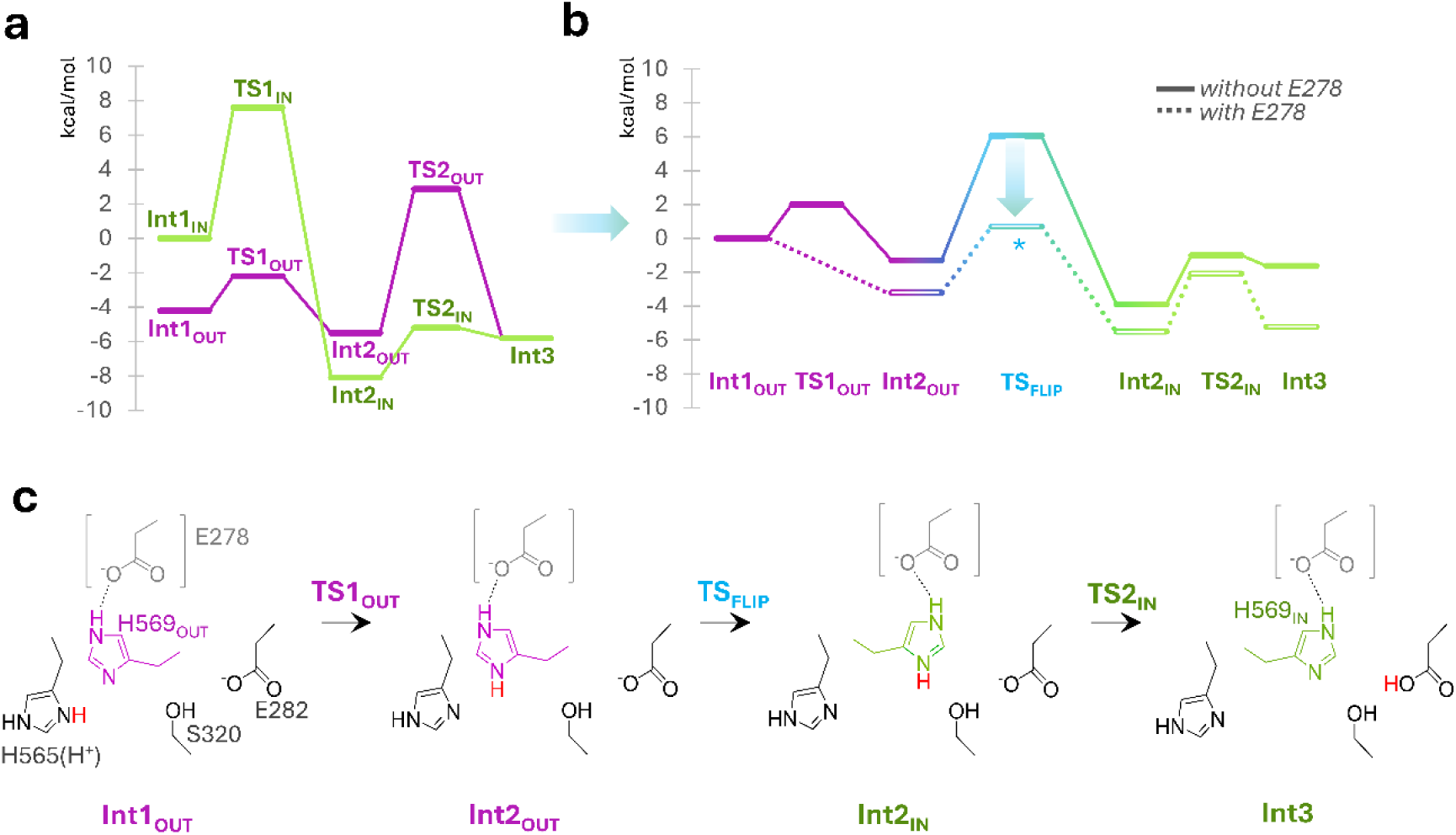
DFT-calculated energy profiles for the PT mechanism. The PT pathway along H565 – H569 – S320 was investigated by DFT considering H569 in the two conformations. (**a**) Two-step PT profiles (H565 → H569 → S320) computed with H569 either in the OUT (purple) or IN (green) conformation throughout the process. (**b**) Composite pathway obtained by combining the lowest-energy steps from panel a: the first PT proceeds via TS_OUT_, whereas the second occurs with H569 in the IN conformation, implying a conformational flip of H569 (TS_FLIP_). The overall profile was recalculated including E278 in the DFT model (dotted line); the corresponding *TS_FLIP_ is indicated with an asterisk to highlight its stabilization. (**c**) Schematic representation of the calculated PT mechanism involving the H569 flip.

The first PT (Int1→Int2, from H565 to H569) was calculated to be significantly more favorable in the OUT conformation (TS1_OUT_ = 2.0 kcal/mol) than in the IN conformation (TS1_IN_ = 7.6 kcal/mol) (Figure 5a). Indeed, although in TS1_IN_, H569 was modeled to rotate slightly toward H565, this movement was less pronounced than calculated in the OUT case, which would probably result in a higher activation barrier (Figure S30). Conversely, the second PT (Int2→Int3, from H569 to S320) is more favorable for the IN (TS2_IN_ = 2.9 kcal/mol) than for the OUT conformation (TS2_OUT_ = 8.3 kcal/mol) (Figure 5a). Thus, the calculations support a first fast PT with H569 in the OUT conformation, followed by a flip of the protonated H569 to the IN orientation, which enables the subsequent fast second PT. The rotational barrier for H569 (TS_FLIP_) was estimated by scanning the Cα–Cβ–Cγ–Cδ torsion of the imidazole ring (Figure S30) and was found to be at 7.4 kcal/mol (Figure 6b), comparable to the highest barriers obtained in the proton transfer profiles shown in Figure 6a. Therefore, although the combined pathway involving TS1_OUT_ followed by TS2_IN_ appears mechanistically reasonable, the barrier calculated for TS_FLIP_ suggests that such a mechanism cannot be regarded as the most favorable one in this model.

However, we noticed that both conformations of H569 are H-bonded with the above-mentioned E278 (Figure 7). E278 seems to ‘anchor’ one side of the side chain of H569 so that the other side alternates between these two conformations for rapid PT. When E278 was explicitly included in the computational model (Figure 6b), the overall energy profile became more favorable. TS1_OUT_ became barrierless, while TS2_IN_ was slightly higher in energy than in the model without E278. This behavior aligns with an expected increase in the pKa of H569 upon formation of a stable H-bond with E278. The resulting energy profile remains accessible in both directions, consistent with the reversibility of PT (and hydrogen oxidation). The most significant result is that the inclusion of E278 reduces the H569 flip barrier by about half (*TS_FLIP_, Figure 6b and Figure S31), from 7.4 to 3.8 kcal/mol. This supports the hypothesis that E278 is important for allowing a rapid PT.

**Figure 7.**
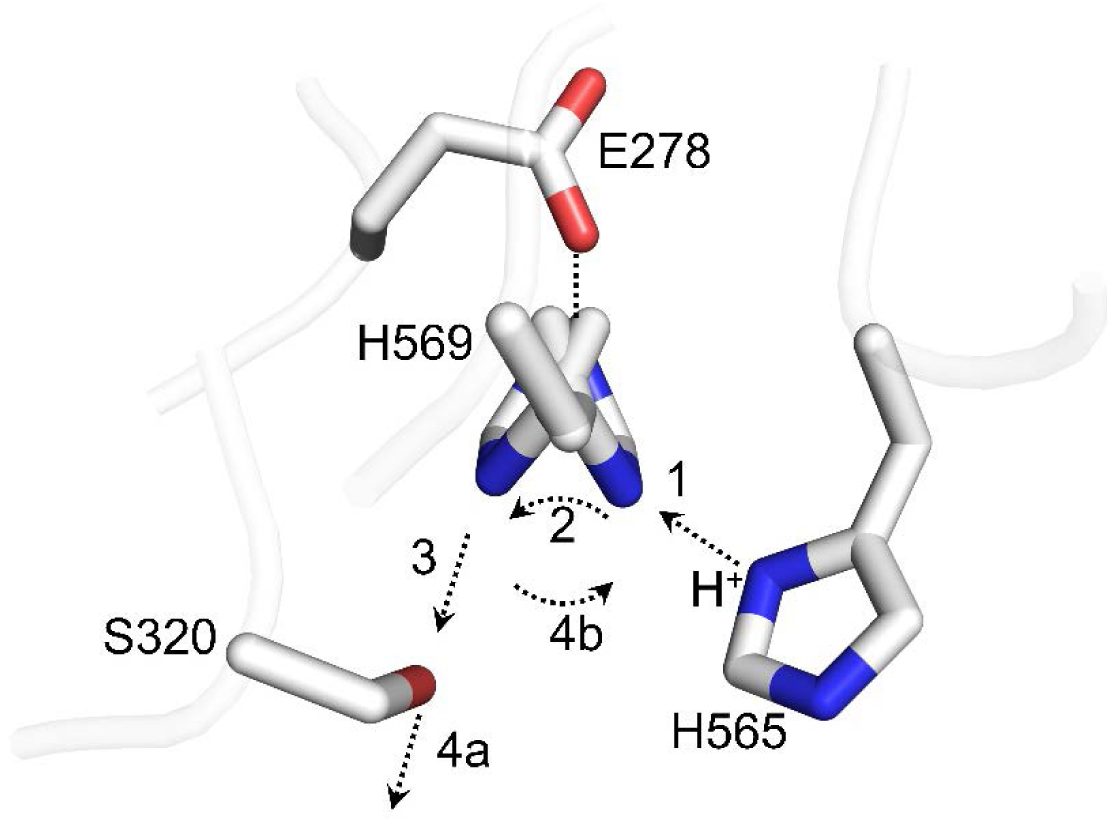
H569-mediated ‘pendulum’ mechanism of proton transfer. When H569 adopts the outward conformation, it is H-bonded with H565 and acts as a proton acceptor. After accepting a proton from H565 (step 1), it changes to the inward conformation (step 2) and is H-bonded with S320 so that it acts as a proton donor to S320 (step 3), that can forward the proton to the PTP (step 4a). In the end, the deprotonated H569 changes back to the outward conformation and is ready for taking up the next proton (step 4b). E278 forms H-bonds with the other side of the side chain of H569 in both conformations. For clarity, the main chains are shown transparently.

## Discussion

### Metal ions exert inhibitory effects by binding to CpI-H565

Inhibitory effects meditated by Zn^2+^ or other metals ions have been investigated with a focus on proton transfer in several other biological systems before, for example cytochrome *c* oxidase^37^, bacterial reaction center^52^ or a voltage-gated proton channel^53^, as well as [FeFe]-hydrogenases^30, 35^. Both cytochrome *c* oxidase and the bacterial photosynthetic reaction center employ histidine residues as proton entry points, and both enzyme complexes could be inhibited by metals ions that bound to these histidine residues^37-38^. In the case of [FeFe]-hydrogenases, the observation that the presence of Zn^2+^ did not decrease the inherently low H_2_ evolution activity of a CpI-E282L variant further led to the suggestion that the metal ion binds to the solvent exposed side chain of E282^30^. It is possible that the apparent resistance of the CpI-E282L variant is due to the very low activity of this variant, which might mask any additional inhibitory effect. We tested this hypothesis by analyzing the effect of ZnCl_2_ on one of our previous PTP variants, CpI-E279A. This variant is strongly affected in proton transport and retains only about 0.1% of the H_2_ production activity of CpI^32^. E279 is buried and therefore very likely not accessible to externally added Zn^2+^ ions so that a relatively active CpI-E279 variant should still be inhibited by zinc. However, the presence of ZnCl_2_ in the reaction mixture had no additional effect on the very low activity of this variant (Figure S31), similar to what was observed for the E282L variant^30^.This observation supports our hypothesis that Zn^2+^ might not inhibit a very compromised CpI variant further or that any effects are not detectable by this assay.

Our crystal structures reveal that Zn^2+^ binds to H565 and H569 (Figure 2), about 5 Å away from E282. Importantly, variant H565A, while possessing WT-like H_2_ production properties, exhibited almost complete resistance to Zn^2+^, while the wildtype enzyme was strongly inhibited by this metal ion. This corroborates the structural snapshots under conditions of active catalysis, and these results do not only provide a molecular understanding of the mechanism of Zn^2+^-mediated inhibition, but also a robust strategy to circumvent the inhibition. Our data also reveal that the Zn^2+^-mediated inhibition shows strong pH-dependence, which is likely due to the necessity of a deprotonation of the coordinating side chains and is reversible in both H_2_ production and uptake.

The data show that the Zn^2+^-meditated inhibitory mechanism can be extended to the other metal ions we tested. Ni^2+^ and Fe^2+^ could also be structurally located at the entrance of the PTP, although their binding modes differed moderately from those of Zn^2+^ (Figure 2). The inhibitory effects these metal ions exerted on hydrogenase activity as determined by PFE at pH 8 (Figure 3; Figures S17 and S18) were in the order Zn^2+^>Ni^2+^>Fe^2+^(>>Mg^2+^). This order followed the well-known Irving-Williams series which stated a general order of binding affinities of metal ions when forming complexes^54^.

### An accessible double-histidine motif might be a Zn^2+^-inhibition signature

By investigating the effect of ZnCl_2_ on additional [FeFe]-hydrogenases, a pattern appeared to emerge that might serve to predict whether a certain [FeFe]-hydrogenase is inhibited by Zn^2+^. From the hydrogenases tested here, those with a freely accessible double-histidine motif at the PTP entrance homologous to H569 and H565 of CpI, namely HydA1 and the monomeric CbA5H-H199A variant, were inhibited by ZnCl_2_ In the case of HydA1, the structure of the Zn^2+^-treated HydA1 protein additionally recapitulates the particular binding mode observed for CpI (Figure S23). In contrast, dimeric wildtype CbA5H was hardly inhibited by zinc, perhaps due to a shielding effect provided by the other protomer. The reduced binding propensity of the PTP entrance of CbA5H to zinc might also suggest that the proton flow from the solvent towards the PTP is moderately different in CbA5H. Notably, the group B [FeFe]-hydrogenase ToHydA was not inhibited by ZnCl_2_ (Figure 3f). As mentioned above, although ToHydA preserves the internal PTP elements, it lacks a counterpart of CpI-H569, and the histidine that aligns to CpI-H565 (Figure S21) appears not to be structurally conserved, according to an AlphaFold model^29, 55^ (Figure S32). This suggests that a Zn^2+^ ion cannot be coordinated at this site of ToHydA because the double-histidine ‘sandwich’ cannot be formed.

### The double-histidine motif of CpI is likely the proton entry point

In addition to inhibitory effects of metal ions, our study also reveals very important insights into catalytic proton transfer of [FeFe]-hydrogenases, with a focus on the surface. Our CpI crystal structures show that Zn^2+^, Ni^2+^ and Fe^2+^ bind to the side chains of H569 and S320, and all binding modes require the involvement of H565. The strongly decreased catalytic activities of variants H569A and S320A suggests that both H569 and S320 are important for catalytic activity (Figure 4a). Notably, the corresponding exchanges in HydA1 and CbA5H also led to enzymes with considerably reduced H_2_ production activities, which was especially pronounced in the case of the CbA5H-H634A variant. The significant overpotential requirements observed during PFE experiments with the variants (Figure 4b-d) indicate that S320 and H569 (and corresponding residues in other group A [FeFe]-hydrogenases) are involved in PCET^33^. In this context it is noteworthy that the coordination modes of the metal ions as observed in the crystal structures reflect deprotonated states of H569 and S320. These would be favored at alkaline pH, which is in line with stronger inhibitory effects of the metals observed at pH 8 than at pH 6 (Figure S18 and S19). The coordination of Fe^2+^ with the hydroxyl group of CpI-S320 is particularly interesting. The side chain of serine possesses very high pKa values and is therefore mostly protonated. This would be even more prominent when it is located at the surface. The deprotonation of S320 is therefore likely tuned by H-bonding with E282 and H569.

According to the closeness of the double-histidine motif to the solvent, it very likely constitutes the proton entry point. Although the CpI-H565A variant was not affected in dithionite-driven H_2_ production assays in solution, moderate but significant effects were observed in the CV analysis of this variant at alkaline pH (Figure S28). Additionally, since H565 is located directly at the interface between PTP and solvent, it seems reasonable to suggest that it plays a significant role in PT, too. Compared with CpI-WT, the presence of new water molecules occupying the original place of the side chain of H565 (Figure S25f) and the increased solvation area of H569 in the H565A variant (0.43 nm^2^, compared to 0.26 nm^2^ in the WT enzyme) may compensate for the absence of H565. A significant role of H565 in PT would indeed explain why it is mostly conserved in group A hydrogenases^51^.

### H569 as a proton-transfer pendulum

Combining the results regarding S320 and the newly identified function of H569 (and corresponding residues in HydA1 and CbA5H) and E278 in catalytic proton transfer leads us to introduce the term ‘SHE-motif’ and to propose a ‘pendulum’ mechanism mediated by H569 that promotes proton exchange between the solvent and the PTP. In this pendulum, although E278 does not appear to be directly involved in catalytic PT, it seems to act as a pivot in ‘anchoring’ one side of side chain of H569. With the stabilization effect exerted by E278, the other side of the H569 side chain is likely to alternate smoothly between the two conformations for efficiently shuttling catalytic protons (Figure 7, supplementary video). Our DFT calculations (Figure 6) support this pendulum mechanism well and also point to the importance of E278 for reasonable energy profiles, which is supported by the reduced H_2_ production activity (Figure 3a) and changed electrocatalytic profile (Figure S28) of the E278Q variant. This pendulum mechanism appears to be operable in variant CpI-H565A, too, because similar conformational alternations of H569 were observed in structures of H565A obtained at pH 4.5 and 7 (Figure S29). Interestingly, this H569-mediated pendulum mechanism shows some similarities with a surface histidine-controlled proton uptake mechanism in cytochrome c oxidase^37^. Indeed, conformational changes of histidines, as well as guidance provided by residues in the vicinity, are fundamental to several enzymes and to PCET^56-57^.

### Does the pendulum-motif correlate with catalytic efficiency?

As mentioned in the introduction, group A [FeFe]-hydrogenases have been studied best, and so far, [FeFe]-hydrogenases from the other phylogenetic groups exhibit considerably lower activities.^23-24, 27-29^ With the exception of CpIII, for which reported activities vary substantially across studies and can reach 10 % of that of CpI, depending on the assay conditions (see also below).^27-28^ Although more members of each group have to be studied to corroborate this trend, and although the PTP is not the only feature that determines catalytic efficiency, as mentioned in the introduction, the observations published so far lead us to speculate on the necessity of the here identified ‘SHE-motif’ for highly active hydrogenases. The prototypical [FeFe]-hydrogenases CpI, CbA5H, DdH and HydA1 are highly active and their PTPs, including the SHE-motifs, are structurally conserved, with an exception that the residue corresponding to CpI-S320 is subject to a conserved replacement by a threonine in DdH (Figure S20). Of particular interest with regard to the findings in this study is H89 of the small subunit of the heterodimeric DdH enzyme, which, based on structural comparisons (Figure S20), is the counterpart of CpI-H569 in the heterodimeric DdH enzyme. The small subunit of DdH has not been identified to be involved in catalysis so far, although it aligns to the C-terminus of CpI^3^. Our results suggest a possible role of the small subunit of DdH in ensuring rapid catalytic proton transfer as identified in CpI, in addition to its possible role in locating the enzyme to the periplasm and its impact on in vitro maturation with the [2Fe]_H_^MIM^ compound^3, 58^. Interestingly, CpI-H565 also has a counterpart in the DdH small subunit (H85^S^; Figure S20), suggesting that this histidine residue indeed might have a functional role as we discuss above. A previous alignment of 828 sequences mostly focused on group A [FeFe]-hydrogenases showed that key residues corresponding to the here proposed pendulum mechanism, CpI-E278, S320, H569 and H565 for the pendulum mechanism are mostly conserved^51^. A survey of nine of these that have been isolated and characterized indeed shows that they are highly active (Figure S33 and Table S7).

In contrast to the monomeric group A [FeFe]-hydrogenases and DdH, group B [FeFe]-hydrogenases such as CpIII (the third [FeFe]-hydrogenase of *C. pasteurianum*) and ToHydA exhibit only about 2-10 % of the H_2_ evolving activity of CpI.^27-29^ Although amino acid differences around the H-cluster or in other parts of the protein may partly explain the observed difference in turnover activities, the absence of an SHE-motif in CpIII and ToHydA, that otherwise share the conserved PTP residues with CpI may contribute to their strongly decreased activities. The group D [FeFe]-hydrogenase TamHydS (a putative sensory hydrogenase from *Thermoanaerobacter mathranii*) is about three thousand-fold less active in H_2_ production than CpI, and it has a very different PTP^24-25^. Very recently, the PTP derived from CpI, but without the -SHE motif, was built into TamHydS. In combination with the conversion of some residues around the [2Fe]_H_ subsite of the H-cluster to residues found in CpI, the resulting TamHydA variant had a 170-fold higher H_2_ production activity^26^. In view of the results we show here, it is well possible that the introduction of the SH-motif could render this TamHydS variant even more active.

## Conclusion

In conclusion, the data shown here reveal the mechanism by which Zn^2+^, Ni^2+^ and Fe^2+^ ions inhibit the [FeFe]-hydrogenase CpI. According to our experiments with additional hydrogenases, and in view of sequence and structural conservation, this mechanism is very likely similar in other group A [FeFe]-hydrogenases. The CpI variant H565A displayed similar catalytic properties as the wildtype enzyme while being resistant to metal inhibition. Thereby, exchanging this residue represents a simple strategy to circumvent metal-dependent inhibition of (group A-) [FeFe]-hydrogenases. The investigation of the crystal structures generated here led us to discover an extension of the prototypical PTP, namely what we term the SHE-motif which contributes to high catalytic activities of several characterized group A [FeFe]-hydrogenases. We hypothesize that the role of the histidine of this motif is a proton-transfer pendulum that allows rapid proton uptake and transport. Notably, H569 is not conserved in [FeFe]-hydrogenases from different phylogenetic classes, and its absence might contribute to the low catalytic activities reported for these enzymes.

## Materials and methods

### Protein preparation

Heterologous protein production and purification were done in anoxic glove boxes (Coy Lab) following a previously published protocol with slight modifications^59^. The construct coding for the specific [2Fe]_H_ (indicated by the suffix - Δ[2Fe]_H_ herein) maturases HydEFG^60^ were not co-expressed, thus the isolated proteins were devoid of [2Fe]_H_ and reconstituted to their holo-forms in vitro (see below). The recombinant proteins harbored C-terminal Strep-tag II peptides if not stated otherwise and were purified using Strep-Tactin affinity chromatography. Subsequently, the purified proteins were maturated by incubating them with a chemically synthesized analogue of [2Fe]_H_ (Fe_2_[μ-(SCH_2_)_2_NH](CN)_2_(CO)_4_[Et_4_N]_2_; abbreviated here as [2Fe]_H_^MIM^)^61^ at a five-fold molar excess for about an hour as described before^62^. Excess of [2Fe]_H_^MIM^ was removed using NAP-5 columns (GE Healthcare). The resulting holoproteins, present in 0.1 M Tris-HCl buffer (pH 8) supplemented with 2 mM sodium dithionite (NaDT), were then concentrated to 15 to 100 mg × ml^-1^ for further use. To obtain Cl**^−^**-free enzymes that were used in experiments shown in Figure S19, an additional buffer exchange of Tris-HCl with potassium phosphate (pH 6.8) was performed. The expression constructs coding enzyme variants were obtained following the published QuikChange site-directed mutagenesis protocol^63^, employing the mismatched oligonucleotides listed in Table S6.

Because the Zn^2+^-binding residues we identified here (H565 and H569 of CpI) are close to the Strep-tag II (residues 577 to 584 in C-terminally tagged recombinant CpI), a second CpI expression construct was designed that introduced an N-terminal Strep-tag II, followed by a TEV (Tobacco Etch Virus) protease cleavage site with the sequence ENLYFQG. After protein production and purification, this N-terminally tagged CpI was incubated with His_6_-tagged TEV protease (New England Biolabs) at 4 °C overnight to cleave the Strep-tag II off. The resulting mixture was applied to a nickel-nitrilotriacetic acid-agarose filled column to remove the His_6_-tagged TEV protease. The eluate, containing tag-less CpI, was then concentrated emplyoing a 50 kDa filter to remove the cleaved short peptide. The obtained CpI proteins (denoted here as no-tag-CpI) were maturated as described above.

### Hydrogenase activity assay

Hydrogenase activity in solution was tested in reaction mixtures containing 400 ng of [FeFe]-hydrogenase, 100 mM NaDT as electron donor and 10 mM methyl viologen as electron mediator. The buffers used for the assay had pH values of pH 6 or pH 8, employing 50 mM MES (2-(N-morpholino) ethanesulfonic acid) for pH 6 and 50 mM HEPES (4-(2-hydroxyethyl) piperazine-1-ethanesulfonic acid) for pH 8. The buffers were supplemented with 500 mM NaCl if not stated otherwise. Other additives such as ZnCl_2_, ZnSO_4_, MgCl_2_ or disodium ethylenediaminetetraacetic acid (EDTA-2Na) were selectively added to the reaction mixtures as needed, and their concentrations are mentioned in the text.

The 2 mL reaction mixtures were prepared in a strictly anoxic glove box in 8 mL headspace flasks and sealed by red rubber Suba seals (Merk). After purging the reaction tubes with Argon for 5 min, they were incubated at 37 °C for 20 min in a shaking water bath. Afterwards, 400 µl of the gaseous headspace of the reaction vessel were injected into a gas chromatograph (model GC-2010-Plus (Shimadzu) equipped with a molesieve 5 Å column (30 m × 0.53 mm, film 25 mm) and a thermal conductivity detector, operated with argon as carrier gas) for quantifying the amount of H_2_ produced.

### Protein crystallization and structure determination

The hanging drop method of vapor diffusion was used to crystallize CpI and HydA1-Δ [2Fe]_H_ in a strictly anoxic glove box. 2 µL of protein solutions at a concentration of 15 mg × mL^-1^ and 2 µL reservoir solution were mixed on a glass cover and equilibrated against reservoir solutions. The chemical compositions of reservoir solutions for each structure obtained here are provided in Table S1. After incubation at 4 °C for about two weeks to achieve full growth, crystals were mounted on loops and flash-frozen in liquid nitrogen. For soaking the crystals with zinc, about 5 mM of ZnCl_2_ were introduced into the crystal drops and the mixtures were incubated for about 3 h before mounting. For soaking with Ni^2+^ and Fe^2+^, final concentrations of 10 mM of NiCl_2_ or FeCl_2_ were added, and soaking was done overnight.

Diffraction data were collected at 100 K in synchrotron beamlines of DESY (Hamburg, Germany) and ESRF (Grenoble, France) as shown in Table S1. The data were processed using XDS^64^. Following the recommendations by Karplus et al.^65^, we applied a generous data cutoff strategy, using the criterion of CC1/2>0.1. The structures filled in the PDB under accession numbers 4XDC^39^ and 6GM5^32^ were used as input models for the procedure of molecular replacement to obtain the phases of CpI and HydA1 structures. Phenix^66^ and Coot^67^ were used for routine refinement and as visualization tools. The structure factors and coordinates have been deposited in the PDB under the accession numbers listed in Table S1. Crystallographic statistics are presented in Tables S3-S5. The identities of metal and chloride ions were confirmed by obtaining anomalous densities from X-ray data collected at certain X-ray energies^40^ as indicated in the respective figure captions and the main text as well as in Tables S3-S5.

### Protein film electrochemistry

Protein film electrochemistry experiments were carried out in a strictly anoxic glove box. The gas-tight electrochemical cell is surrounded by a water jacket and the measuring temperature was kept at 10 °C. The electrochemical cell was equipped with a Ag/AgCl reference electrode under potential control of a PalmSen potentiostat and was connected to a H_2_ purging tube with adjustable flow rates. A 3 M KCl solution was filled into the reference electrode and the standard hydrogen electrode (SHE) potential was adapted based on the equation: E_SHE_ = E_Ag/AgCl_ + 0.2174 V. A platinum wire and a pyrolytic graphite edge rotating disk electrode were used as the counter-electrode and the working electrode respectively. The working electrode was connected to a rotator with a rotation speed of 3000 rpm to ensure efficient mass transport at the interface of the working electrode. 2 – 5 µl of matured [FeFe]-hydrogenases at concentrations of approximately 10 µM were applied on the pre-polished working electrode, which was rinsed with water to remove unbound enzyme molecules after 2 – 5 min of incubation. A mixed buffer containing 15 mM each of MES, HEPES, Na-acetate, TAPS ([tris(hydroxymethyl)methylamino] propanesulfonic acid) and CHES (N-cyclohexyl-2-aminoethanesulfonic acid) supplemented with 100 mM NaCl was pH-adjusted with NaOH or HCl to desired pH (5-9) before use. Other additives such as ZnCl_2_, NiCl_2_, FeSO_4_ and EDTA-2Na were injected into the cell when needed in concentrations indicated in the text. Normalizations and the calculation of the first derivative dI/dE were done in Origin.

### Attenuated Total Reflection-Fourier Transform Infrared (ATR-FTIR) Spectroscopy

A Bruker Tensor II spectrometer (Bruker Optik) equipped with a BioATR cell II (Harrick Scientific) with a double-reflection ZnSe/Si crystal was employed as described before^18^. In brief, scan resolution was set to 2 cm^-1^, and sample incubation as well as measurements were done under a nitrogen gas (N_2_) atmosphere. Samples were dried for 5 to 10 min. Origin was used for data analysis.

### Density Functional Theory (DFT) calculations

Two structural models with identical composition were generated starting from PDB structures 6N59^27^ and 4XDC^38^, which display distinct orientations of H569 (Figure S35). The models therefore differ only in the orientation of this residue. A cluster approach was adopted to describe the proton transfer pathway (PTP), including residues H565, H569, S320, and E282, together with backbone fragments of adjacent residues along the sequence. During geometry optimizations, only these backbone fragments were constrained to preserve the overall structural context, while all side chains directly involved in the proton transfer steps were fully relaxed (Figure S34). An additional set of models including E278 was generated to assess its influence on the reaction energetics. All calculations were performed with the TURBOMOLE 7.4.1 suite^68^ using the BP86 functional^69-70^ with the TZVP basis set^71^. The resolution-of-identity (RI) approximation was used to accelerate Coulomb integral evaluation, and D4 dispersion corrections were included^72^. Environmental effects were treated using the COSMO implicit solvent model^73^ with dielectric constant ε = 40. Proton transfer transition states were located using an eigenvector-following procedure, whereas the conformational flip of H569 was investigated through a relaxed potential energy scan of the Cα–Cβ– Cγ–Cδ dihedral angle of the imidazole ring, performed in 5° increments. The solvent-accessible surface area (SASA) of H569 was calculated using GROMACS^74^ with the gmx sasa tool and a 1.4 Å probe radius (rolling-sphere algorithm).

## Data availability

The coordinates and structural factors of new protein structures resulting from this study have been deposited under the following accession number: 9S7W (https://www.rcsb.org/structure/9S7W), 9S9S (https://www.rcsb.org/structure/9S9S), 9S8H (https://www.rcsb.org/structure/9S8H), 9SAG (https://www.rcsb.org/structure/9SAG), 9S9W (https://www.rcsb.org/structure/9S9W), 9S9V (https://www.rcsb.org/structure/9S9V), 9S9D (https://www.rcsb.org/structure/9S9D), 9S9Z (https://www.rcsb.org/structure/9S9Z), 9T7S(https://www.rcsb.org/structure/9T7S), 9T7O(https://www.rcsb.org/structure/9T7O), 29EI (https://www.rcsb.org/structure/29EI) and 9SAD (https://www.rcsb.org/structure/9SAD) in protein data bank. The electron density maps have been deposited in Figshare. All other data are available in the main text or the supplementary materials. Source data are provided.

## Acknowledgments

J.D. received funding from the German Research Foundation (Deutsche Forschungsgemeinschaft (DFG)) under project number 461338801. E.H., A.H. and T.H. are grateful for funding from DFG Research Training Group GRK2341 “Microbial Substrate Conversion (Micon)”. T.H. and U.P.A. are also funded by the DFG under Germany’s Excellence Strategy–EXC 2033–390677874– RESOLV. T.H. additionally thanks the funding from VolkswagenStiftung (Az 98621). We appreciate Jan Jaenecke and Dr. Martin Winkler for some preliminary experiments and analysis. Dr. Subhasri Ghosh and Cedric Hesse are acknowledged for providing ToHydA protein samples and performing some exploratory experiments, respectively. We acknowledge the following synchrotron radiation facilities for providing beamtime and technical support for X ray diffraction data collection: EMBL Hamburg at DESY (PETRA III, beamlines P13 and P14, Proposal No. MX 875, MX-911, MX987, and MX-1022) and the European Synchrotron Radiation Facility (ESRF, Grenoble, France, beamlines ID23-2, ID30A-3, ID30B, Proposal No. MX 2412, MX-2485, MX-2603, and MX-2696). We thank corresponding beamline scientists for their valuable assistance during the experiments.

## Author contribution

J.D. conceived the project. J.D. and T.H. supervised and led the project. Z.Y. and F. Z. cloned the constructs that were used for expressing the protein variants. Z.Y., L.v.I. and F.Z. performed protein purification, enzymatic activity assays and protein film electrochemistry experiments. J.D., Z.Y. and O.L. analyzed the data from electrochemical experiments. F. Z., Z.Y. and J.D. performed the Infrared experiments. Z.Y., L.v.I. and J.D. crystallized the proteins. J.D., Z.Y. and E.H. collected and processed diffraction data, and performed structural analysis. F.A., C.G. and L.D.G. performed the DFT calculations. J.K. and U.P.A. synthesized the [2Fe]_H_ mimic complex for in vitro maturation. J.D., A.H. T.H., Z.Y., O.L. and F.A. wrote the manuscript with input from other coauthors. All authors approved the manuscript.

