## supporting information for "A histidine-mediated, pendulum-like proton transport mechanism is required for the high catalytic activity of [FeFe]-hydrogenases"

### Content


Figure S1. CpI structures with a focus on the entrance of the proton transfer pathway 4

Figure S2. $Zn^{2+}$ binding modes on the surface of wildtype CpI. 5

Figure S3. Structural assignment of $Zn^{2+}$ and $Cl^-$ ions bound at the entrance of the PTP in wildtype CpI. 6

Figure S4. $Fe^{2+}$ binding mode at the entrance of the PTP of wildtype CpI. 7

Figure S5. Structural assignment of $Fe^{2+}$ ions bound at the entrance of the PTP in wildtype CpI. 8

Figure S6. $Ni^{2+}$ binding at the entrance of the PTP in CpI-WT at different pH values. 9

Figure S7. Structural assignment of $Ni^{2+}$ ions bound at the entrance of the PTP in wildtype CpI. 10

Figure S8. $Ni^{2+}$ binding profiles on the surface of CpI-WT. 11

Figure S9. In vitro $H_2$ production activities of CpI WT and variants 12

Figure S10. Determination of the half-maximal inhibitory concentration ($IC_{50}$) of $ZnCl_2$ for CpI WT. 13

Figure S11. $Zn^{2+}$ binding profiles on the surface of the CpI-H565A variant. 14

Figure S12. Attenuated Total Reflection-Fourier Transform Infrared (ATR-FTIR) spectroscopy on of CpI proteins. 15

Figure S13. Effect of metals on cyclic voltammetry (CV) scans in the absence of enzymes. 16

Figure S14. CV scans of CpI-WT and variants in the absence and presence of $Zn^{2+}$. 17

Figure S15. Effects of EDTA-2Na on the $Zn^{2+}$-mediated inhibition of CpI. 18

Figure S16. Effect of $ZnCl_2$ preincubation and subsequent dilution on $H_2$ production activities of CpI WT and variant H565A. 19

Figure S17. Effect of $FeSO_4$ on hydrogenase activity of CpI WT and variant H565A as determined by PFE experiments. 20

Figure S18. Effect of $NiCl_2$ on the electrochemical activity of CpI-WT and variant H565A. 21

Figure S19. Effect of pH and $Cl^-$ ions on the $Zn^{2+}$-mediated inhibition of CpI WT. 22

Figure S20. Proton transfer pathways in [FeFe]-hydrogenases CpI, CbA5H, HydA1 and DdH. 23

Figure S21. Sequence alignment of CpI, HydA1, CbA5H, ToHydA, TmHydS and TamHydS. 24

Figure S22. Profiles of $Zn^{2+}$ binding on the surface of HydA1-Δ$[2Fe]_H$. 25

Figure S23. $Zn^{2+}$ binding at the entrance of the PTP in HydA1-Δ$[2Fe]_H$. 26

Figure S24. Analysis of the structure of CbA5H with a focus on the PTP entrance. 27

Figure S25. Comparison of proton transfer pathways in structures of CpI-WT and CpI variants S320A, H569A, E278Q and H565A. 28

Figure S26. Comparison of electrocatalytic properties of CpI-WT, CpI-H569A and S320A at pH 5-9. 29

Figure S27. Comparison of electrocatalytic properties of [FeFe]-hydrogenases HydA1 and CbA5H and variants targeting positions that correspond to CpI-H569 and S320 30

Figure S28. Comparison of electrocatalytic properties of CpI-WT and variants CpI-H565A, E278Q and H569A. 31

Figure S29. Two conformations of H569 in structures of CpI-WT and variant H565A under different pH conditions. 32

Figure S30. DFT-optimized structures of key species along the proton transfer pathway. 33

Figure S31. Comparison of the effect of $ZnCl_2$ on $H_2$ production activities of CpI-WT and variant E279A. 34

Figure S32. Proton transfer pathways (PTP) and surface-exposed histidine residues at the entrances of the PTPs in [FeFe]-hydrogenases CpI and ToHydA. 35

Figure S33. Protein sequence alignment of exemplary group A [FeFe]-hydrogenases. 36

Figure S34. Structural models used for DFT calculations. 37

Table S1 Crystallization conditions and beamlines used for X-ray diffraction. 38

Table S2 Root mean square deviations (in Å) of all α-carbon atoms when superimposing the structures with untreated wildtype CpI (a) and HydA1 (b). 39

Table S3 X-ray data collection and refinement statistics of crystal structures CpI-Zn and CpI-Ni-2 40

Table S4 X-ray data collection and refinement statistics of crystal structures CpI-Ni-1, CpI-Fe, CpI-H565A and CpI-H565A-Zn 41

Table S5 X-ray data collection and refinement statistics of crystal structures CpI-S320A, CpI-H569A, CpI-E278Q, CpI-WT-2, and HydA1-Δ$[2Fe]_H$-Zn 42

Table S6. Oligonucleotides employed to generate variants by QuikChange PCR 43

Table S7. Catalytic activities of different [FeFe]-hydrogenases 44

References 45

**Figure S1. CpI structures with a focus on the entrance of the proton transfer pathway**

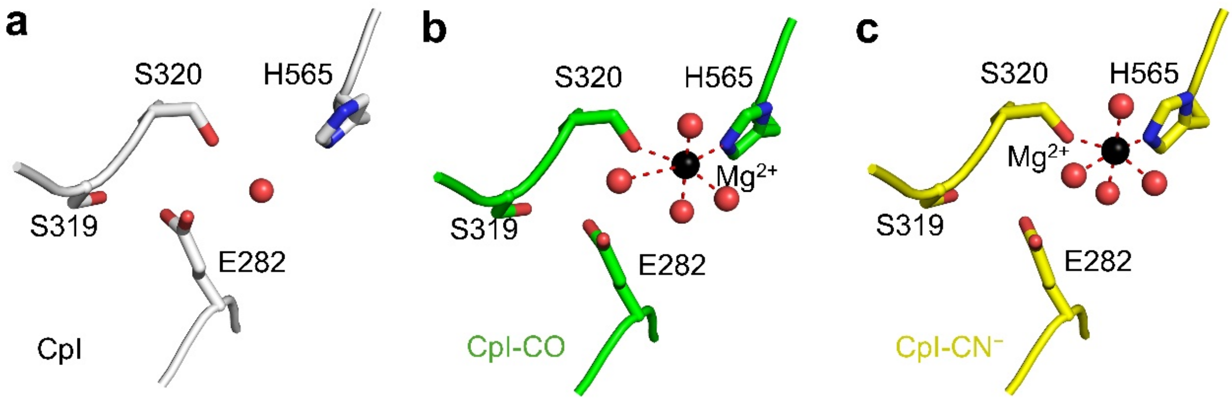


**Figure S1. CpI structures with a focus on the entrance of the proton transfer pathway.** The carbon atoms of structures obtained from untreated (**a**) (PDB ID: 4XDC)[1], CO- (**b**) (PDB ID: 8ALN) and $CN^-$-treated (**c**) (PDB ID: 8AP2) crystals[2] are colored white, green and yellow, respectively. In the structures of CO- and $CN^-$-treated CpI (**b** and **c**), a $Mg^{2+}$ ion (black sphere) was observed to be coordinated by four water molecules (red spheres) and the sidechains of S320 and H565.

**Figure S2. $Zn^{2+}$ binding modes on the surface of wildtype CpI.**

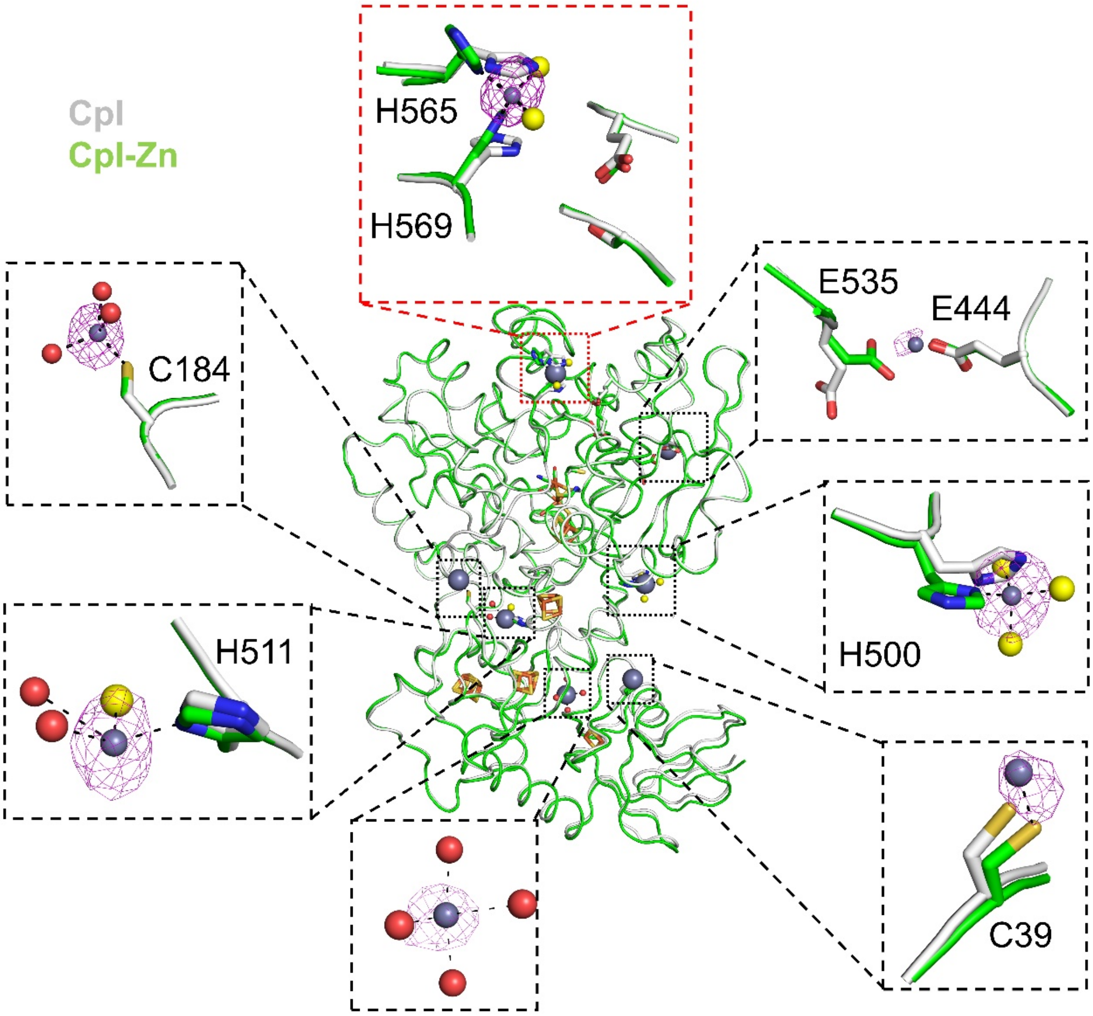


**Figure S2. $Zn^{2+}$ binding modes on the surface of wildtype CpI.** The structure obtained from $Zn^{2+}$-soaked CpI-WT crystals (CpI-Zn; carbon atoms in green; PDB: 9S7W) is superimposed on the structure of untreated CpI (carbon atoms in white; PDB: 4XDC[1]). The anomalous map for the identified $Zn^{2+}$ ions (shown as gray spheres) were contoured at 4 σ. The coordinated water molecules and $Cl^-$ are shown as red and yellow spheres, respectively.

**Figure S3. Structural assignment of $Zn^{2+}$ and $Cl^{-}$ ions bound at the entrance of the PTP in wildtype CpI.**

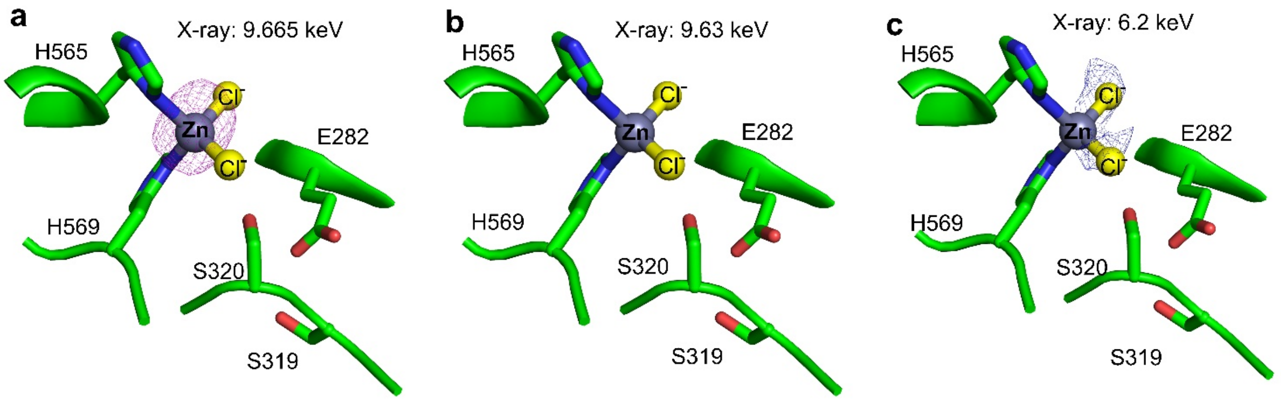


**Figure S3. Structural assignment of $Zn^{2+}$- and $Cl^{-}$ ions bound at the entrance of the PTP in wildtype CpI (CpI-WT).** Three different X-ray energies were used to collect full X-ray datasets of the same CpI-WT $Zn^{2+}$-soaked crystal: 9.665 keV (**a**), 9.63 keV (**b**), 6 keV and 6.2 keV (**c**). Anomalous maps were contoured at 4 σ (in panels **a** and **b**) and 3.5 σ (in **c**). Compared to **a**, the anomalous density of $Zn^{2+}$ is not visible in panels **b** and **c** due to the lower X-ray energies used. In panel **c**, the anomalous densities of $Cl^{-}$ result from two merged datasets collected at X-ray energies of 6 keV and 6.2 keV. More crystallographic details are provided in Table S3.

**Figure S4. $Fe^{2+}$ binding mode at the entrance of the PTP of wildtype CpI.**

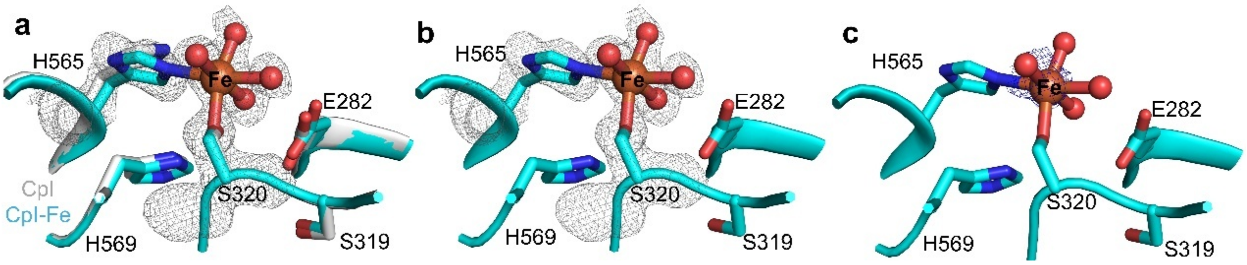


**Figure S4. $Fe^{2+}$ binding mode at the entrance of the PTP of wildtype CpI (CpI-WT).** (**a**)The structure of $Fe^{2+}$-soaked CpI-WT (CpI-Fe; carbon atoms colored in cyan; PDB: 9S9S) is superimposed on the structure of untreated CpI-WT (carbon atoms in white; PDB: 4XDC[1]). The omit map of the structure of CpI-Fe is contoured at 2 σ. For clarity, the structure of CpI-Fe is shown alone in panels **b** and **c**. In panel **c**, the anomalous map of $Fe^{2+}$ from data collected at the X-ray energies of 7.145 keV is contoured at 2 σ.

**Figure S5. Structural assignment of $Fe^{2+}$ ions bound at the entrance of the PTP in wildtype CpI.**

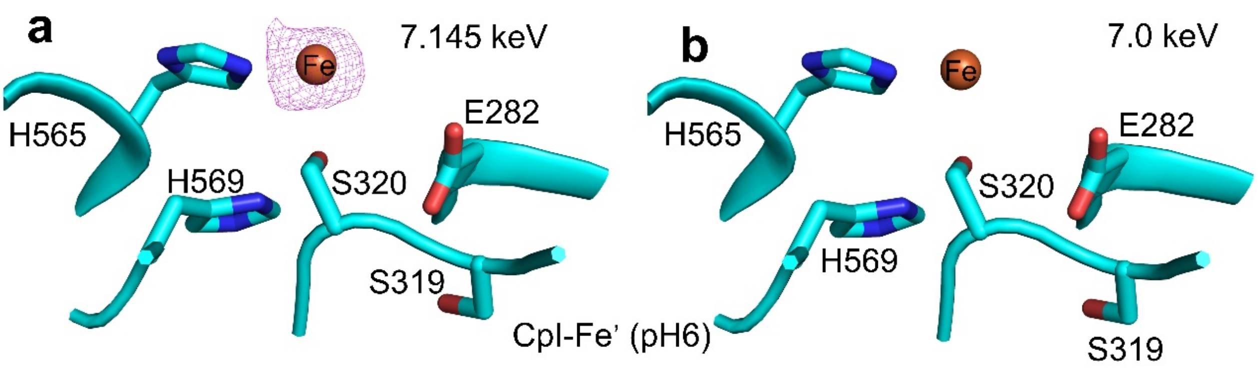


**Figure S5. Structural assignment of $Fe^{2+}$ ions bound at the entrance of the PTP in wildtype CpI.** Two diffraction datasets were collected at X-ray energies of 7.145 keV (**a**) and 7.0 keV (**b**) for a second $Fe^{2+}$-soaked CpI-WT crystal (CpI-Fe'). Anomalous maps were contoured at 3 σ.

**Figure S6. $Ni^{2+}$ binding at the entrance of the PTP in CpI-WT at different pH values.**

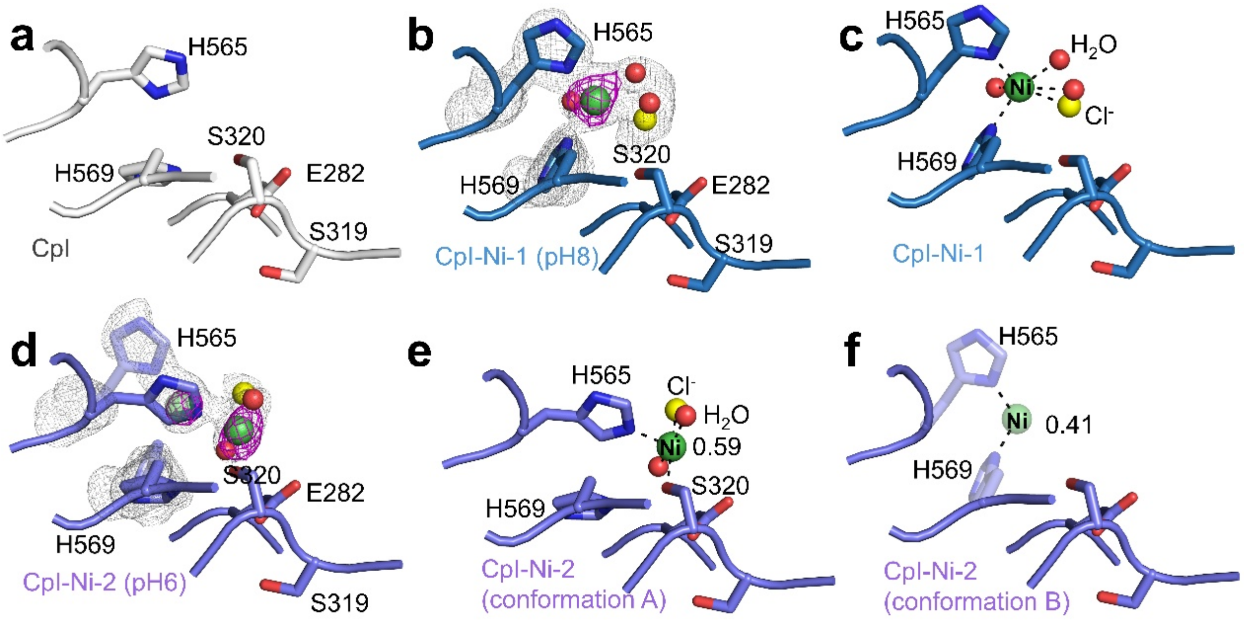


**Figure S6. $Ni^{2+}$ binding at the entrance of the PTP in CpI-WT at different pH values.** The structures of wildtype CpI obtained by soaking crystals with $Ni^{2+}$ at pH 8 (CpI-Ni-1; PDB: 9S8H) and at pH 6 (CpI-Ni-2; PDB: 9SAG) show different binding modes of the $Ni^{2+}$ ion at the entrance of the PTP. After soaking at pH 8, the $Ni^{2+}$ ion is coordinated by three water molecules, a $Cl^-$ ion and the side chains of H565 and H569 (**b** and **c**). After soaking at pH 6, two $Ni^{2+}$ binding modes were observed as two conformations that are overlain in panel **d** (also see Figure S7). Conformation A with an occupancy of about 0.59 is shown opaquely, and conformation B with an occupancy of 0.41 in a transparent mode. In conformation A, the $Ni^{2+}$ ion is coordinated by the side chains of H565 and S320 as well as two water molecules and a $Cl^-$ ion (**e**). In conformation B (shown in **f**), the $Ni^{2+}$ ion is coordinated by the side chains of H565 and H569, and the conformation is very similar to that observed in CpI-Ni-1 (**c**). Because these two conformations overlap, the coordinating partners of $Ni^{2+}$ such as water molecules and $Cl^-$ could not be modelled conclusively in CpI-Ni-2. The anomalous maps (pink mesh in **b** and **d**) of $Ni^{2+}$ ions (shown as green spheres) were contoured at 3.5 σ. The anomalous densities of $Ni^{2+}$ ions in CpI-Ni-1 and CpI-Ni-2 were obtained from data collected at an X-ray energy of 8.35 keV. More crystallographic details are provided in Table S3 and S4. The occupancies in panels **d**, **e**, and **f** were determined by employing iterative group occupancy refinement until the occupancies converged (<0.05). Because the $Ni^{2+}$ ions and the histidine side chains each adopt two conformations (A and B), two separate refinement groups were defined for conformations A and B, with the sum of occupancies within each group constrained to 1.

**Figure S7. Structural assignment of $Ni^{2+}$ ions bound at the entrance of the PTP in wildtype CpI.**

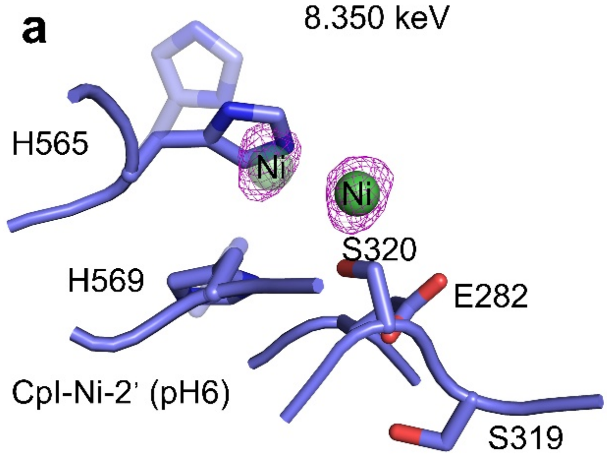


**Figure S7. Structural assignment of $Ni^{2+}$ ions bound at the entrance of the PTP in wildtype CpI.** Two diffraction datasets were collected at X-ray energies of 8.35 keV (**a**) and 8.335 keV (**b**) for a second $Ni^{2+}$-soaked CpI-WT crystal at pH 6 (CpI-Ni-2'). Anomalous maps were contoured at 3 σ.

**Figure S8. $Ni^{2+}$ binding profiles on the surface of CpI-WT.**

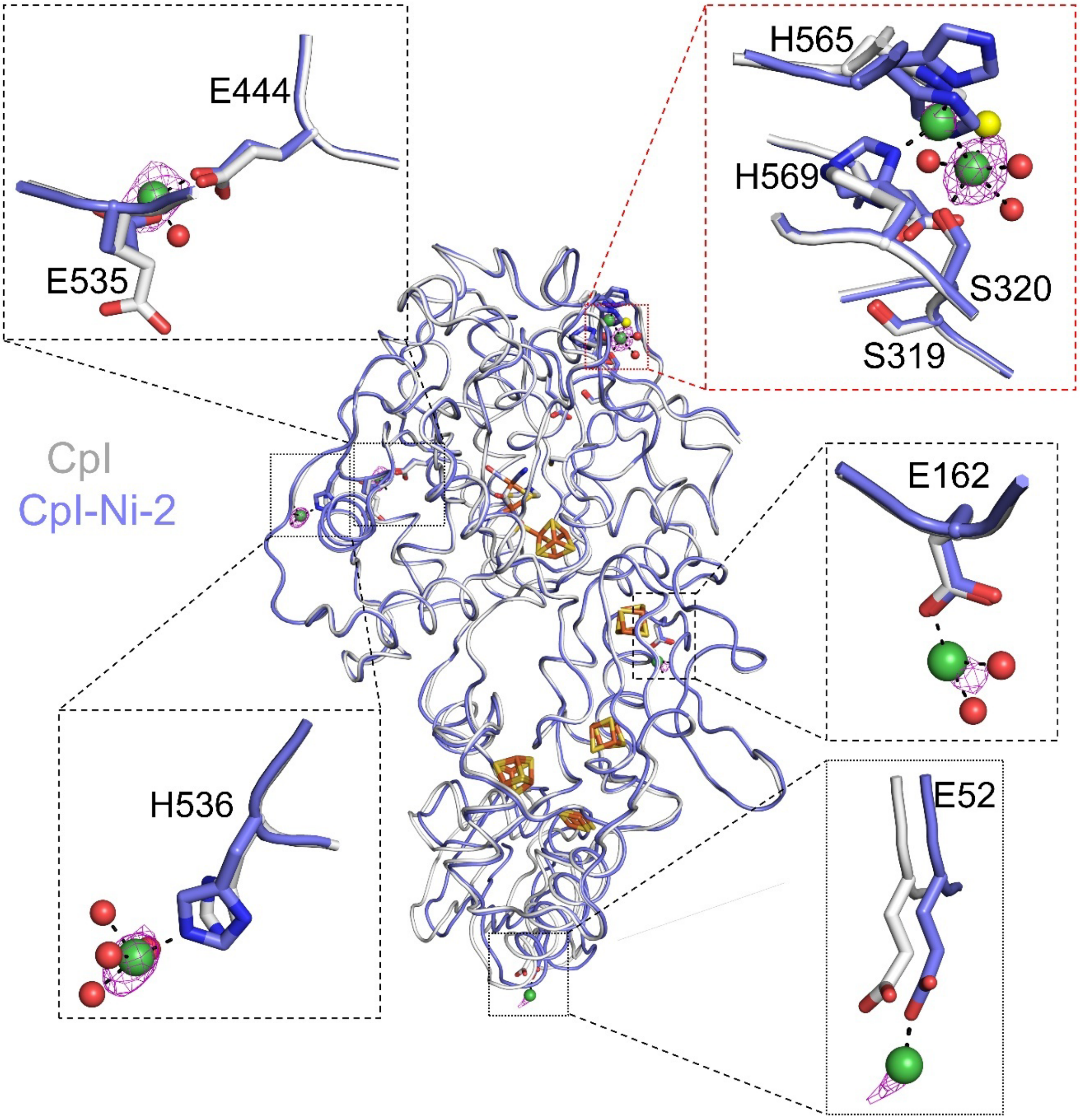


**Figure S8. $Ni^{2+}$ binding profiles on the surface of CpI-WT.** The structure of CpI-WT soaked with $Ni^{2+}$ at pH 6 (also see Figures S6-S7; CpI-Ni-2, carbon atoms colored sky-blue; PDB: 9SAG) is superimposed on that of untreated CpI crystals (white carbon atoms; PDB: 4XDC[1]). The anomalous maps (pink mesh) for the identified $Ni^{2+}$ ions (shown as green spheres) were contoured at 4 σ. Coordinated water molecules and $Cl^-$ ions are shown as red and yellow spheres, respectively.

**Figure S9. In vitro $H_2$ production activities of CpI WT and variants**

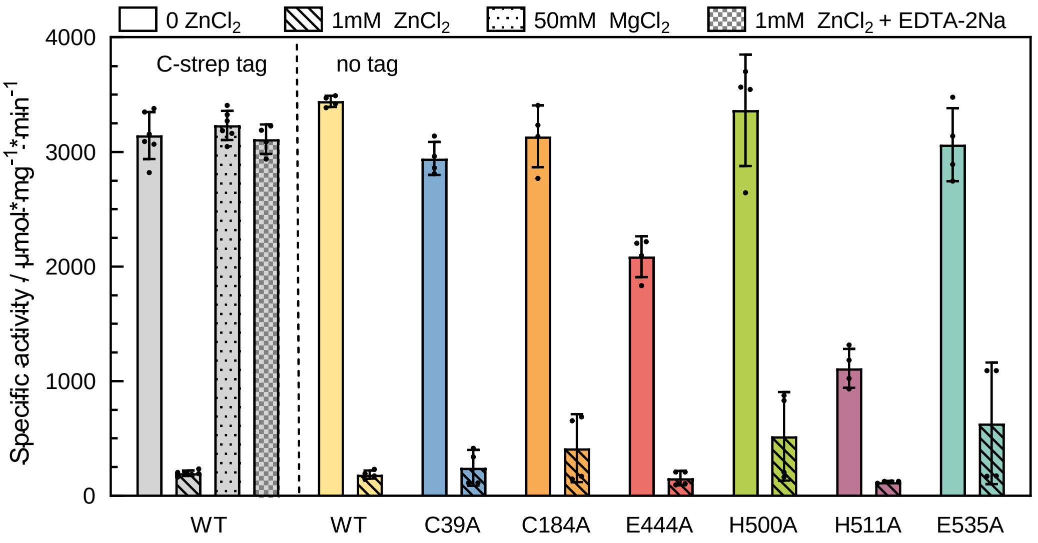


**Figure S9. In vitro $H_2$ production activities of CpI-WT and variants.** $H_2$ production activities of CpI-WT (WT; gray bars) were determined in a dithionite-driven, methyl viologen-mediated hydrogenase activity assay at pH 8 in the absence (plain bars) or the presence of 1 mM $ZnCl_2$ (striped bars), 50 mM $MgCl_2$ (dotted bars) or with a combination of 1mM $ZnCl_2$ and 1mM EDTA-2Na (checkered bars). To the right side of the dashed line, $H_2$ production activities of CpI-WT and variants C39A, C184A, E444A, H500A, H511A and E535A in the absence (plain bars) and presence of 1 mM $ZnCl_2$ (striped bars) were determined. Note that the Strep II-tags of these proteins were removed by TEV digestion after protein purification (indicated by the 'no tag' label at the top). Details are described in the experimental section. The bars show averages of two biological duplicates (referring to independent protein batches), determined in four to six technical duplicates, and the error bars indicate the standard deviation.

**Figure S10. Determination of the half-maximal inhibitory concentration ($IC_{50}$) of $ZnCl_2$ for CpI WT.**

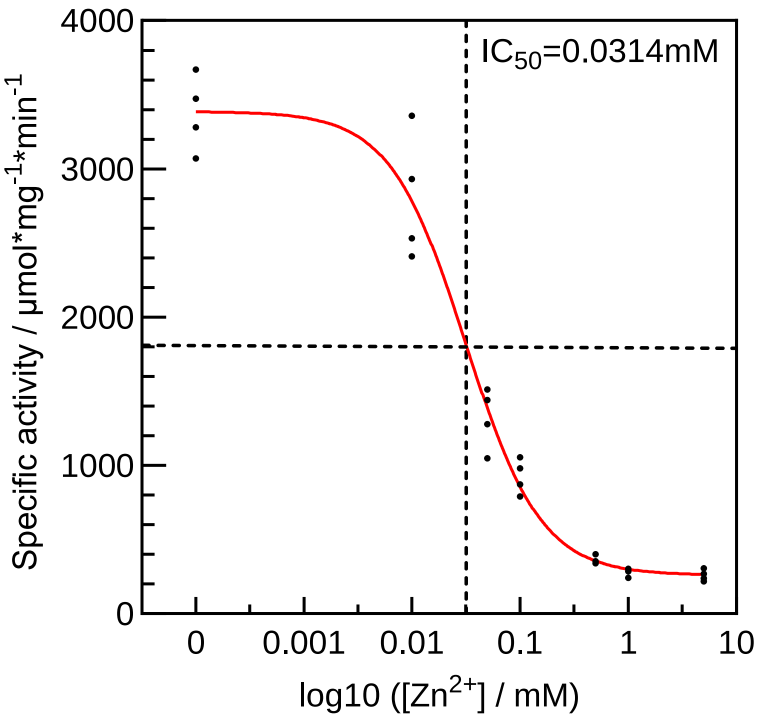


**Figure S10. Determination of the half-maximal inhibitory concentration ($IC_{50}$) of $ZnCl_2$ for CpI WT.** $H_2$ production activities of CpI-WT were determined in a dithionite-driven, methyl viologen-mediated hydrogenase activity assay at pH 8, with the supplement of $ZnCl_2$ in the concentrations of 0, 0.01, 0.05, 0.1, 0.5, 1 and 5 mM. Measurements were performed in quadruplicate (n=4) (two biological replicates times two technical replicates). $IC_{50}$ value was calculated using the GraphPad Prism software (GraphPad Software, Boston, MA, USA). Data were analyzed using a non-linear regression model based on the four-parameter logistic equation (log(inhibitor) vs. response -- Variable slope). The "Top" parameter was constrained to 100%, while the "Bottom" parameter was left unconstrained to account for the observed residual activity at high metal concentrations. The concentration of 0 mM $ZnCl_2$ was handled by assigning a nominal low value $10^{-5}$ mM to permit logarithmic transformation.

**Figure S11. $Zn^{2+}$ binding profiles on the surface of the CpI-H565A variant.**

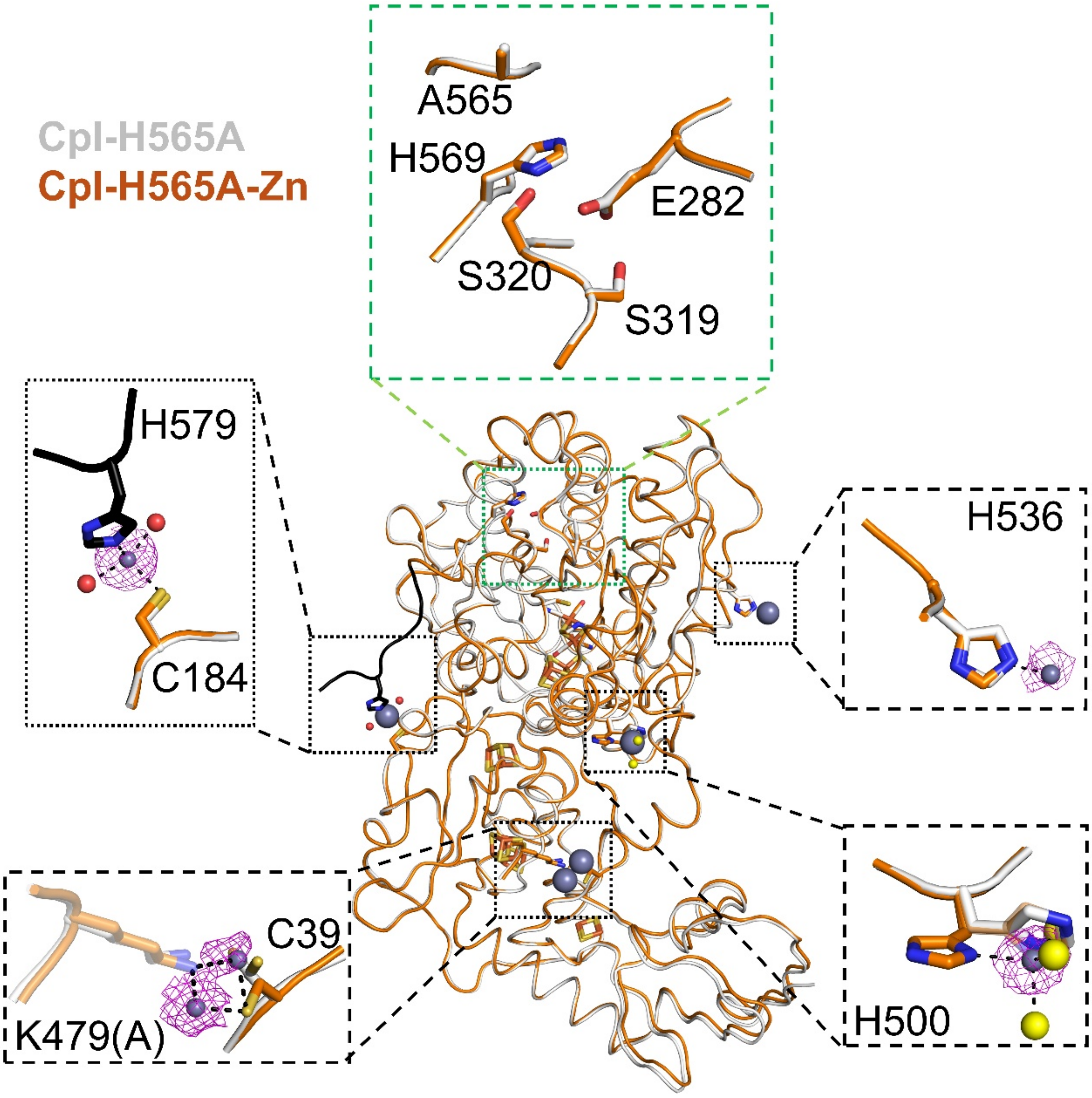


**Figure S11. $Zn^{2+}$ binding profiles on the surface of the CpI-H565A variant.** The structure obtained from $Zn^{2+}$-soaked CpI-H565A crystals (chain B; orange carbon atoms; PDB: 9S9Z) is superimposed on the structure of untreated Cp-H565A (white carbon atoms; PDB: 9S9D). The anomalous maps of the identified $Zn^{2+}$ ions (shown as gray spheres) were contoured at 4 σ and are shown as pink mesh. Coordinated water molecules and $Cl^-$ ions are depicted as red and yellow spheres, respectively. Note that a conformational change of the C-terminus (to the left; carbon atoms colored black) in the $Zn^{2+}$-bound H565A variant was observed that is apparently a consequence of a joint coordination of $Zn^{2+}$ by the side chains of C184 and H579. This conformation was only observed in chain B. K479(A) (lower left corner) refers to K479 in chain A and is shown here because it coordinates two $Zn^{2+}$ ions together with C39 from chain B.

**Figure S12. Attenuated Total Reflection-Fourier Transform Infrared (ATR-FTIR) spectroscopy on of CpI proteins.**

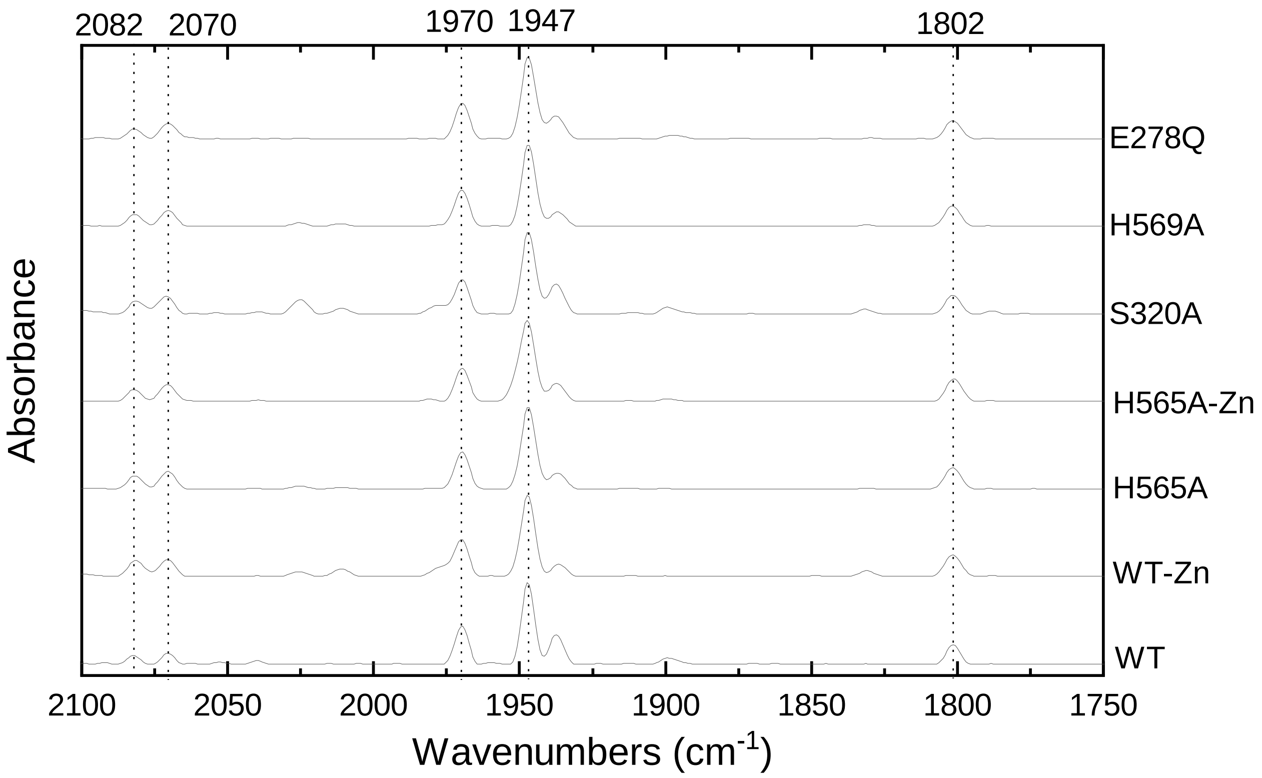


**Figure S12. Attenuated Total Reflection-Fourier Transform Infrared (ATR-FTIR) spectroscopy on of CpI proteins.** Infrared spectra of CpI-WT and variants H565A, S320A, H569A and E278Q were collected under a nitrogen gas ($N_2$) atmosphere as described in the materials and methods section and normalized to the same intensities at 1947 $cm^{-1}$. In the case of CpI-WT and variant CpI-H565A, spectra were also collected after mixing the protein samples with 5 mM $ZnCl_2$. The peaks at 2082, 2070, 1970, 1947 and 1802 $cm^{-1}$ represent an oxidized H-cluster[3]. The spectra were collected using attenuated total reflectance Fourier transform infrared spectroscopy method and on a Bruker Tensor II spectrometer (Bruker Optik, Germany).

**Figure S13. Effect of metals on cyclic voltammetry (CV) scans in the absence of enzymes.**

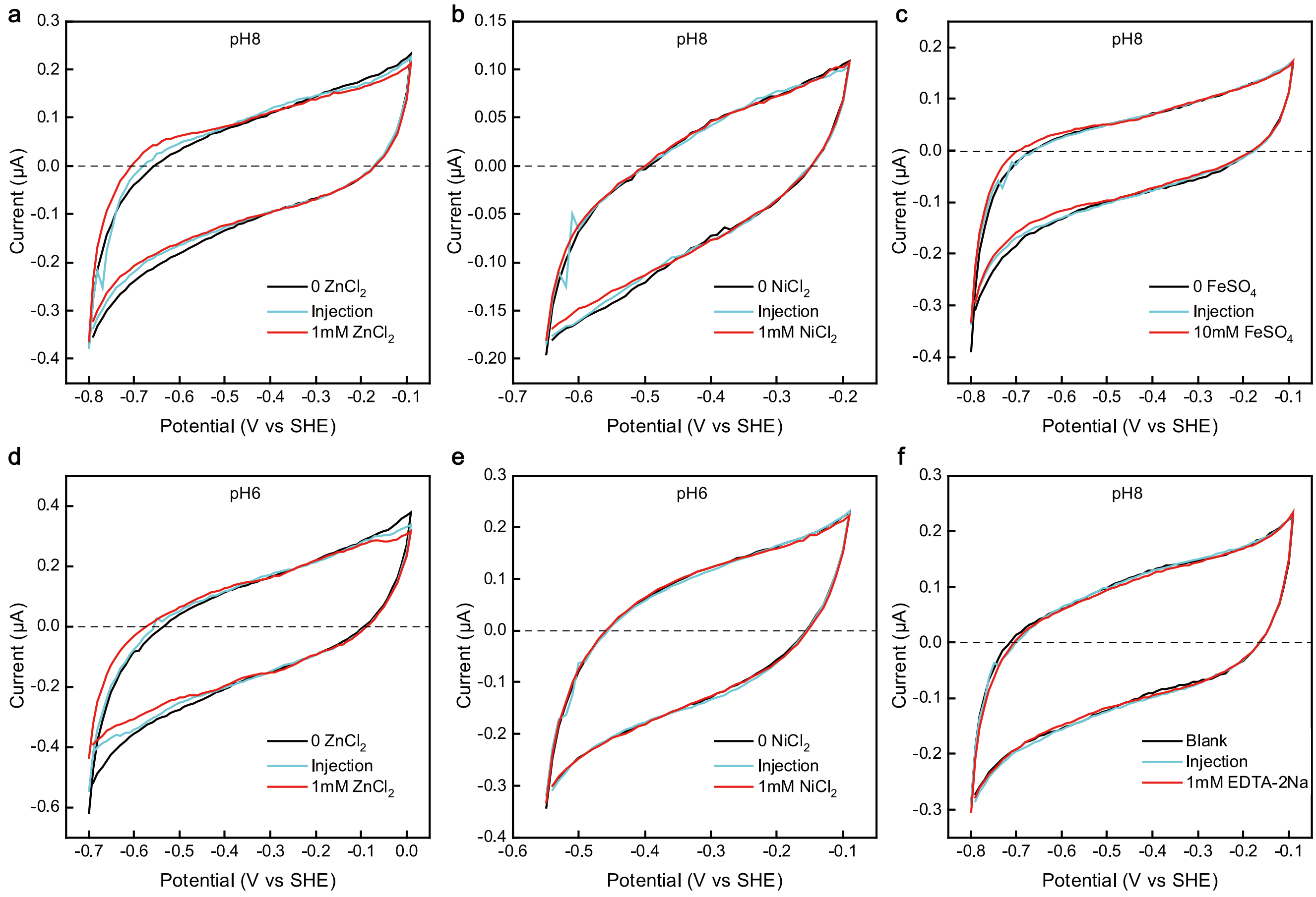


**Figure S13. Effect of metals on cyclic voltammetry (CV) scans in the absence of enzymes.** In each case, three CV scans, namely before (black lines), during (blue lines) and after the injection of metal salt solutions (red lines) were performed. Metal ions were injected into the buffer of the electrochemical cell during the second CV scan. To minimize the precipitation of $Fe^{2+}$ and $Ni^{2+}$, smaller potential ranges were applied in experiments in which $FeSO_4$ or $NiCl_2$ were used than in those applied for analyses of the effect of $ZnCl_2$ or EDTA-2Na. The CV scans were performed under the following experimental conditions: working electrode rotation at 3000 rpm, temperature of 10 °C, $H_2$ flow rate of 2 L × $min^{-1}$ and scan rate of 10 mV × $s^{-1}$. A mixed buffer system was employed to adjust the pH to either pH 8 (**a**, **b**, **c**, **f**) or pH 6 (**d**, **e**) as indicated at the top of each panel.

**Figure S14. CV scans of CpI-WT and variants in the absence and presence of $Zn^{2+}$.**

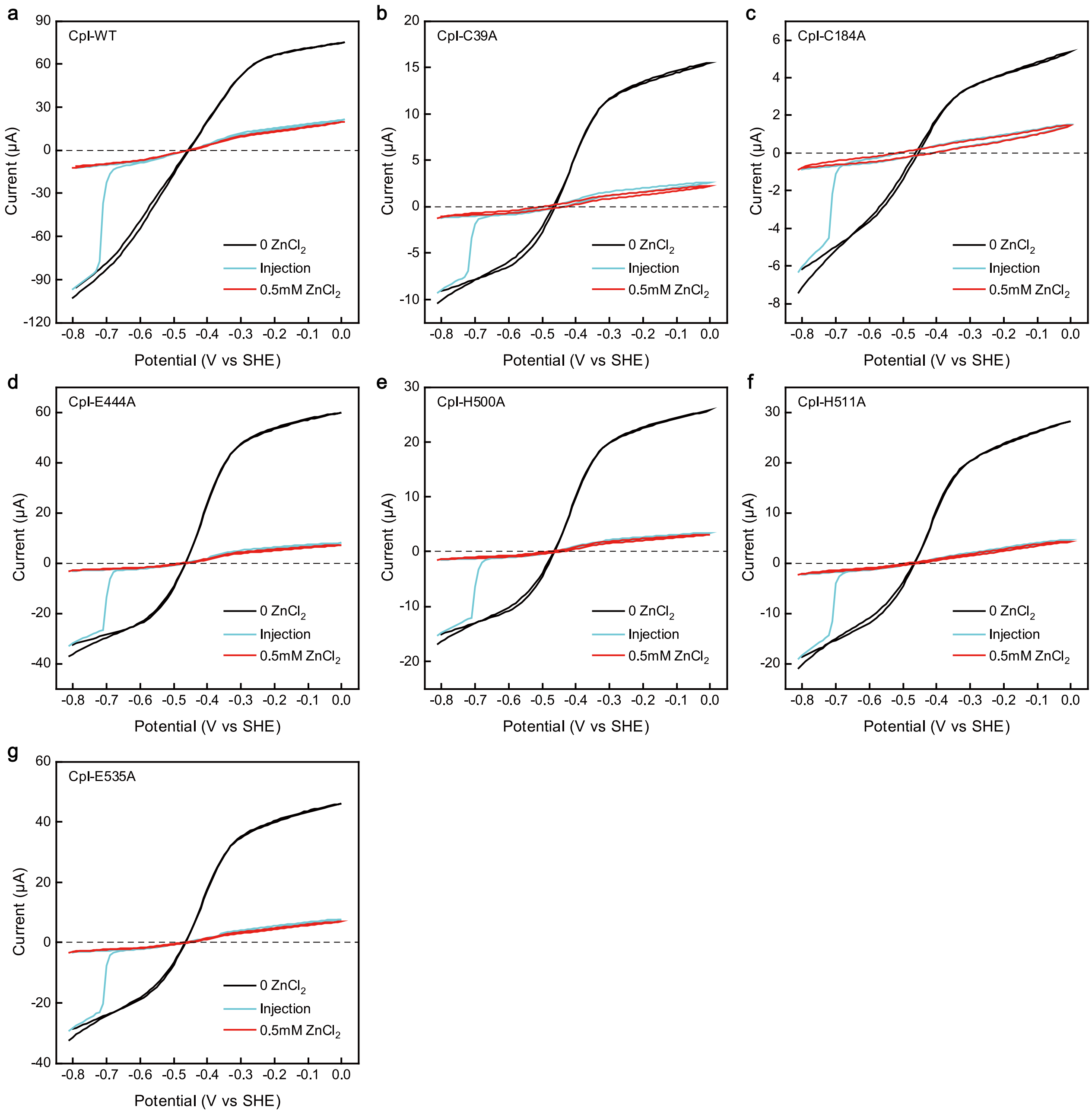


**Figure S14. CV scans of CpI-WT and variants in the absence and presence of $Zn^{2+}$.** CpI variants with exchanges of amino acids involved in $Zn^{2+}$ coordination apart from H565 were tested for inhibition by $ZnCl_2$ through cyclic voltammetry (CV) at pH 8. The exchanges are indicated at the top of each panel. All enzymes were analyzed after their Strep II-tags had been removed. Three CV scans were performed, one before $ZnCl_2$ injection (black lines), one during injection of 0.5 mM $ZnCl_2$ (blue lines) and one after that (red lines). The experimental conditions were the same as described in the caption of Figure S13, except that a $H_2$ flow rate of 30 L × $min^{-1}$ was applied.

**Figure S15. Effects of EDTA-2Na on the $Zn^{2+}$-mediated inhibition of CpI.**

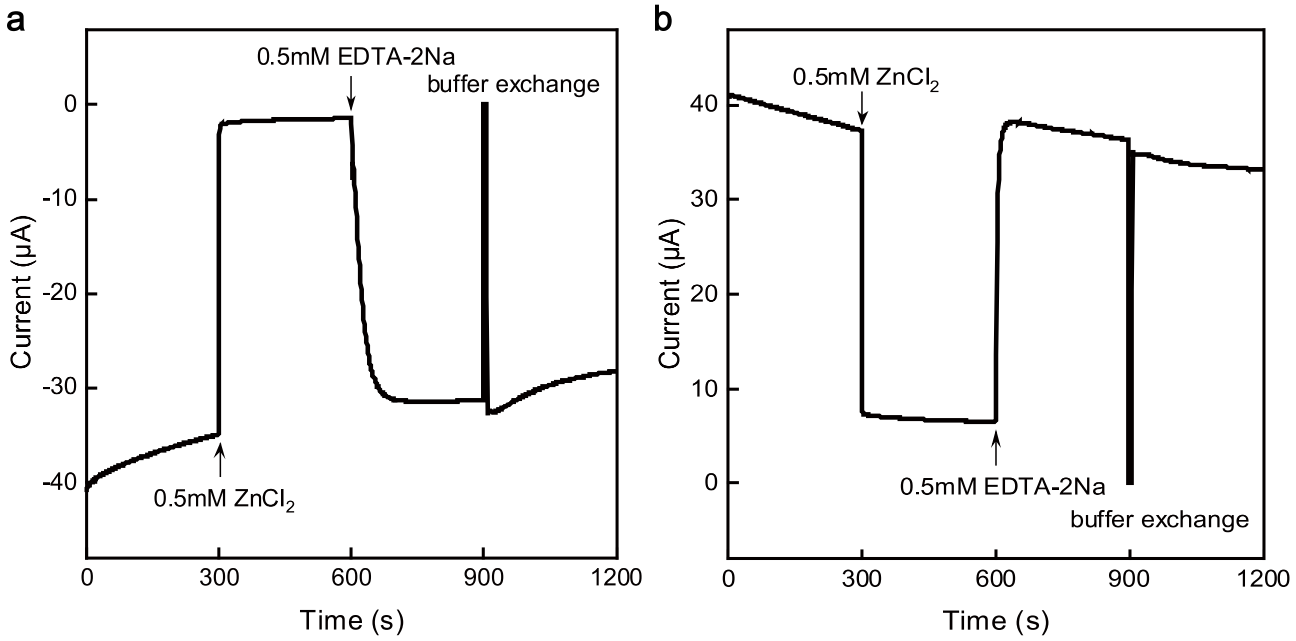


**Figure S15. Effects of EDTA-2Na on the $Zn^{2+}$-mediated inhibition of CpI.** The CpI-WT enzyme was applied to the electrode and electrochemical chronoamperometric currents were recorded at a potential of –0.8 V (**a**) or –0.1 V (**b**). After 300 s, 0.5 mM $ZnCl_2$ was injected. After 600 s, 0.5 mM EDTA-$Na_2$ was injected, and after 900 s, the buffer was exchanged to $ZnCl_2$-free buffer. The experiments were done under the following experimental conditions: pH 8, working electrode rotation at 3000 rpm, temperature of 10 °C, $H_2$ flow rate of 0 and 30 L × $min^{-1}$ in (**a**) and (**b**), respectively.

**Figure S16. Effect of $ZnCl_2$ preincubation and subsequent dilution on $H_2$ production activities of CpI WT and variant H565A.**

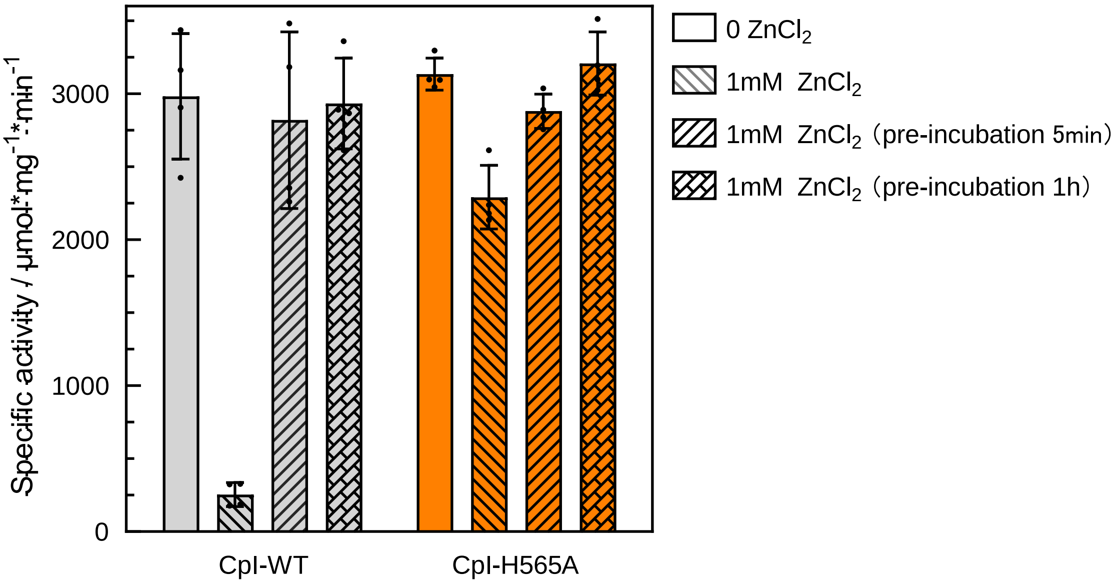


**Figure S16. Effect of $ZnCl_2$ preincubation and subsequent dilution on $H_2$ production activities of CpI WT and variant H565A.** $H_2$ production activities of 400 ng each of the CpI-WT enzyme (gray bars) and the CpI H565A variant (orange bars) were determined employing the dithionite- and methyl viologen-dependent hydrogenase assay in the absence (plain bars) or presence of 1 mM $ZnCl_2$ (bars striped from upper left to lower right). To test for a reversibility of $Zn^{2+}$ inhibition by dilution, each protein was additionally preincubated with 1 mM $ZnCl_2$ for 5 min (bars striped from upper right to lower left) or 1 h (brick-patterned bars). Afterwards, 2 µL of these enzyme mixtures were added to 2 mL of the otherwise $Zn^{2+}$-free hydrogenase assay reaction mixture (50 mM HEPES, pH 8, 500 mM NaCl, 100 mM sodium dithionite, 10 mM methyl viologen). In all cases, the hydrogenase assay mixtures were incubated at 37 °C for 20 min. The experiments were done with biological duplicates, determined in four technical duplicates. The bars show the averages, and the error bars indicate the standard deviation.

**Figure S17. Effect of $FeSO_4$ on hydrogenase activity of CpI WT and variant H565A as determined by PFE experiments.**

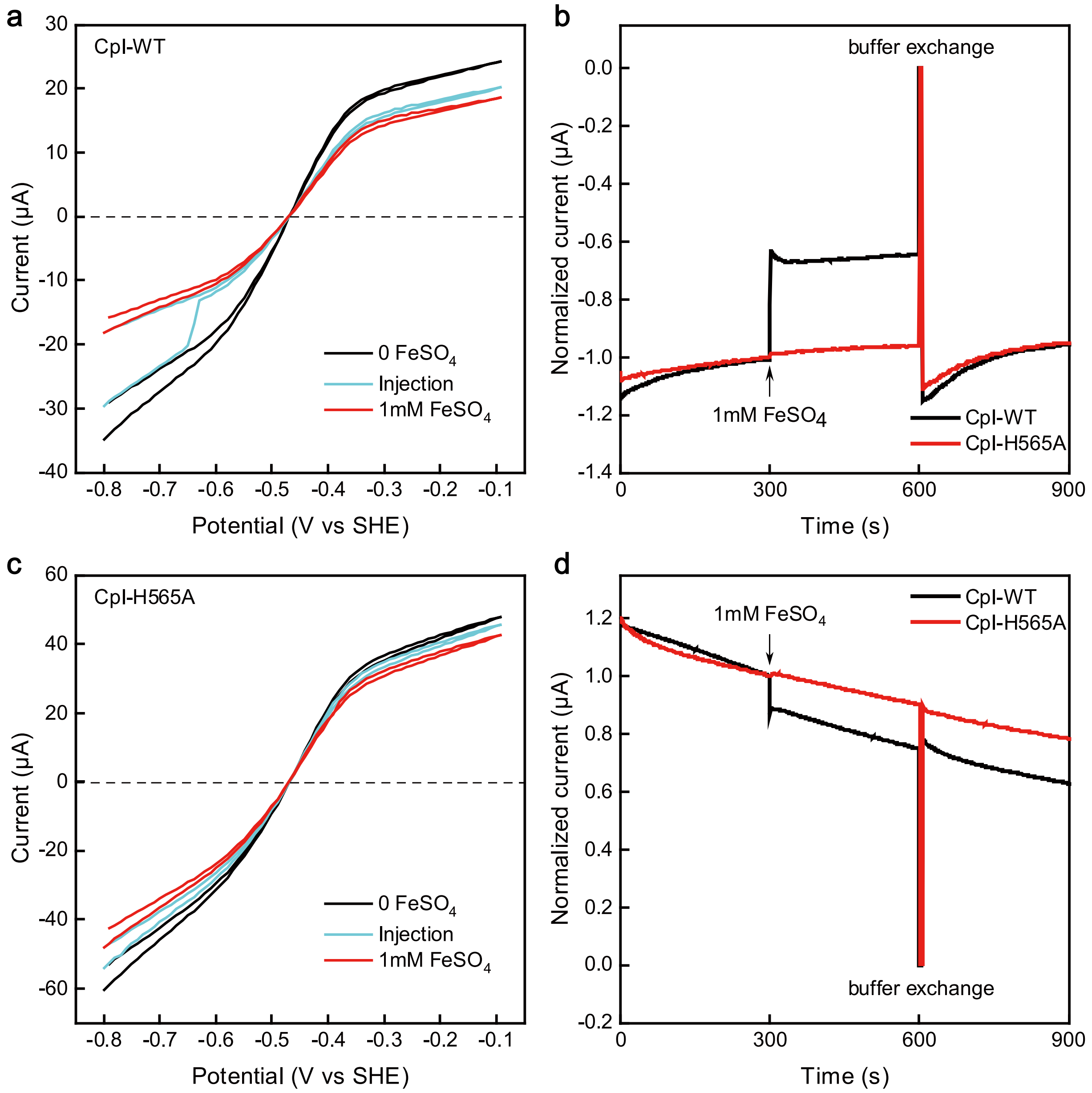


**Figure S17. Effect of $FeSO_4$ on hydrogenase activity of CpI WT and variant H565A as determined by PFE experiments. a** and **c**: Three Cyclic Voltammetry (CV) scans, namely before (black lines), during (blue lines) and after the injection of 1 mM $FeSO_4$ were performed in the presence of CpI-WT (**a**) or the H565A variant (**c**) on the electrode. **b** and **d**: Chronoamperometric PFE experiments were performed at –0.8 V (**b**) and –0.1 V (**b**). After 300 s, 1 mM $FeSO_4$ was injected. After 600 s, the buffer was exchanged to $FeSO_4$-free buffer. The currents were normalized to the respective currents measured at the 300 s timepoint. **a** to **d**: The PFE experiments were done at pH 8, applying a working electrode rotation of 3000 rpm and a temperature of 10 °C. The $H_2$ flow was set to 30 L × $min^{-1}$ (**a**, **c** and **d**) or no $H_2$ was supplied (**b**). The scan rates in **a** and **c** were 10 mV × $s^{-1}$.

**Figure S18. Effect of $NiCl_2$ on the electrochemical activity of CpI-WT and variant H565A.**

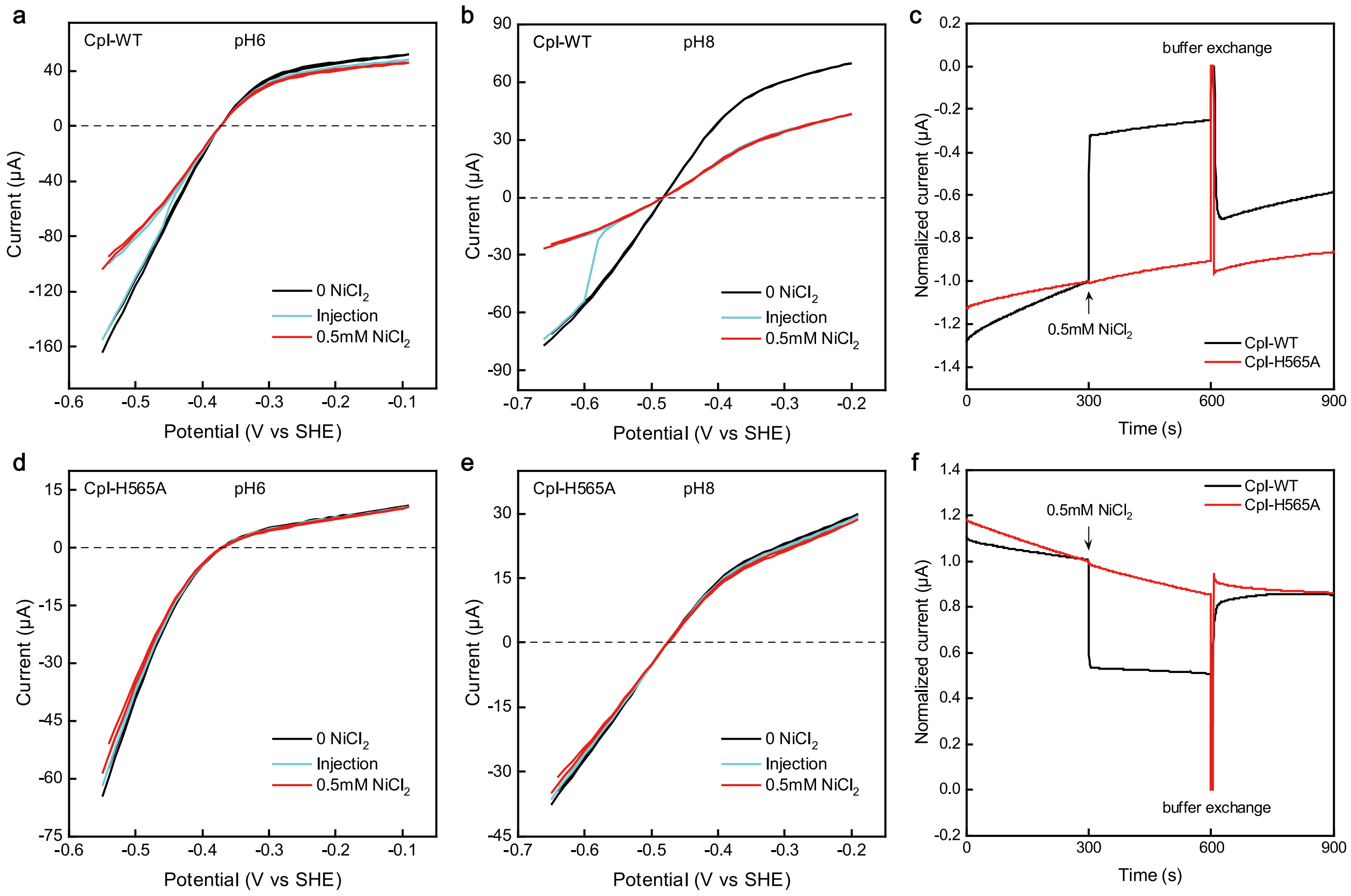


**Figure S18. Effect of $NiCl_2$ on the electrochemical activity of CpI-WT and variant H565A.** Protein film experiments were performed applying the same settings as described in the caption of Figure S17, purging all samples with $H_2$ at a flow rate of 30 L × min$^{-1}$, except for the experiments shown in **c**. **a**, **b**, **d** and **e**: Cyclic voltammetry scans were recorded at a scan rate of 10 mV × s$^{-1}$ before (black lines), during (blue lines) and after (red lines) the addition of 0.5 mM $NiCl_2$ to the buffer within the electrochemical cell, which was at pH 6 (**a**, **d**) or pH 8 (**b**, **e**). **c** and **f**: Chronoamperometric currents were recorded at –0.65 V (**c**) and –0.2 V (**f**). After 300 s, 0.5 mM $NiCl_2$ was injected, and after 600 s, the buffer was exchanged to $NiCl_2$-free buffer. The currents were normalized to their values at 300 s.

**Figure S19. Effect of pH and Cl⁻ ions on the $Zn^{2+}$-mediated inhibition of CpI WT.**

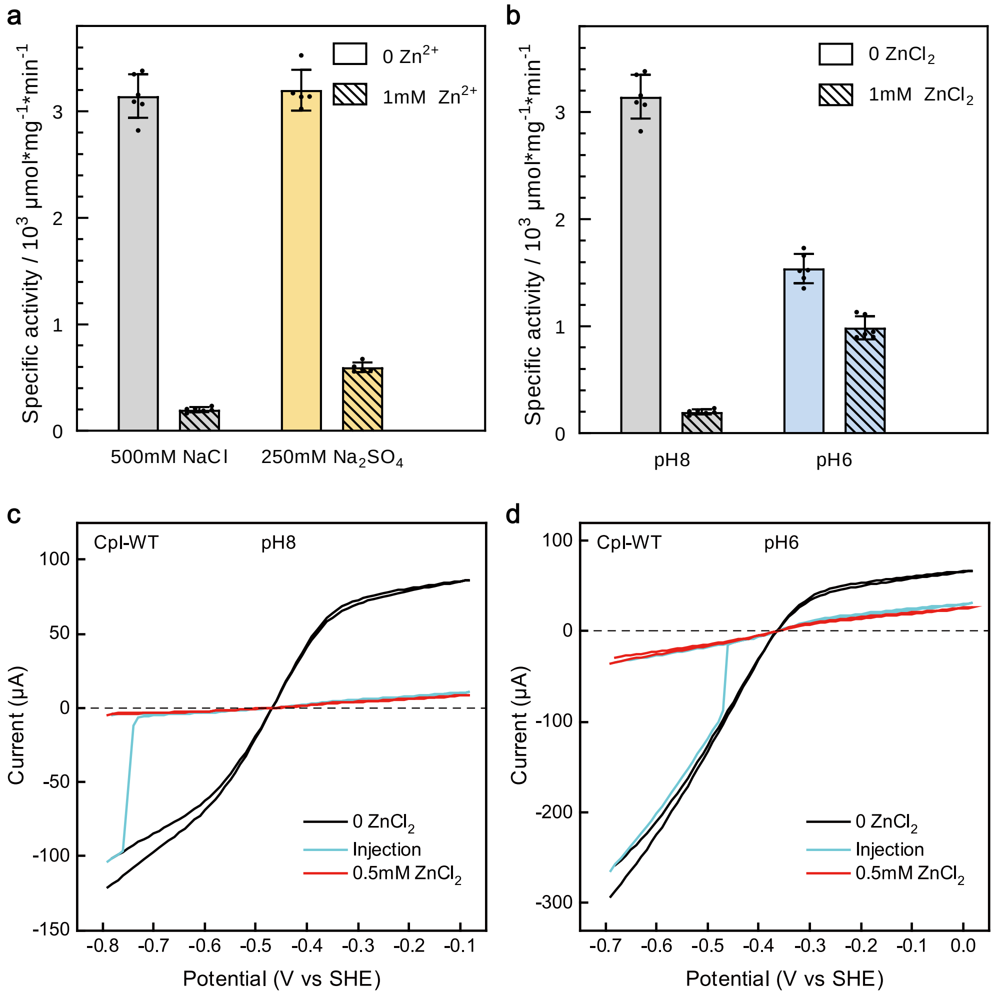


**Figure S19. Effect of pH and Cl⁻ ions on the $Zn^{2+}$-mediated inhibition of CpI-WT.** (**a**) The influence of 1 mM $ZnCl_2$ (striped gray bars) or 1 mM $ZnSO_4$ (striped yellow bars) on $H_2$ production activities of CpI-WT were tested in the presence of different salts, namely 500 mM NaCl (gray bars) that was a standard ingredient of the assays employed here, or 250 mM $Na_2SO_4$ that was added instead of NaCl (yellow bars). (**b**) The effect of 1 mM $ZnCl_2$ (striped bars) on $H_2$ production activities (plain bars) of the CpI-WT enzyme were measured in the dithionite- and methyl viologen-based hydrogenase assay that was adjusted to pH 8 employing 50 mM HEPES buffer (gray bars) or pH 6 using 50 mM MES buffer (blue bars). The hydrogenase assay reaction mixture also contained 500 mM NaCl. (**c**) and (**d**) Three CV scans, namely before (black lines), during (blue lines) and after the injection of 0.5 mM $ZnCl_2$ were recorded at pH 8 (**c**) and pH 6 (**d**) in the presence of the CpI wild type enzyme. The experiments were performed under the following experimental conditions: working electrode rotation at 3000 rpm, temperature of 10 °C, $H_2$ flow rate of 30 L × $min^{-1}$ and scan rate of 10 mV × $s^{-1}$.

**Figure S20. Proton transfer pathways in [FeFe]-hydrogenases CpI, CbA5H, HydA1 and DdH.**

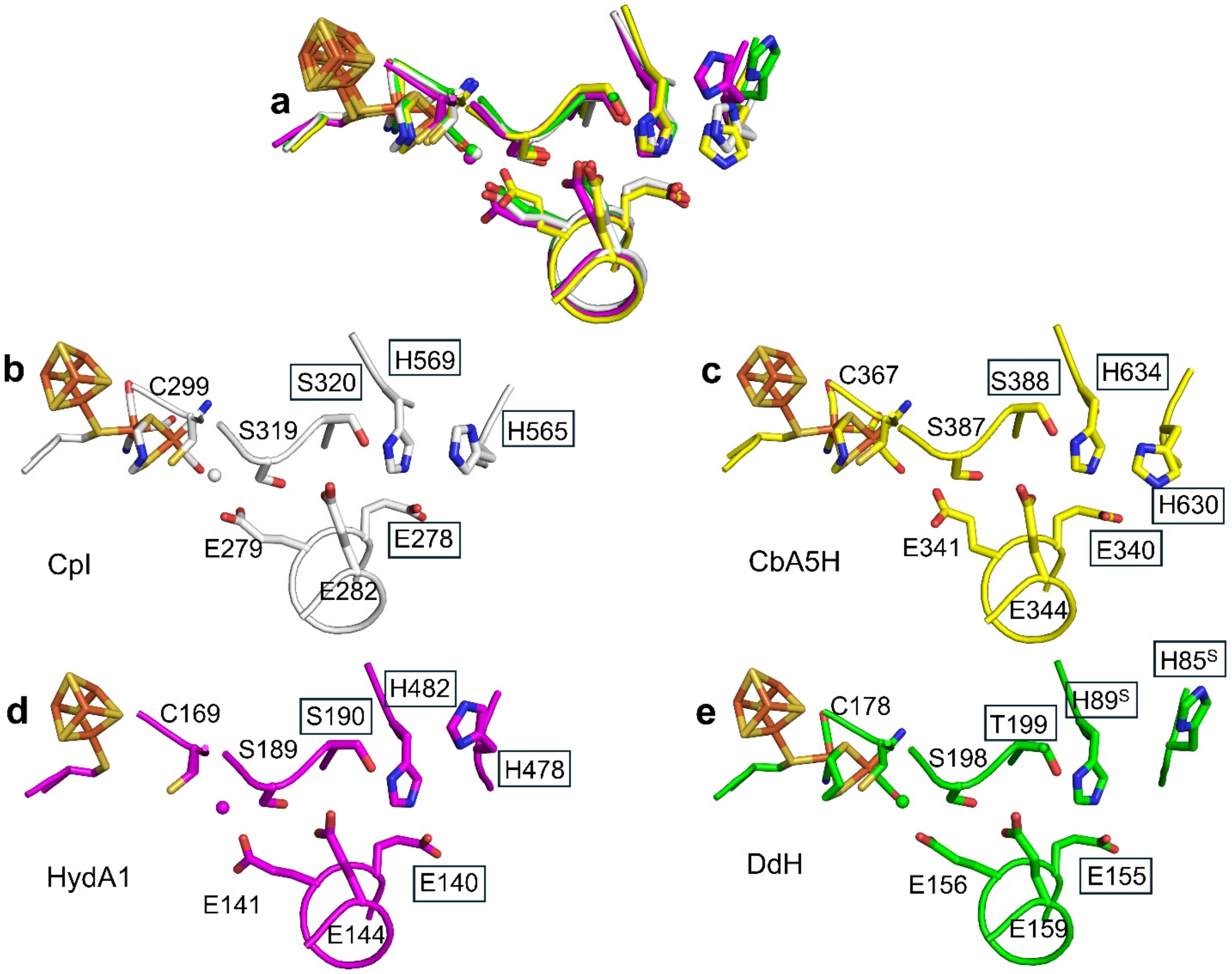


**Figure S20. Proton transfer pathways in [FeFe]-hydrogenases CpI, CbA5H, HydA1 and DdH.** The graphic (**a**) shows an overlay of the structures of the proton transfer pathways (PTP) of CpI, CbA5H, HydA1 and DdH. The PTP of each structure is individually shown for CpI (**b**), CbA5H (**c**), HydA1(**d**) and DdH (**e**). CpI E278, S320, H565 and H569 as well as their counterparts in other [FeFe]-hydrogenases are labeled with rectangles. In panel **d**, the numbering of residues in HydA1 is adapted to be consistent with the numbering used elsewhere in this study and in other studies[3-4]. In panel **e**, the superscripts "S" in H89$^{S}$ and H86$^{S}$ indicate that these two residues belong to the small second subunit of the heterodimeric DdH enzyme[5]. Note that DdH features a conserved substitution of CpI-S320 to a threonine (T199). The PDB accession numbers of the structures employed for this figure are 4XDC (CpI)[1], 8ZQD (CbA5H)[6], 3LX4 (HydA1)[4] and 1HFE (DdH)[5].

**Figure S21. Sequence alignment of CpI, HydA1, CbA5H, ToHydA, TmHydS and TamHydS.**

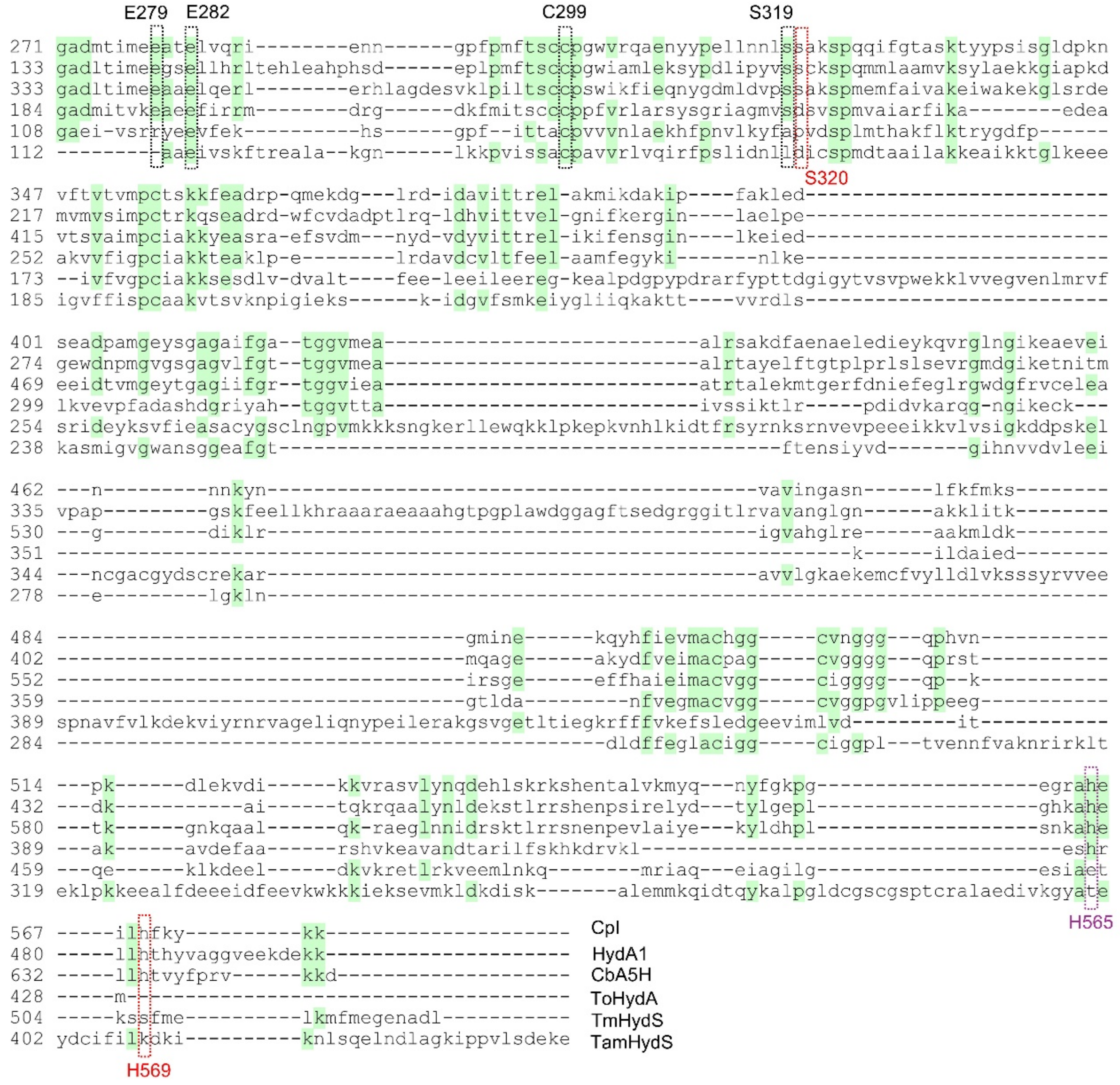

**Figure S21. Sequence alignment of CpI, HydA1, CbA5H, ToHydA, TmHydS and TamHydS.** The residues of the proton transfer pathway are bordered by dashed lines. Residues corresponding to H565 and H569 of CpI are additionally indicated. For clarity, the N-termini of the sequences were omitted. CpI[7], HydA1[4], CbA5H[8] (all group A), ToHydA[9] (group B), TmHydS[10] and TamHydS[11] (group D) represent [FeFe]-hydrogenases from *Clostridium pasteurianum, Chlamydomonas reinhardtii, Clostridium beijerinckii, Thermosediminibacter oceani, Thermotoga maritima and Thermoanaerobacter mathranii*.

**Figure S22. Profiles of $Zn^{2+}$ binding on the surface of HydA1-Δ[2Fe]$_H$.**

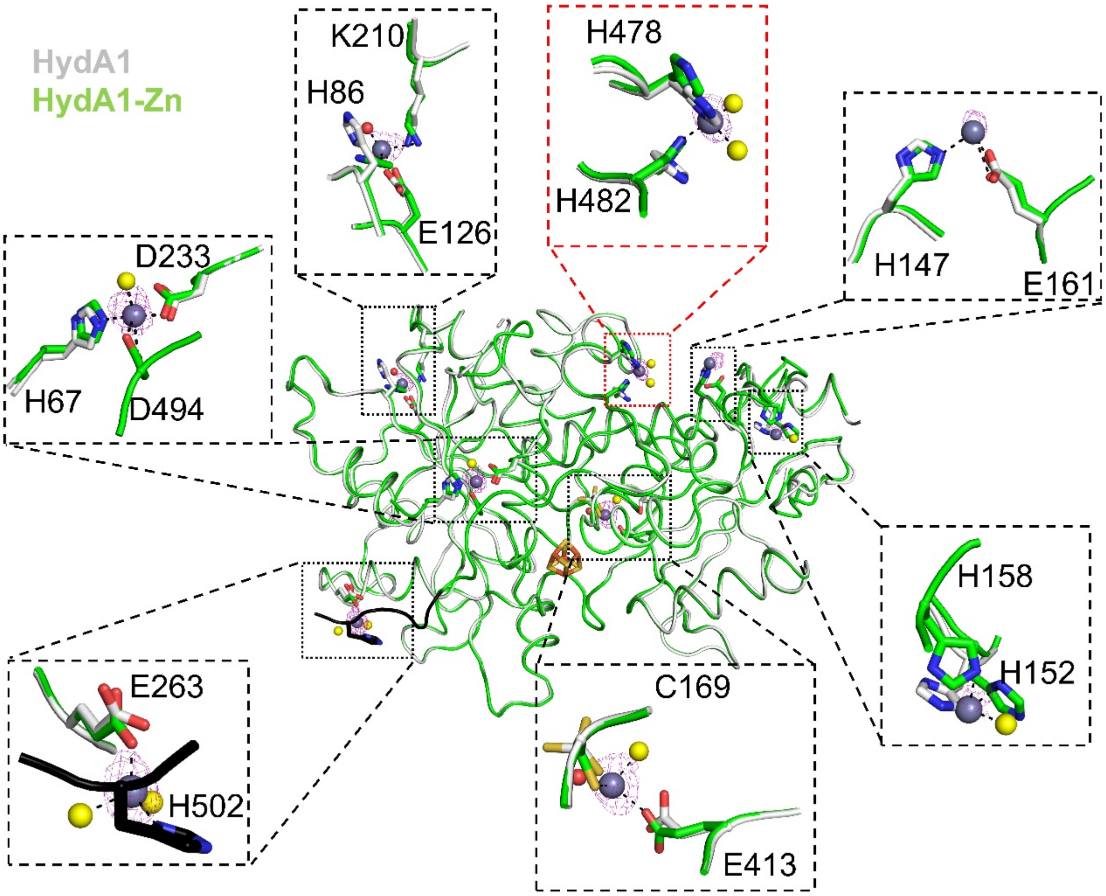


**Figure S22. Profiles of $Zn^{2+}$ binding on the surface of HydA1-Δ[2Fe]$_H$.** The structure of $Zn^{2+}$-soaked crystals of HydA1 without the diiron site of the H-cluster (HydA1-Δ[2Fe]$_H$) (HydA1-Zn; carbon atoms colored in green; PDB: 9SAD) is superimposed on the structure of untreated HydA1-Δ[2Fe]$_H$ published before (carbon atoms in white; PDB ID: 3LX4)[7]. The anomalous map of the identified $Zn^{2+}$ ions (shown as gray spheres) are shown as pink mesh and were contoured at 3 σ. Coordinated water molecules and $Cl^-$ ions are shown as red and yellow spheres, respectively. The red dashed border indicates the $Zn^{2+}$-binding site at the PTP entrance that is zoomed in in Figure S23. A conformational change of the C-terminus (carbon atoms colored black) in HydA1-Zn was observed and is due to a joint coordination of the $Zn^{2+}$ ion by the side chains of E263 and H502.

**Figure S23. $Zn^{2+}$ binding at the entrance of the PTP in HydA1-Δ[2Fe]$_H$.**

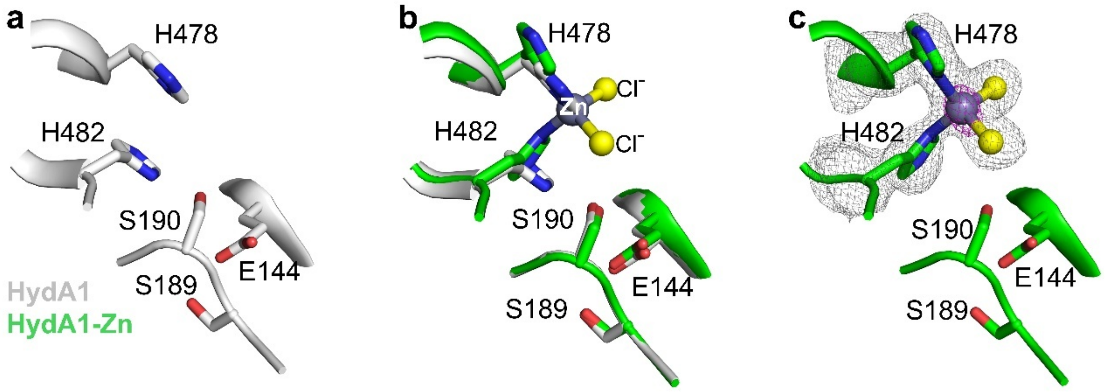


**Figure S23. $Zn^{2+}$ binding at the entrance of the PTP in HydA1-Δ[2Fe]$_H$**. Panel (**a**) shows the HydA1**-**Δ[2Fe]$_H$ structure obtained previously without $Zn^{2+}$ soaking (PDB ID: 3LX4[4]; white carbon atoms). In (**b**), the structure of HydA1-Δ[2Fe]$_H$ obtained after soaking crystals with $Zn^{2+}$ (HydA1-Zn; green carbon atoms; PDB: 9SAD) is superimposed on the structure of untreated HydA1**-**Δ[2Fe]$_H$. In (**c**), the anomalous map of the $Zn^{2+}$ ions is contoured at 4 σ and shown as magenta-colored mesh. The omit map, depicted as gray mesh, was contoured at 2.5 σ.

**Figure S24. Analysis of the structure of CbA5H with a focus on the PTP entrance.**

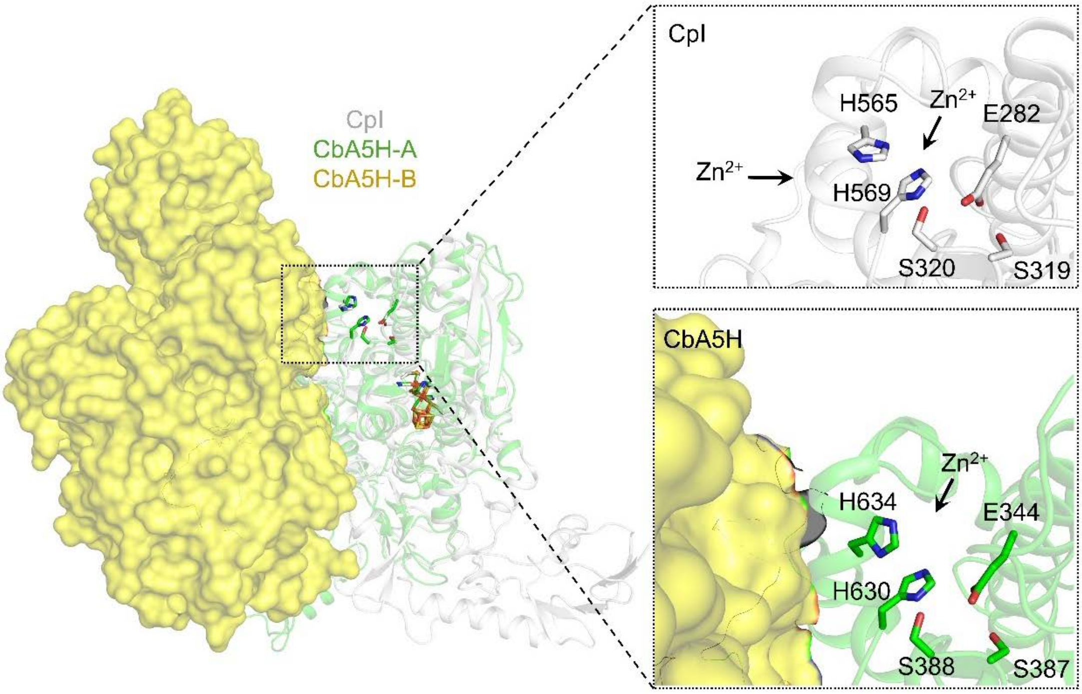


**Figure S24. Analysis of the structure of CbA5H with a focus on the PTP entrance.** The structure of the monomeric CpI enzyme is superimposed on chain A of the homodimeric CbA5H hydrogenase. Carbon atoms of CpI, CbA5H chain A and CbA5H chain B are colored white, green and yellow, respectively. Probable $Zn^{2+}$ entry routes are indicated by the arrows. The presence of the second subunit of CbA5H appears to restrict the accessibility of the entrance of the PTP to the solvent. The figures were created using the structures deposited under the PDB IDs 4XDC (CpI)[1] and 8ZQD (CbA5H)[6].

**Figure S25. Comparison of proton transfer pathways in structures of CpI-WT and CpI variants S320A, H569A, E278Q and H565A.**

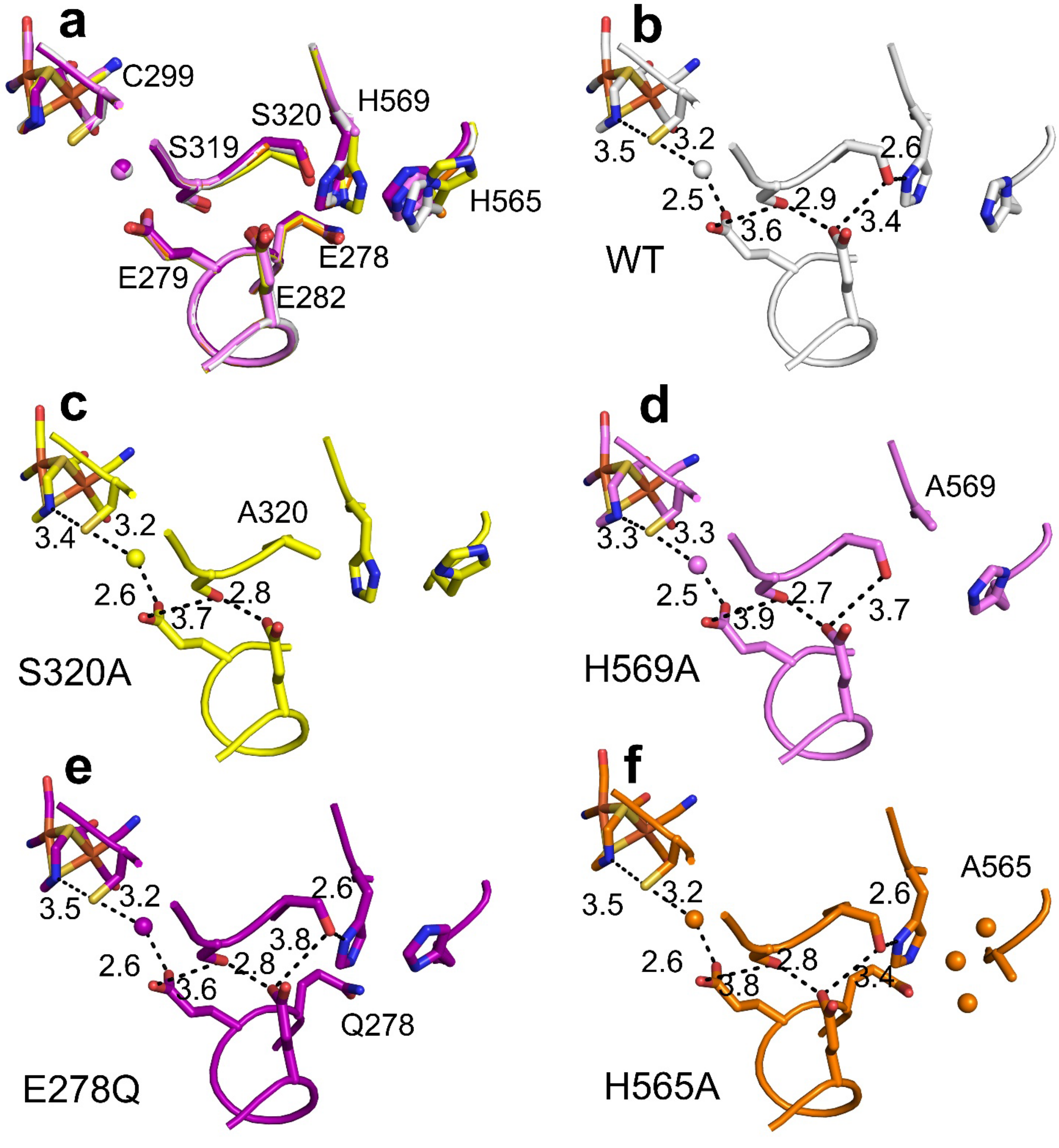


**Figure S25. Comparison of proton transfer pathways in structures of CpI-WT and CpI variants S320A, H569A, E278Q and H565A.** (**a**) shows an overlay of the structures of the proton transfer pathways (PTP) of CpI-WT, S320A, H569A, E278Q and H565A. The PTP of each structure is individually shown for CpI-WT[1](**b**), S320A (**c**), H569A(**d**), E278Q (**e**) and H565A (**f**). Dashed lines represent H-bond interactions between adjacent elements within the PTPs. The numbers indicate H-bond distances in Å.

**Figure S26. Comparison of electrocatalytic properties of CpI-WT, CpI-H569A and S320A at pH 5-9.**

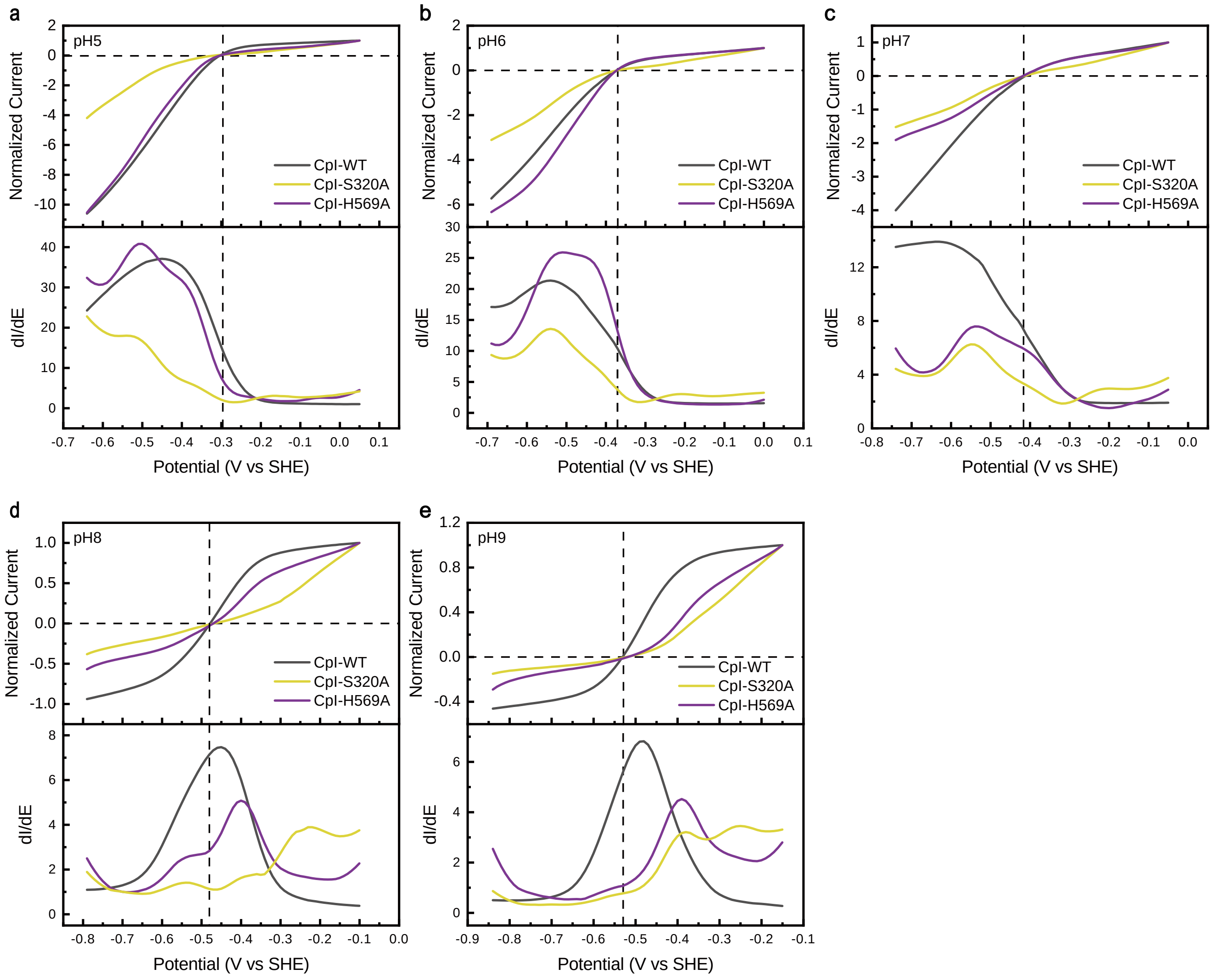


**Figure S26. Comparison of electrocatalytic properties of CpI-WT, CpI-H569A and S320A at pH 5-9.** Comparisons of CV scans of the variants with scans obtained with CpI-WT are shown in each panel: at pH 5 (**a**), pH 6 (**b**), pH 7 (**c**), pH 8 (**d**) and pH 9 (**e**). Underneath each CV scan, a first derivate plot is presented. The CV scans were normalized to the maximum $H_2$ oxidization currents and recorded under the following experimental conditions: working electrode rotation at 3000 rpm, temperature of 10 °C, $H_2$ flow rate of 30 L × min$^{-1}$ and scan rate of 10 mV × s$^{-1}$.

**Figure S27. Comparison of electrocatalytic properties of [FeFe]-hydrogenases HydA1 and CbA5H and variants targeting positions that correspond to CpI-H569 and S320**

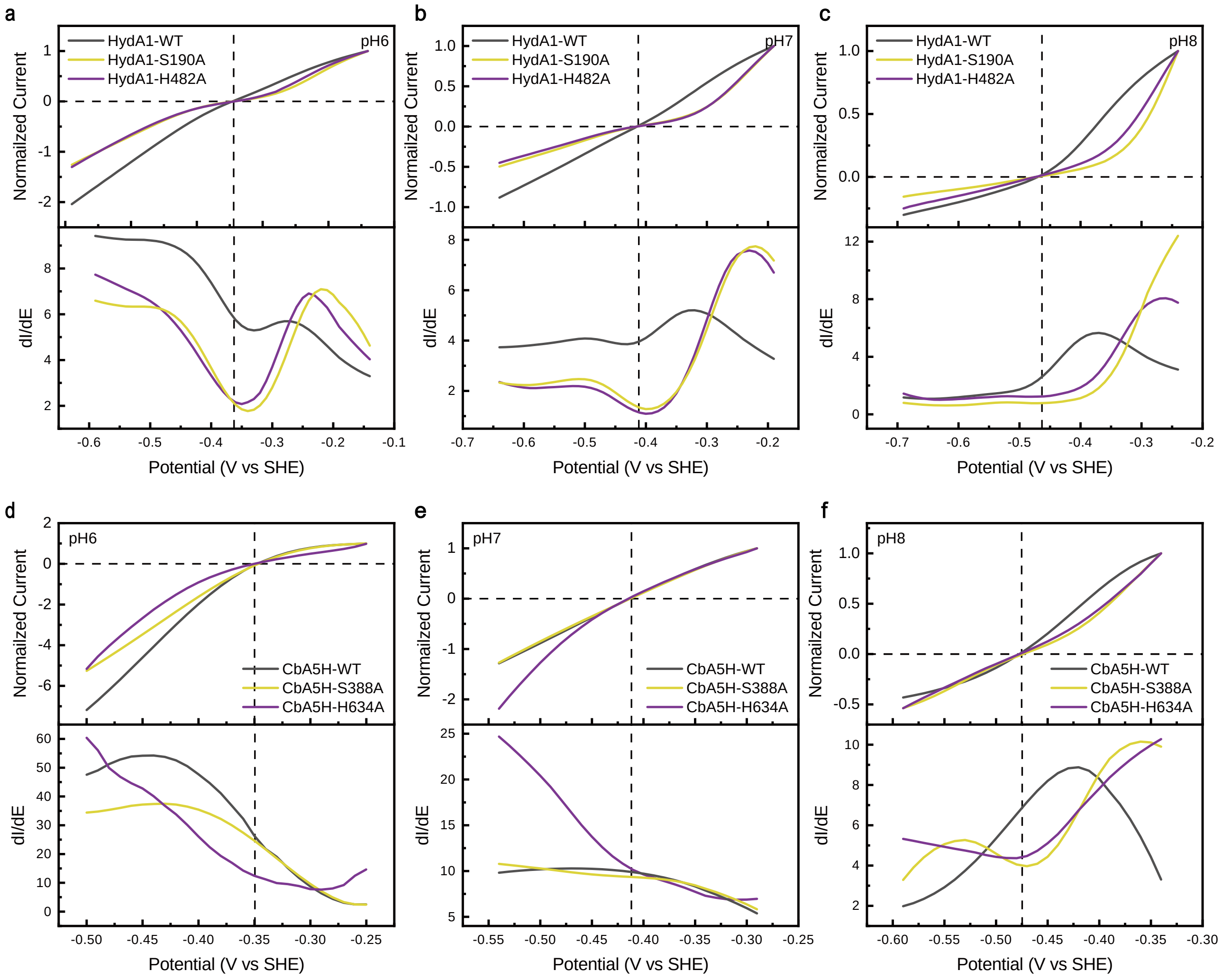


**Figure S27. Comparison of electrocatalytic properties of [FeFe]-hydrogenases HydA1 and CbA5H and variants targeting positions that correspond to CpI-H569 and S320.** Comparisons of normalized CV scans of the variants with scans obtained with the corresponding WT enzyme are shown in each panel: HydA1 at pH 6 (**a**), pH 7 (**b**) and pH 8 (**c**); CbA5H at pH 6 (**d**), pH 7 (**e**) and pH 8 (**f**). Underneath each CV scan, a first derivate of the current against potential was plotted. The CV scans were recorded under the following experimental conditions: working electrode rotation at 3000 rpm, temperature of 10 °C, $H_2$ flow rate of 30 L × min$^{-1}$ and scan rate of 10 mV × s$^{-1}$.

**Figure S28. Comparison of electrocatalytic properties of CpI-WT and variants CpI-H565A, E278Q and H569A.**

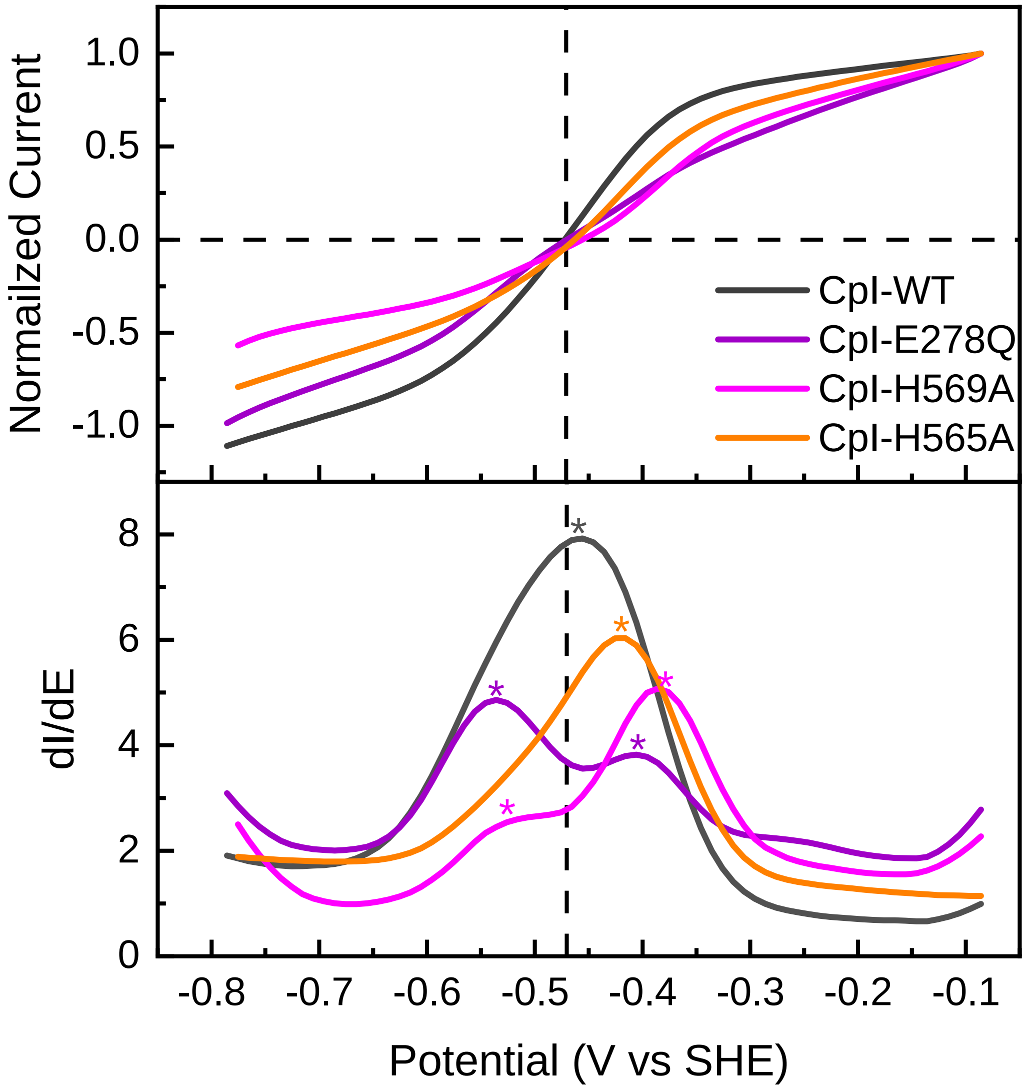


**Figure S28. Comparison of electrocatalytic properties of CpI-WT and variants CpI-H565A, E278Q and H569A.** CV scans were obtained at pH 8 by protein film electrochemistry as described in the caption of Figure S27 and normalized to the maximum $H_2$ oxidation currents. The lower part shows the first derivate of the current against potential. The asterisks indicate the local maximums in the plots.

**Figure S29. Two conformations of H569 in structures of CpI-WT and variant H565A under different pH conditions.**

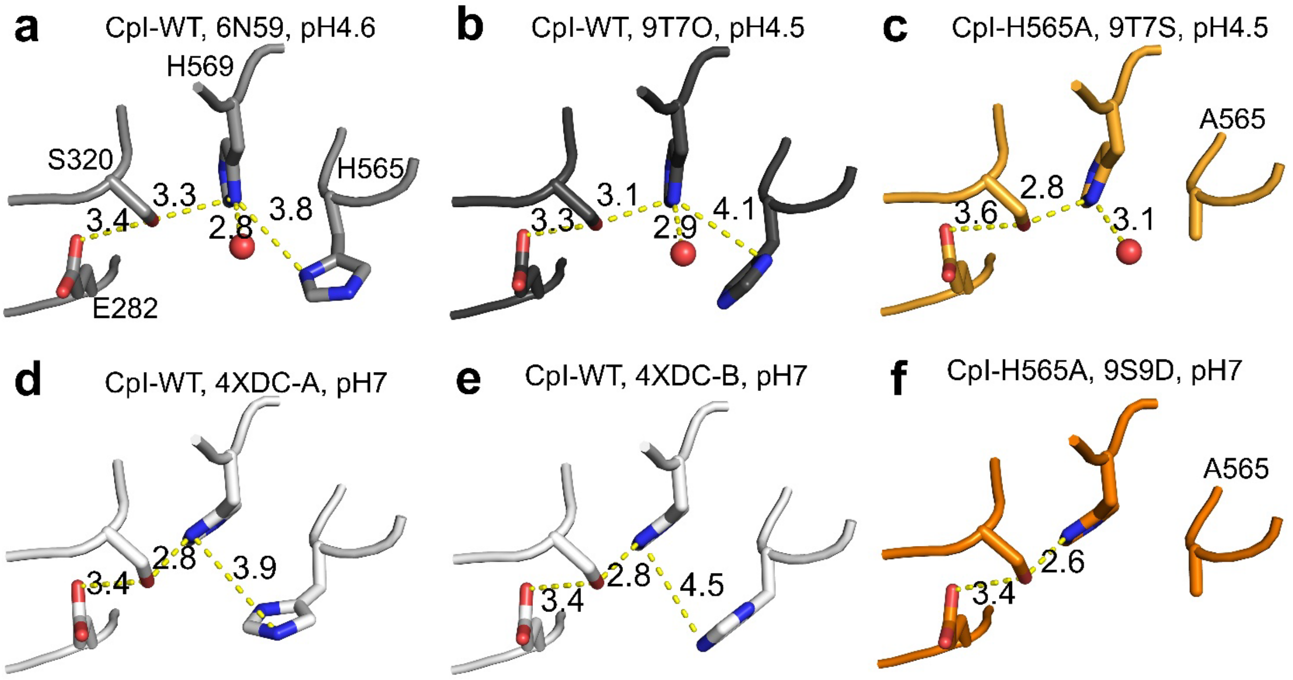


**Figure S29. Two conformations of H569 in structures of CpI-WT and variant H565A under different pH conditions.** H569 exhibits an outward conformation in structures crystallized at pH 4.6 (PDB: 6N59)[12](**a**), crystallized at pH 6 and then soaked in pH 4.5 (this study; chain A of PDB: 9T7O) (**b**) and crystallized at pH 6, but soaked in pH 4.5 of the CpI-H565 variant (this study; chain A of PDB: 9T7S) (**c**), whereas H569 is in an inward conformation in structures of CpI-WT obtained at pH 7 (PDB: 4XDC)[1](**d** and **e**) and of the CpI-H565A variant obtained at pH 7 (this study; PDB: 9S9D) (**f**). Notably, H569 exhibits the outward conformation in chain B of 9T7O and 9T7S. The dashed lines with numbers indicate the H-bonds and respective distances in angstrom.

**Figure S30. DFT-optimized structures of key species along the proton transfer pathway.**

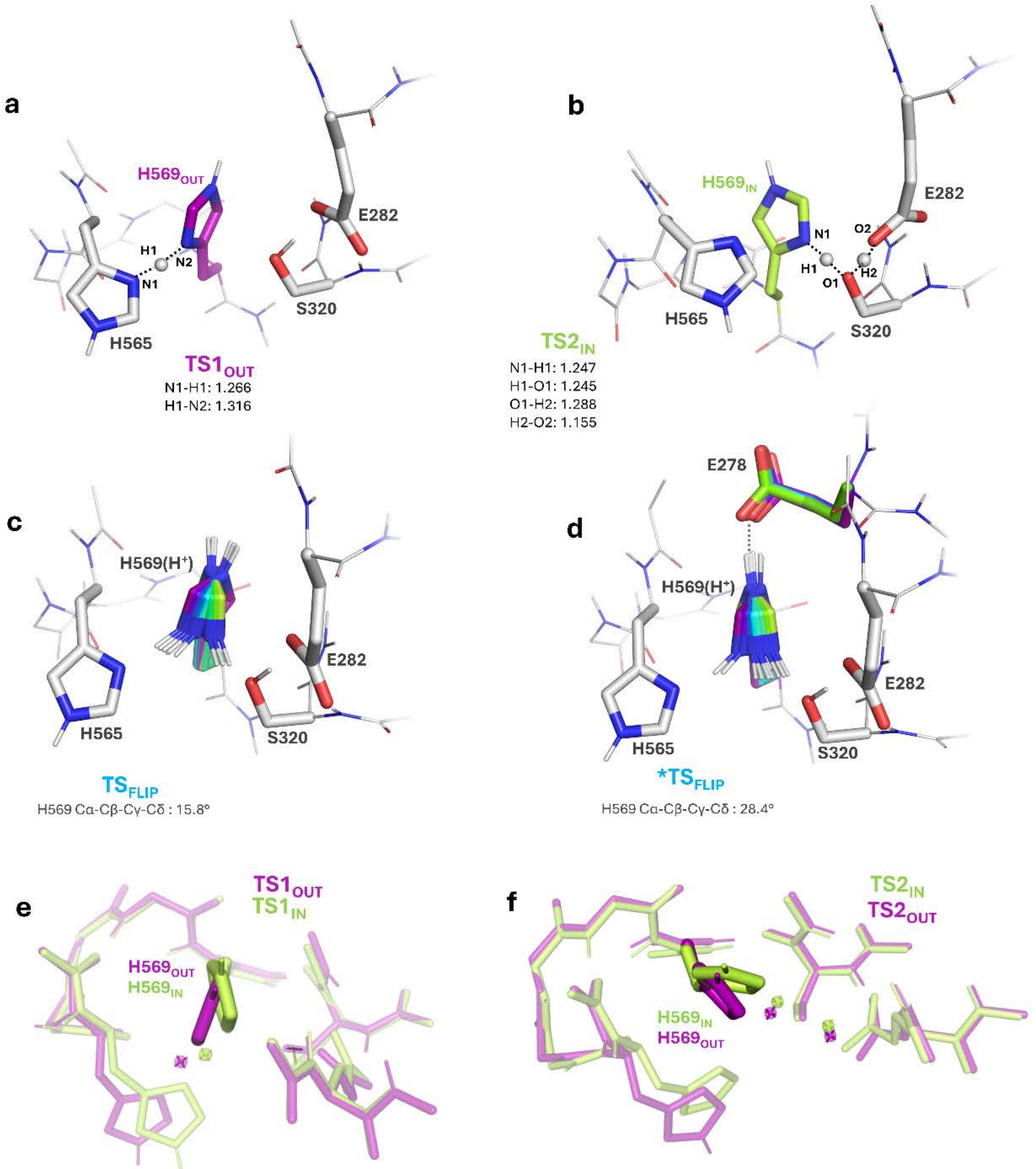


**Figure S30. DFT-optimized structures of key species along the proton transfer pathway.** Selected geometrical parameters are reported (distances are given in Å). (**a**, **b**) Optimized structures of the most stable transition states for the PT steps, TS1$_{OUT}$ (**a**) and TS2$_{IN}$ (**b**). (**c**, **d**) Transition states associated with the conformational flip of protonated H569 obtained from a scan of the Cα–Cβ–Cγ–Cδ dihedral angle, calculated without (**c**) and with (**d**) E278. TS$_{FLIP}$ and *TS$_{FLIP}$ (cyan) are displayed together with representative structures along the scan, colored from purple (OUT) to light green (IN). The comparison highlights the role of E278 as a pivot controlling H569 reorientation. (**e**, **f**) Transition states obtained when starting from opposite H569 orientations: TS1 with H569 initially IN vs OUT (**e**), and TS2 with H569 initially OUT vs IN (**f**). When H569 initially points opposite to the direction of proton transfer, it partially rotates at the transition state (accompanied by a slight rearrangement of the adjacent H565), but its orientation remains suboptimal for proton transfer, indicating that a prior flip of H569 may be required.

**Figure S31. Comparison of the effect of $ZnCl_2$ on $H_2$ production activities of CpI-WT and variant E279A.**

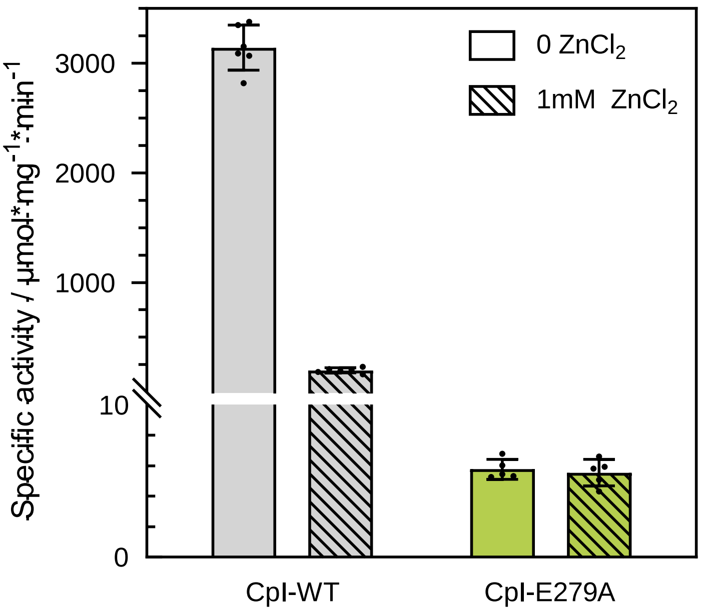


**Figure S31. Comparison of the effect of $ZnCl_2$ on $H_2$ production activities of CpI-WT and variant E279A.** Hydrogenase activity was assayed in the absence (plain bars) or presence (striped bars) of 1 mM $ZnCl_2$ as described in detail in the experimental section at 37 °C in 50 mM HEPES (pH 8) that contained 500 mM NaCl, 100 mM NaDT as reducing reagent and 10 mM methyl viologen as redox mediator. 400 ng of CpI WT or variant E279A were used for each reaction assay.

**Figure S32. Proton transfer pathways (PTP) and surface-exposed histidine residues at the entrances of the PTPs in [FeFe]-hydrogenases CpI and ToHydA.**

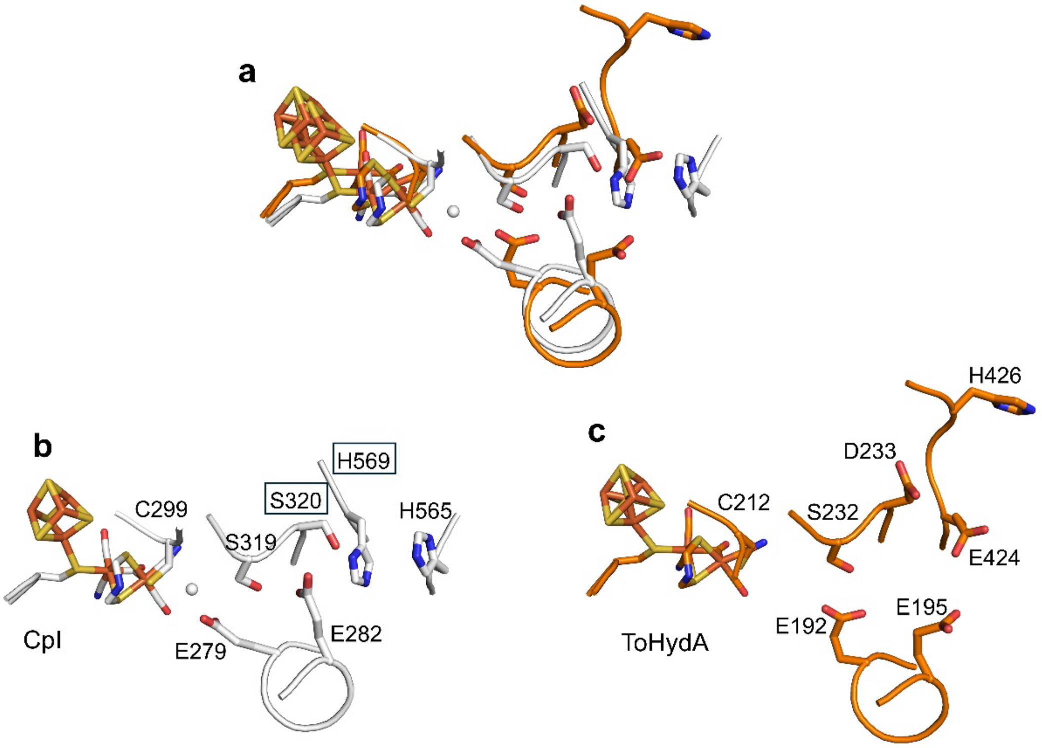


**Figure S32. Proton transfer pathways (PTP) and surface-exposed histidine residues at the entrances of the PTPs in [FeFe]-hydrogenases CpI and ToHydA.** An AlphaFold model of ToHydA[9, 13] was compared to the crystal structure of CpI[1]. Panel **a** shows an overlay of the proton transfer pathways (PTP) of both enzymes. The PTP of each enzyme is individually shown for CpI (**b**) and ToHydA (**c**). The counterparts of CpI-S320 and H569 were identified as D233 and E424, respectively, in the AlphaFold model of ToHydA. No residues that correspond to the position of CpI-H565 could be identified in the respective region of the ToHydA AlphaFold model. Instead, a histidine (H426) located C-terminal of E424 was identified. This analysis suggests either large structural differences between these two enzymes or a significant inaccuracy of the AlphaFold model in this region.

**Figure S33. Protein sequence alignment of exemplary group A [FeFe]-hydrogenases.**

```
274 mtimeeatelvqri--------e--------nn----------------------------------gpfpmftsccpgwvrqaenyype
143 ltiteeaselvqri--------t--------sgk---------------------------------nplpqftsccpawvefveiyype
336 ltimeeaaelqerl--------e--------rhlagdes----------------------------vklpiltsccpswikfieqnygd
273 mtimeeatellgrv--------k--------nn----------------------------------gpfpmftsccpawvrlaqnyhpe
274 mtimeeatelieri--------k--------nn----------------------------------gpfpmltsccpswvrevenyfpe
136 ltimeegsellhrltehleahph--------sd----------------------------------eplpmftsccpgwiamleksypd
 86 ltimeegtellhrl--------kdhlegnpkne----------------------------------eplpmftsccpgwvelveksypd
393 ltimeegsellqrl--------q--------hhwqhpdspahtpphedprfkgagsagapkaghhtpgplpmftsccpawineverdrpe
E278 E279 E282                                                                       C299
314 llnnlssakspqqifgtasktyypsisgldpknvftvtvmpctskkfeadrpqm-----------------------ekdg-------l-
184 linnlssskspilmqgptiktyfakkagidpkkivnvavtpctakkyeitraek-----------------------ndsgkyyndetm-
382 mldvpssakspmemfaivakeiwakekglsrdevtsvaimpciakkyeasraef-----------------------svdm-------n-
313 lldnlssakspqqifgtasktyypsisgiapedvytvtimpcndkkyeadipfm-----------------------etns-------l-
314 lvenlssakspqqifgaasktyypqvadidpkkvftvtvmpctskkfeadrpem-----------------------eneg-------i-
184 lipyvssckspqmmlaamvksylaekkgiapkdmvmvsimpctrkqseadrdwf-----------------------cvdadpt----l-
134 mipylssckspqmmlgaiiknyfadvagyapqdviscsvmpctrkqgeadrpag-----------------------attg-------la
467 liphlstckspqgmmgavvksiwakkmglnpedvvlvsvmpctakkhearreellpeqpqgdaltpadtttsvytstsgdg-------it
S319 S320
373 -----------rdidavittrelakmikdakipfakledseadpamgeysgagaifgatggvmeaalrsakdfa--enaeledieykqvr
250 -----------rdmdyvitvrelsnwikesnidfkalqgaefdsllsrgsgggiifgitggvmesairtayyyitkknppknlynleavg
441 -----------ydvdyvittrelikifensginlkeiedeeidtvmgeytgagiifgrtggvieaatrtalekm--tgerfdniefeglr
372 -----------rdidasltttrelakmikdakikfadledgevdpamgtysgagaifgatggvmeaairsakdfa--enkelenvdytevr
373 -----------rnidavittrelarmikaakidfakledgevdpamgeytgagvifgatggvmeaalrtakdfm--endnldnvdyeavr
246 -----------rqldhvittvelgnifkerginlaelpegewdnpmgvgsgagvlfgttggvmeaalrtayelf--tgtplprlslsevr
194 -----------rdvdhvittaelakifqdrgidlpnlpespldnpigegsgagqlfgttggvmeaavrtvyelv--tgqpmerinltevr
550 ggdrvvcqepardvdfvlttrelgtmlrqmgiplasleagdydhplaggtgaavlfgatggvmeaalrtvaavv--ggaplprlemeair

450 glngikeaevei---nn-------nkyn------------------------------------------vaving---asnlfkfmksg
329 nmdgireaqvti---gd-------ytin------------------------------------------iaivhg---tanarkliekv
518 gwdgfrvcelea---gd-------iklr------------------------------------------igvahg---lreaakmldki
449 gfkgikeaevei---ag-------nkln------------------------------------------vaving---asnffefmksg
450 glagikeaevei---ag-------neyk------------------------------------------lavvsg---aanvfelvksg
323 gmdgiketnitmvpapg-------skfeellkhraaaraeaaahgtpgplawdggagftsedgrggitlrvavang---lgnakklitkm
271 gldgikeatlvl---kpapdsil-gkwsgegeglpir---------------------------------iavang---lgnakklinni
638 gmagvkeatlql---egpglpggakevr------------------------------------------vavasgigaaqqlldkvqsg
                                                                                H565  H569
485 minekqyhfievmachggcvngggqphvnpkdlekvdikkvrasvlynqdeh-lskrkshentalvkmyqnyfgkpgegraheilhfky-
364 ksgekkydfvevmtcrggciggggqpkvkipmadkvrqkriag--lynkdqs-vtqrlahenpdiikvydeffekplsplaeellhtvy-
553 rsgeeffhaieimacvggciggggqp--ktkgnkqaalqk-raeglnnidrs-ktlrrsnenpevlaiyekyldhplsnkahellhtvyf
484 kmnekqyhfievmacpggcingggqphvnaldrenvdyrklrasvlynqdknvlskrkshdnpaiikmydsyfgkpgeglahkllhvky-
485 kind--yhfievmacpggcvngggqphisaedsdkmdirevrasvlynqdkn-lekrkshqnsallkmyenymgkpghgrahellhmky-
403 qageakydfveimacpagcvggggqprstdk-----aitqkrqaalynldek-stlrrshenpsirelydtylgeplghkahellhthyv
321 kdgsakydfvevmacpggcisgggqpr-npdk----qiatkrqqsmytider-mtlrrshdnvfiqqlyakflgkpgshkahdllhtpy-
683 qvn---ydfievmscpggciggggqp--ktkdptai-lkrmdatytidqdst-l--rqshenpdvqqlykdaleapgshdahkllhthy-

573 ---------kk--------   C. pasteurianum -I
450 ---------sskkhilge-   C. pasteurianum -II
639 prv------kkd-------   C. beijerinckii
573 ---------tkdknvskhe   C. acetobutylicum
571 ---------kk--------   C. perfringens
487 aggveekdekk--------   C. reinhardtii -I
404 ---------ipggpakq--   U. belkae-I
763 ---------tdrssevks-   U. belkae-II
```

**Figure S33. Protein sequence alignment of exemplary group A [FeFe]-hydrogenases.** The sources of [FeFe]-hydrogenases are *Clostridium pasteurianum*, *C. beijerinckii*, *C. acetobutylicum*, *C. perfringens*, *Chlamydomonas reinhardtii* and *Uronema belkae*. Roman numbers I and II placed after the organism names represent [FeFe]-hydrogenase isoforms I and II from the organism. All of these enzymes have been characterized regarding their activity before[3, 12, 14-16], see details in Table S7. The residues that putatively form the proton transfer pathway are labelled by dashed frames. Note that the sequence of DdH[5] cannot be included in this alignment because of its heterodimeric structure. Instead, its structurally conserved PTP is presented in Figure S20.

**Figure S34. Structural models used for DFT calculations.**

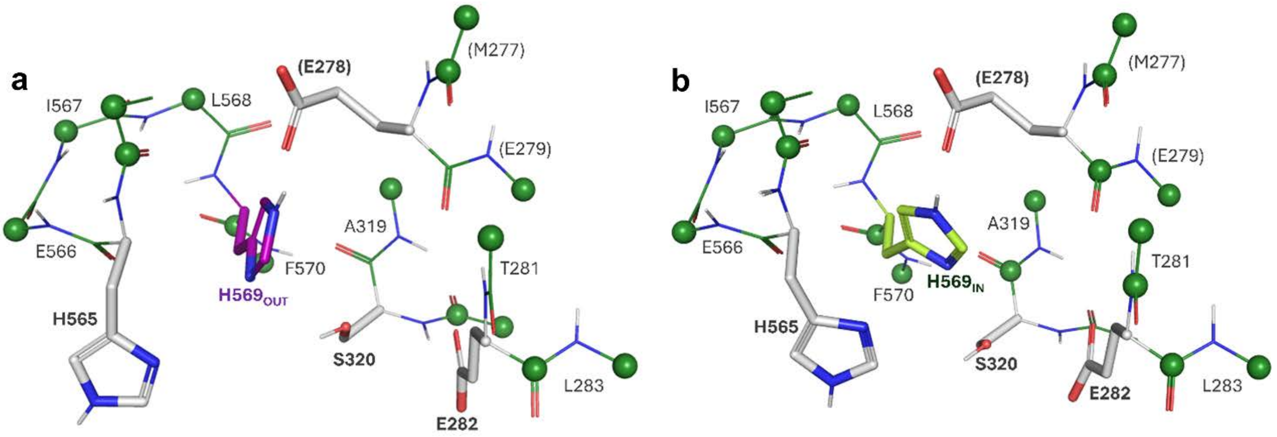


**Figure S34. Structural models used for DFT calculations.** (**a**) model derived from PDB 6N59[12], corresponding to the H569$_{OUT}$ conformation; (**b**) model derived from PDB 4XDC[1], corresponding to the H569$_{IN}$ conformation. Green spheres indicate the carbon atoms whose positions were kept frozen during geometry optimizations. For both models, additional calculations were performed in the absence of E282, in which case the residues listed in parentheses were removed from the cluster.

**Table S1 Crystallization conditions and beamlines used for X-ray diffraction.**

| | Entry | Buffer & pH | Precipitant (PEG4000[b]) | Additives (Glycerol) | Salt | Soaking |
|---|---|---|---|---|---|---|
| CpI-Zn | 9S7W | 0.1 M Tris[a] pH 8 | 17 % | 23 % | 0.4M $MgCl_2$ | 5 mM $ZnCl_2$, ~3 h |
| CpI-Fe | 9S9S | 0.1 M Mes[c] pH 6 | 21 % | 19 % | 0.4 M $MgCl_2$ | 10 mM $FeCl_2$, ~15 h |
| CpI-Ni-1 | 9S8H | 0.1 M Tris pH 8 | 20 % | 20 % | 0.4 M $MgCl_2$ | 10 mM $NiCl_2$, ~15 h |
| CpI-Ni-2 | 9SAG | 0.1 M Mes pH 6 | 17 % | 23 % | 0.4 M NaCl[d] | 10 mM $NiCl_2$, ~15 h |
| CpI-S320A | 9S9W | 0.1 M Tris pH 8 | 20 % | 20 % | 0.4 M $MgCl_2$ | / |
| CpI-H569A | 9S9V | 0.1 M Tris pH 8 | 22 % | 18 % | 0.4 M $MgCl_2$ | / |
| CpI-H565A | 9S9D | 0.1 M Tris pH 8 | 20 % | 20 % | 0.4 M $MgCl_2$ | / |
| CpI-H565A-Zn | 9S9Z | 0.1 M Tris pH 8 | 17 % | 23 % | 0.4 M $MgCl_2$ | 5 mM $ZnCl_2$, ~3 h |
| CpI-H565A-2 | 9T7S | 0.1 M Mes pH 6 | 19 % | 21 % | 0.4 M $MgCl_2$ | pH4.5[e], ~4.5 h |
| CpI-WT-2 | 9T7O | 0.1 M Mes pH 6 | 19 % | 21 % | 0.4 M $MgCl_2$ | pH4.5[e], ~4.5 h |
| CpI-E278Q | 29EI | 0.1 M Tris pH 8 | 19 % | 21 % | 0.4 M $MgCl_2$ | / |
| HydA1-Δ[2Fe]$_H$-Zn | 9SAD | 0.1 M Mes pH 6 | 20 % | / | 0.5 M NaCl | 5 mM $ZnCl_2$, ~3 h |

a "Tris" stands for tris(hydroxymethyl) aminomethane.

b "PEG4000" stands for polyethylene glycol of molecular weight c.a. 4000.

c "Mes" stands for 2-(N-morpholino) ethanesulfonic acid.

d $MgCl_2$ was replaced with NaCl here. Upon the change of salt in the crystallization buffer, changes in the unit cell constants were observed (Table S2). In addition, two conformations of $Ni^{2+}$ coordination at the entrance of PTP were observed in this structure (Figures S5-S6). To exclude a possible link between these two conformations and the salt used for crystallization, we obtained another $Ni^{2+}$-soaked CpI structure without changing crystallization conditions, i.e. using $MgCl_2$ instead of NaCl as salt. The structure showed the same two $Ni^{2+}$ coordination conformations at the PTP entrance as shown in CpI-Ni-2. In the end, CpI-Ni-2 is presented here because of its higher solution.

e The soaking buffer is composed of 0.1 M Mes pH4.5, 19% PEG4000, 21% glycerol and 0.4 M $MgCl_2$.

**Table S2 Root mean square deviations (in Å) of all α-carbon atoms when superimposing the structures with untreated wildtype CpI (a) and HydA1 (b).**

| **a** | 4XDC (A) | 4XDC (B) |
|---|---|---|
| CpI-Zn | 0.378 | 0.379 |
| CpI-Fe | 0.749 | 0.512 |
| CpI-Ni-1 | 0.544 | 0.344 |
| CpI-Ni-2 | 0.491 | 0.524 |
| CpI-S320A | 0.694 | 0.376 |
| CpI-H569A | 0.784 | 0.525 |
| CpI-H565A | 0.728 | 0.352 |
| CpI-H565A-Zn | 0.388 | 0.735 |
| CpI-H565A-2 | 0.606 | 0.556 |
| CpI-WT-2 | 0.528 | 0.394 |
| CpI-E278Q | 0.332 | 0.286 |

| **b** | 3LX4 (A) | 3LX4 (B) |
|---|---|---|
| HydA1-Δ$[2Fe]_H$-Zn | 0.437 | 0.424 |

Notes:

1. A or B represent the respective chains in the asymmetric unit.
2. The number of α-carbon atoms involved in calculating the RMSDs were 565-577 and 405, respectively for CpI and HydA1.

**Table S3 X-ray data collection and refinement statistics of crystal structures CpI-Zn and CpI-Ni-2**

| | Data collection | | | | | |
|---|---|---|---|---|---|---|
| | CpI-Zn | | | | CpI-Ni-2 | |
| PDB | 9S7W | | | | 9SAG | |
| Beamline | DESY-P13 | DESY-P13 | DESY-P13 | DESY-P13 | DESY-P13 | DESY-P13 |
| X-ray energy (keV) | 12.7 | 9.665 | 9.63 | 6.2 | 12.69 | 8.35 |
| Space group | P 1 $2_1$ 1 | | | | P 1 $2_1$ 1 | |
| a, b, c (Å) | 90.81, 73.71, 103.57 | 90.87, 73.80, 103.75 | 90.86, 73.82, 103.73 | 90.68, 73.55, 103.41 | 67.55, 106.33, 86.41 | 67.42, 106.23, 86.32 |
| α, β, γ (°) | 90.00, 98.314, 90.00 | 90.00, 98.409, 90.00 | 90.00, 98.353, 90.00 | 90.00, 98.512, 90.00 | 90.00, 96.548, 90.00 | 90.00, 96.472, 90.00 |
| Resolution (Å) | 43.53-1.65 (1.709-1.65) | 42.13-1.90 (1.968-1.90) | 44.95-1.68 (1.74-1.68) | 47.55-2.23 (2.311-2.23) | 49.593-1.45 (1.502-1.45) | 45.16-1.602 (1.659-1.602) |
| $R_{merge}$ | 0.1138 (2.078) | 0.1080 (1.223) | 0.06486 (1.000) | 0.0677 (0.3792) | 0.1554 (2.073) | 0.09979 (0.7580) |
| I / σ(I) | 9.18 (1.10) | 7.28 (0.98) | 9.45 (1.00) | 15.55 (2.6) | 8.84 (1.10) | 7.79 (1.08) |
| Completeness (%) | 99.87 (99.94) | 98.63 (97.80) | 97.51 (91.34) | 96.46 (69.85) | 99.11 (99.25) | 91.19 (45.47) |
| Multiplicity | 6.9 (6.8) | 3.5 (3.2) | 3.4 (2.8) | 6.3 (4.0) | 7.0 (7.1) | 3.3 (2.1) |
| CC1/2 | 0.998 (0.415) | 0.997 (0.366) | 0.998 (0.387) | 0.999 (0.936) | 0.998 (0.144) | 0.997 (0.181) |
| **Refinement** | | | | | | |
| Resolution (Å) | 43.53-1.65 | | | | 33.55-1.45 | |
| No. reflections | 162372 | | | | 212072 | |
| $R_{work}$ / $R_{free}$ | 0.1691 / 0.2149 | | | | 0.1789 / 0.2164 | |
| **No. atoms** | | | | | | |
| Protein | 8961 | | | | 9011 | |
| Ligand / ion | 106 / 46 | | | | 106 / 55 | |
| Water | 580 | | | | 736 | |
| **B-factors** | | | | | | |
| Protein | 40.42 | | | | 27.30 | |
| Ligand /ion | 34.08 | | | | 23.42 | |
| Water | 41.17 | | | | 36.63 | |
| **R.m.s deviations** | | | | | | |
| Bond lengths (Å) | 0.012 | | | | 0.009 | |
| Bond angles (°) | 1.06 | | | | 0.97 | |

*Numbers in brackets indicate values in the highest resolution shell

**Table S4 X-ray data collection and refinement statistics of crystal structures CpI-Ni-1, CpI-Fe, CpI-H565A and CpI-H565A-Zn**

| | **Data collection** | | | | | |
|---|---|---|---|---|---|---|
| | **CpI-Ni-1** | | **CpI-Fe** | | **CpI-H565A** | **CpI-H565A-Zn** |
| PDB | 9S8H | | 9S9S | | 9S9D | 9S9Z |
| Beamline | DESY-P14 | DESY-P14 | DESY-P13 | DESY-P13 | ESRF ID30B | DESY-P13 |
| X-ray energy (keV) | 14.2 | 8.35 | 12.7 | 7.145 | 14.2 | 11.8 |
| Space group | P 1 $2_1$ 1 | | P 1 $2_1$ 1 | | P 1 $2_1$ 1 | P 1 $2_1$ 1 |
| a, b, c (Å) | 90.09, 72.58, 102.93 | 90.39 73.04 103.15 | 87.50, 71.76, 102.95 | 87.54, 72.19, 102.95 | 89.87, 72.30, 102.68 | 91.53, 73.04, 103.39 |
| α, β, γ (°) | 90.00, 97.037, 90.00 | 90.00, 96.942, 90.00 | 90.00, 101.731, 90.00 | 90.00, 101.245, 90.00 | 90.00, 97.467, 90.00 | 90.00, 98.363, 90.00 |
| Resolution (Å) | 47.81-1.5 (1.554-1.5) | 48.04-1.76 (1.823-1.76) | 47.91-1.50 (1.554-1.50) | 47.60-1.94 (2.009-1.94) | 47.55-1.34 (1.388-1.34) | 47.60-1.45 (1.502-1.45) |
| $R_{merge}$ | 0.1606 (1.615) | 0.1278 (0.4664) | 0.1467 (0.8393) | 0.179 (0.6405) | 0.1005 (2.171) | 0.0644 (1.477) |
| I / σ(I) | 8.23 (1.30) | 6.72 (2.08) | 8.06 (2.05) | 5.01 (1.46) | 10.71 (0.99) | 9.41 (0.96) |
| Completeness (%) | 99.57 (99.53) | 96.30 (84.32) | 98.90 (99.68) | 97.41 (85.69) | 98.94 (99.00) | 98.50 (98.65) |
| Multiplicity | 6.9 (6.9) | 3.4 (3.0) | 6.8 (7.0) | 3.4 (2.8) | 7.0 (7.1) | 3.5 (3.3) |
| CC1/2 | 0.998 (0.190) | 0.989 (0.102) | 0.998 (0.200) | 0.984 (0.125) | 0.999 (0.228) | 0.999 (0.337) |
| **Refinement** | | | | | | |
| Resolution (Å) | 39.23-1.50 | | 47.91-1.50 | | 33.94-1.34 | 37.58-1.45 |
| No. reflections | 209581 | | 197262 | | 288830 | 462786 |
| $R_{work}$ / $R_{free}$ | 0.1821 / 0.2275 | | 0.1859 / 0.2325 | | 0.1547 / 0.1847 | 0.1687 / 0.2012 |
| **No. atoms** | | | | | | |
| Protein | 8988 | | 8943 | | 9064 | 9081 |
| Ligand / ion | 106 / 43 | | 106 / 20 | | 106 / 20 | 106 / 43 |
| Water | 997 | | 838 | | 932 | 804 |
| **B-factors** | | | | | | |
| Protein | 28.23 | | 25.31 | | 26.78 | 35.32 |
| Ligand /ion | 22.61 | | 16.86 | | 18.90 | 29.41 |
| Water | 37.02 | | 35.20 | | 37.90 | 41.71 |
| **R.m.s deviations** | | | | | | |
| Bond lengths (Å) | 0.008 | | 0.008 | | 0.007 | 0.007 |
| Bond angles (°) | 0.87 | | 0.95 | | 0.86 | 0.84 |

*Numbers in brackets indicate values in the highest resolution shell

**Table S5 X-ray data collection and refinement statistics of crystal structures CpI-S320A, CpI-H569A, CpI-E278Q, CpI-WT-2 and HydA1-Δ[2Fe]$_H$-Zn**

| | CpI-S320A | CpI-H569A | CpI-E278Q | CpI-WT-2 | CpI-H565A-2 | HydA1-Δ[2Fe]$_H$-Zn |
|---|---|---|---|---|---|---|
| **Data collection** | | | | | | |
| PDB | 9S9W | 9S9V | 29EI | 9T7O | 9T7S | 9SAD |
| Beamline | ESRF ID30B | ESRF ID30A-3 | DESY-P13 | DESY-P13 | DESY-P13 | ESRF ID30B |
| X-ray energy (keV) | 14.2 | 12.8 | 12.7 | 11.7 | 11.7 | 9.665 |
| Space group | P 1 $2_1$ 1 | P 1 $2_1$ 1 | P 1 $2_1$ 1 | P 1 $2_1$ 1 | P 1 $2_1$ 1 | P $3_2$ 2 1 |
| a, b, c (Å) | 89.83, 72.24, 103.07 | 89.51, 71.94, 103.01 | 90.50, 72.83, 103.40 | 87.58, 73.25, 103.42 | 87.76, 72.35, 103.35 | 70.53, 70.53, 153.72 |
| α, β, γ (°) | 90.00, 97.619, 90.00 | 90.00, 97.31, 90.00 | 90.00, 97.33, 90.00 | 90.00, 101.32, 90.00 | 90.00, 101.25, 90.00 | 90.00, 90.00, 120.00 |
| Resolution (Å) | 47.53-1.35 (1.398-1.35) | 46.94-1.62 (1.678-1.62) | 47.19-1.31 (1.357-1.31) | 37.04-1.34 (1.388-1.34) | 43.04-1.65 (1.709-1.65) | 47.82-2.00 (2.072-2.00) |
| $R_{merge}$ | 0.131 (1.897) | 0.2256 (2.074) | 0.1332 (1.384) | 0.1786 (2.128) | 0.2137 (2.052) | 0.3047 (2.786) |
| I / σ(I) | 7.45 (1.03) | 6.18 (1.02) | 6.67 (1.25) | 6.90 (1.02) | 6.16 (0.79) | 6.26 (0.94) |
| Completeness (%) | 99.92 (99.96) | 99.23 (99.87) | 90.63 (54.30) | 94.75 (92.60) | 99.38 (99.31) | 99.98 (100.00) |
| Multiplicity | 6.8 (6.8) | 7.1 (7.2) | 6.8 (5.5) | 7.1 (7.2) | 6.9 (7.0) | 10.5 (10.7) |
| CC1/2 | 0.997 (0.326) | 0.996 (0.119) | 0.997 (0.0981) | 0.998 (0.172) | 0.994 (0.388) | 0.994 (0.273) |
| **Refinement** | | | | | | |
| Resolution (Å) | 47.53-1.35 | 46.94-1.62 | 47.19-1.31 | 37.04-1.34 | 43.04-1.65 | 47.82-2.00 |
| No. reflections | 285951 | 163559 | 289245 | 272004 | 151649 | 57812 |
| $R_{work}$ / $R_{free}$ | 0.1525 / 0.1817 | 0.2077 / 0.2372 | 0.1898 / 0.2093 | 0.1745 / 0.2086 | 0.1722 / 0.2253 | 0.2065 / 0.2484 |
| **No. atoms** | | | | | | |
| Protein | 9098 | 8990 | 9096 | 9042 | 8928 | 3266 |
| Ligand / ion | 106 / 23 | 106 / 19 | 106 / 38 | 106 / 24 | 106 / 8 | 8 / 16 |
| Water | 1063 | 564 | 977 | 1013 | 850 | 293 |
| **B. factors** | | | | | | |
| Protein | 25.17 | 28.82 | 27.18 | 23.88 | 30.97 | 36.31 |
| Ligand /ion | 18.02 | 19.32 | 18.93 | 15.46 | 22.39 | 37.20 |
| **R.m.s deviations** | | | | | | |
| Water | 35.46 | 33.84 | 32.93 | 34.61 | 40.34 | 38.04 |
| Bond lengths (Å) | 0.007 | 0.01 | 0.011 | 0.008 | 0.007 | 0.002 |
| Bond angles (°) | 0.90 | 1.04 | 1.09 | 0.90 | 0.77 | 0.46 |

*Numbers in brackets indicate values in the highest resolution shell

**Table S6. Oligonucleotides employed to generate variants by QuikChange PCR**

| | Forward 5'-3' | Reverse 5'-3' |
|---|---|---|
| **CpI-C39S** | CTGAATAATGCTAATAATGACATC AATAAGTGTGAAATCTGTAC | GTCATTATTAGCATTATTCAGAAA ACACAGTGCGGAG |
| **CpI-C184A** | GAAAAAGCCTTCGATGACACCA ATTGTCTGCTG | ATCGAAGGCTTTTTCATCCTCTG CGCCAATG |
| **CpI-E278Q** | ATTATGCAAGAGGCTACCGAACT GGTTC | CGGTAGCCTCTTGCATAATGGTC ATATCTG |
| **CpI-S320A** | GAATAATCTTTCCGCCGCTAAAT CCCCTCAACAG | TAGCGGCGGAAAGATTATTCAGC AGTTCAGG |
| **CpI-E444A** | GATATCGCATATAAGCAAGTTCG CGGCCTG | CTTATATGCGATATCTTCAAGTTC AGCGTTTTCAG |
| **CpI-H500A** | GCTTGTGCTGGCGGCTGTGTAA ATGGTGGT | GCCGCCAGCACAAGCCATTACT TCGATGAAATG |
| **CpI-H511A** | CCAGCCTGCTGTAAACCCAAAA GACCTGG | GTTTACAGCAGGCTGGCCACCA CCATTTAC |
| **CpI-E535A** | CAGGATGCACATCTTTCCAAGC GCAAATC | AAGATGTGCATCCTGATTATACA GTACAGAAGC |
| **CpI-H565A** | CGTGCCGCTGAAATCCTGCACT TTAAATAT | GATTTCAGCGGCACGACCTTCA CCTGG |
| **CpI-H569A** | ATCCTGGCCTTTAAATATAAAAAA TCAGCCTGGTCC | TTTAAAGGCCAGGATTTCATGGG CACGACC |
| **HydA1-S190A** | GTGAGCGCGTGCAAAAGCCCG CAGATG | TTTGCACGCGCTCACATACGGA ATCAGATCC |
| **HydA1-H482A** | GAACTGCTGGCGACCCATTATGT GGCG | ATGGGTCGCCAGCAGTTCATGC GCTTTATG |
| **CbA5H-S388A** | CCGAGCGCCGCAAAAAGCCCG ATG | TTTTGCGGCGCTCGGAACATCC AGC |
| **CbA5H-H634A** | CTGCTGGCAACCGTTTATTTTCC GCGT | AACGGTTGCCAGCAGTTCATGT GCTTTATT |

**Table S7. Catalytic activities of different [FeFe]-hydrogenases**

| [FeFe]-hydrogenases | $H_2$ production activities | $H_2$ oxidation activities |
|---|---|---|
| *Clostridium pasteurianum* -I | 2576 ± 107 µmol $H_2$ /(mg* min)[3] | 1.8 x $10^4$ µmol MB[b] /(mg* min)[12] |
| *Clostridium pasteurianum* -II[a] | 16.2 ± 0.14 µmol $H_2$ /(mg* min)[12] | (1.1± 0.1) x $10^5$ µmol MB /(mg* min)[12] |
| *Clostridium beijerinckii* | 3479.95± 351.26 µmol $H_2$ /(mg* min)[17] | |
| *Clostridium acetobutylicum* | 1750 µmol $H_2$ /(mg* min)[15] | |
| *Clostridium perfringens* | 1645 ± 16 $s^{-1}$ [14] | |
| *Chlamydomonas reinhardtii* -I | 862 ± 46.5 µmol $H_2$ /(mg* min)[3] | |
| *Uronema belkae* -I | 990 ± 35 µmol $H_2$ /(mg* min)[16] | |
| *Uronema belkae* -II | 2097 ± 261 µmol $H_2$ /(mg* min)[16] | |

Additional notes:

a. CpII is strongly biased to $H_2$ oxidation.
b. MB stands methylene blue in the reduced form.
